# A New Class of Precision Therapeutics that Inhibit Prostate Cancer Mediated Bone Destruction

**DOI:** 10.1101/2025.05.20.654990

**Authors:** Fangfang Qiao, Ryan Gordon, Abhinandan Pattanayak, Wenqi Li, Jaclyn Knibbe-Hollinger, Katelyn O’Neill, Weining Chen, Prathap Reddy Mukthapuram, Amarnath Natarajan, Raymond Bergan

## Abstract

Cancer mediated bone destruction remains a common cause of morbidity and mortality. Here we couple an inhibitor of cell movement to bone trophic bisphosphonates, forming heterobifunctional compounds termed Dual Acting Bone Defender agents (DABDs). DABDs inhibit prostate cancer cell motility and osteoclast mediated bone destruction *in vitro*. Our lead compound, BTE-EN1, inhibits circulating human prostate cancer cells from forming bone destructive lesions in mice and prolongs life in a dose-dependent fashion. In mice with established metastatic lesions, BTE-EN1 significantly decreases bone destruction by over 60% compared to the control. Mechanistically, BTE-EN1 binds hydroxyapatite, inhibits prenylation of Rap1A, induces apoptosis in osteoclasts, and inhibits Raf1 activation and formation of bone resorptive cavities in osteoclasts, thus retaining the function of its constituent chemicals. BTE-EN1 is not toxic to mice at doses 4,000-fold over those required for efficacy, exhibits anti-cell motility activity across other bone-destructive cancer cell types, and does not interfere with standard-of-care hormone therapy or chemotherapy. BTE-EN1 is a new class of highly active therapeutics that inhibit cancer mediated bone destruction and holds high potential for translation into humans.

## INTRODUCTION

Prostate cancer (PCa) is the most common cause of cancer in US men, and the second most common cause of cancer death. Death results from the formation of metastasis. Bone metastases are present in over 90% of those with metastasis, and they dominate the clinical management of such patients (*1*). Bone metastases destroy bone architecture, causing pain, fracture, hypercalcemia, replacement of bone marrow, and nerve and spinal cord compression, leading to paralysis and bowel and bladder incontinence (*2, 3*). Therapeutics that mitigate bone destruction include bisphosphonates, inhibitors of receptor activator of nuclear factor kappa-Β ligand (RANKL), and bone trophic radioisotopes (*4–6*). They have been in existence for several decades, are not considered to have high efficacy, and each has a significant toxicity profile that limits use.

Normal bone is a dynamic heterogenic organ. Three main cell types coordinately regulate continuous remodeling: osteoblasts, osteocytes and osteoclasts (*7–9*). Osteoblasts, generate new bone matrix, then terminally differentiate into osteocytes which become embedded in bone, wherein they serve as a source of cytokines that regulate cell growth (*10–12*). Osteoblasts and osteocytes secrete factors that simulate the maturation and activity of osteoclasts (*7*). Osteoclasts are multinucleated cells that form actin-ringed cavities that degrade bone as they move across the bone surface (*13*). All bone cells reside within the bone matrix. The bone matrix consists of calcium-phosphate mineral, in the form of hydroxyapatite, along with high levels of collagen I. The bone matrix provides a structural framework that anchors cells as well as houses a rich source of growth factors that sustain resident cells (*14–16*). It is the dynamic interplay between these cells and their mineralized environment that maintains the structural integrity of bone (*10*).

For PCa to form a new bone destructive metastatic lesion, several processes must happen. PCa cells must first physically move to bone, doing so by traversing the circulatory system. Once PCa cells are in the bone microenvironment, they co-opt the normal balance described above into a pathogenic state. Individual elements of this process interact to form a positive feed-forward process referred to as the vicious cycle (*17–19*), Supplementary Fig. 1. Briefly, PCa cells secrete cytokines, e.g. interleukin 11 (IL-11), parathyroid hormone-related protein (PTHrP), that stimulate osteoblasts and osteocytes to increase secretion of several factors. A key factor secreted by osteoblasts and osteocytes is receptor activator of nuclear factor-κB ligand (RANKL).

RANKL recruits and stimulates the maturation of osteoclast progenitors. Mature osteoclasts degrade bone matrix, thereby releasing factors created by osteocytes embedded within the matrix that stimulate PCa cell growth, e.g. insulin-like growth factor 1 (IGF-1) (*20–22*). This growth of PCa cells returns us back to the beginning of the vicious cycle. We recently demonstrated that osteocytes, which represent over 90% of bone cells (*23*), play a vital role in stimulating the growth of human PCa cells and that this in turn is regulated by the structural make up of their microenvironment (*24*).

Cell motility represents a fundamental cellular process, and its dysregulation plays an essential role in the pathogenic process of PCa-mediated bone destruction. It is required for cells to move to bone, prototypically from their primary site. It is also required for the vicious cycle of bone destruction, where cooperatively engaged cells must move across the bone surface in order to degrade it. Until recently, it had not been possible to therapeutically target cell motility with precision. This is because it is regulated by a wide array of signaling proteins, but none of them were specific to the process of cell motility. Using cheminformatics to interrogate biological processes our group synthesized the first precision-acting therapeutic, KBU2046, that selectively inhibits cell motility (*25*). We demonstrated that KBU2046 works by inhibiting the phosphorylation of Ser338 on Raf1’s negative charge regulatory region (N-region), selectively regulating Raf1 function. KBU2046 is not a Raf1 kinase inhibitor. Inhibitors that bind Raf1’s active site inhibit its kinase activity, induce cell death and are systemically toxic. KBU2046 lacks all of these effects. Raf1 regulates many processes and has been deemed a “gatekeeper of cell fate”. Through KBU2046, we have been able to selectively regulate this gatekeeper protein so that the only downstream effects are on motility, and systemic toxicity is nil.

We previously demonstrated that when KBU2046 is given prior to intracardiac (IC) administration of human PCa cells into mice it prevents cells from trafficking to and destroying bone (*25*). It was unexpected, however, to find that it had no effect when given after PCa cells had already trafficked to bone. A pharmacokinetic analysis of KBU2046 demonstrated a short half-life of 2.3 hours and supported a suboptimal exposure time in bone. Separately, we demonstrated the additive therapeutic effects of KBU2046 given in combination with the bisphosphonate, zoledronic acid (*26*). Zoladronic acid binds to and chemically alters the bone microenvironment; it is widely used clinically in patients with metastatic PCa. Additive efficacy was observed with *in vitro* models of bone destruction, as well as in murine models of human PCa mediated bone destruction. The latter was also a pretreatment prevention model.

We hypothesized that antimotility therapy could be delivered to and retained in bone by chemically linking KBU2046 to a bisphosphonate. Further, the combined pharmacologic action of the resultant heterobifunctional compound would result in high therapeutic efficacy in established metastatic lesions. Here we describe our synthesis of heterobifunctional compounds, termed Dual-Acting Bone-Defending agents (DABDs). We demonstrate their ability to inhibit bone destruction and cancer cell motility *in vitro*. We demonstrate the ability of one compound, BTE-EN1, to prolong life in a prevention model, which has never been reported before. We demonstrate that BTE-EN1 is highly active at inhibiting bone destruction even in the face of established and advanced lesions. BTE-EN1 was non-toxic, did not interfere with standard-of-care hormone therapy or chemotherapy, and was active against multiple types of bone-destructive cancers. Together, this study identifies a highly active first-in-class agent that operates through a novel mechanism, addresses a long-standing clinical need, and has high potential for translation into humans.

## RESULTS

### Synthesis of Dual Acting Bone Defender Agents

Six different compounds were synthesized (Fig. 1 and Supplementary Table 1). Each compound consists of a bisphosphonate group at one end, a linker group in the middle, and a KBU2046 group at the other end. As such, we refer to them as DABDs. Alendronic acid (alendronate) and zoledronic acid (zoledronate) are each used clinically to inhibit bone resorption (*27*). We therefore examined both alendronate and zoledronate. BTE-EN1, BTE-EN2 and BTE-EN3 have alendronate as the bisphosphonate, differ by the length of the linker moiety, and have molecular weights ranging from 662 to 751. BTC-GN1, BTC-HN1 and BTC-IN1 have zoledronate as the bisphosphonate, differ by linker length, and have molecular weights ranging from 760 to 848.

**Fig. 1.**
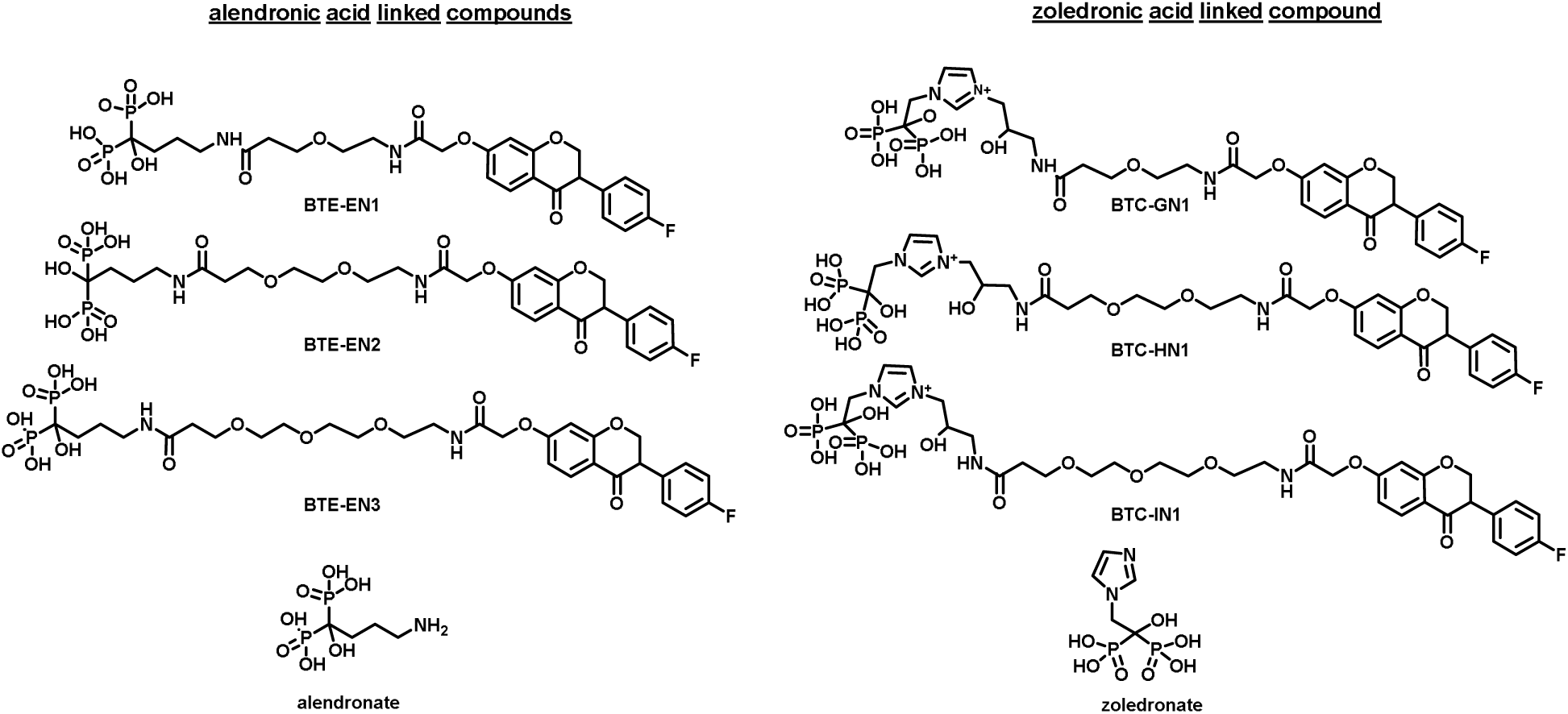
Chemical structures of Dual Acting Bone Defenders (DABDs), alendronate, and zoledronate. BTE-EN1, BTE-EN2 and BTE-EN3 are alendronate linked compounds. BTC-GN1, BTC-HN1 and BTC-IN1 are zoledronate linked compounds.

Detailed description of the synthesis and purification of DABDs are provided in Supplementary Information. These studies define new synthetic pathways that lead to the development of a class of new chemical entities. DABDs support incorporation of bisphosphonate groups and groups that inhibit cell motility into a single chemical whose molecular weight is below 900, and whose water solubility constitutes a desirable pharmacologic property.

### Functional Screening of DABDs

Two up front functional screens were performed in parallel: one for inhibition of bone destruction, the other for inhibition of cell motility. In an Osteo Assay, RAW264.7 cells differentiate into mature osteoclasts under the influence of RANKL, degrade hydroxyapatite matrix coated plates and the resultant decrease in bone surface area is quantified. This assay has previously been used to measure the activity of bisphosphonates (*28*). All six compounds exhibited dose-dependent and significant inhibition of osteoclast-mediated bone mineral degradation (Fig. 2, a to c). With alendronate-based DABDs, 3 of 3 significantly increased bone surface area by over 150% compared to controls at 20 µM, with 1 of 3 significantly increasing it at 1 µM. Zoledronate-based DABDs are more potent, with 3 of 3 significantly increasing bone surface area by over 175% at 20 µM, and 2 of 3 significantly increasing it at 1.0 µM. This finding is consistent with a prior report demonstrating that zoledronate is 30 to 100 fold more potent than alendronate (*29*). Interestingly, alendronate-based DABDs, BTE-EN2 and BTE-EN3, along with alendronate itself, exhibited a biphasic effect, enhancing bone degradation at lower concentrations while inhibiting it at higher concentrations. Importantly, BTE-EN1 did not exhibit a biphasic effect, nor did zoledronate-based DABDs. KBU2046 was previously shown by us to inhibit osteoclast-mediated bone mineral degradation (*26*). It was included as an additional control. Together, these findings demonstrate that DABDs inhibit osteoclast-mediated degradation of bone minerals in a concentration-dependent fashion.

**Fig. 2.**
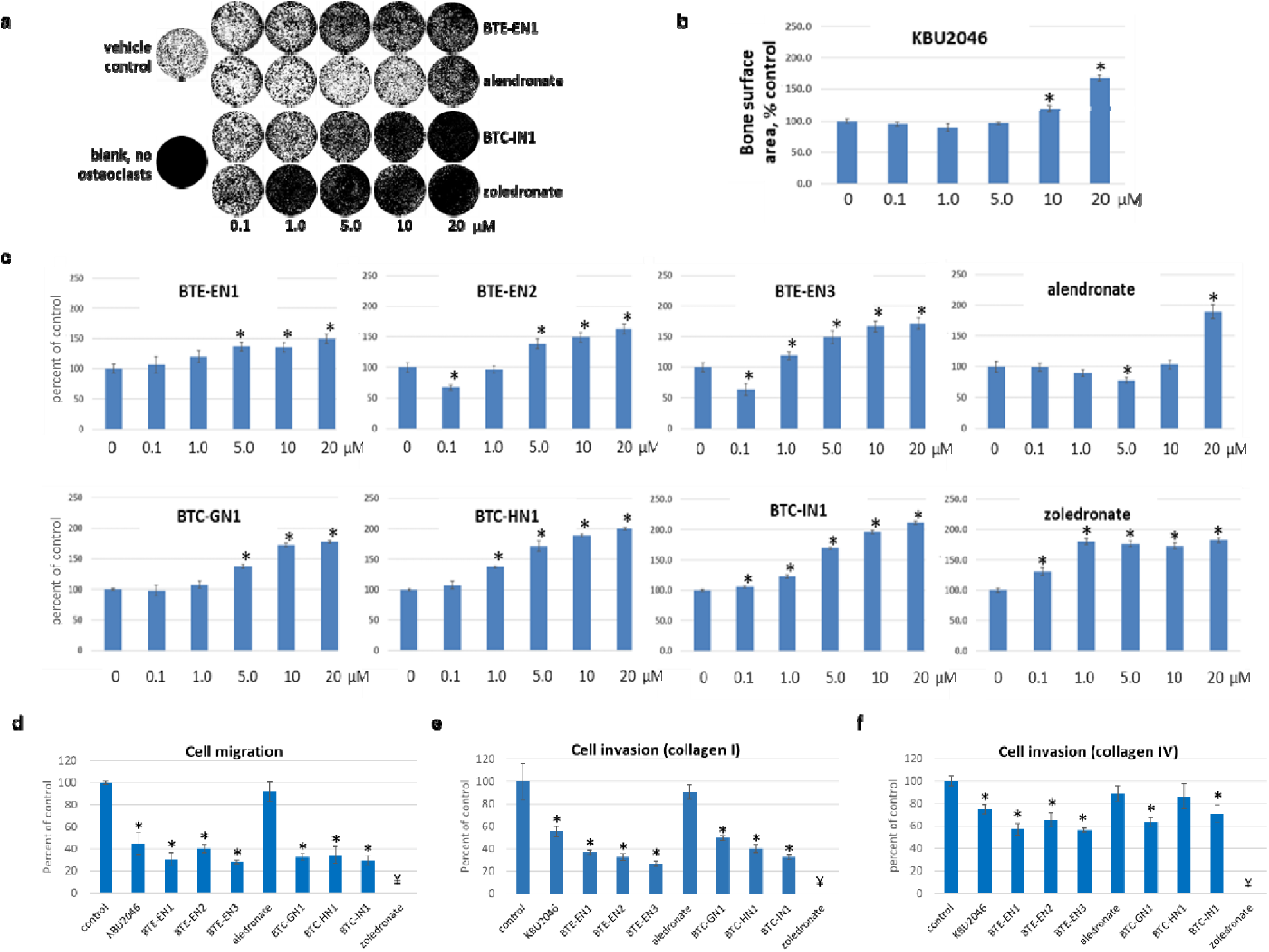
DABDs inhibit osteoclast-mediated bone degradation and cell motility. **(a-c)** RAW 264.7 cells, RANKL and the denoted concentrations of DABDs, alendronate, zoledronate or KBU2046, were added to Osteo Assay plates. After 6 days incubation, the area of remaining bone matrix was quantified. Control wells contained vehicle, while blank wells lacked osteoclasts. (a) Representative photomicrographs of wells treated as denoted. Quantification of bone area in wells treated with KBU2046 (b) and denoted compounds (c). (**d-f**) PC3 cells were treated with denoted compounds at 10 µM for a total of three days, and effects upon migration (d) and invasion through collagen I (e) or collagen IV (f) were measured. ¥: under these assay conditions zoledronate induced cell death. Data are normalized to the percent of control, expressed as mean ± SEM (N=3); * denotes Student’s t test P ≤ 0.05, compared to control; similar findings were observed in a repeat experiment conducted at a separate time.

The second up front screen was for inhibition of cell motility, including cell migration and cell invasion. Cell migration measures movement of cells across a surface, while cell invasion constitutes a composite measure of cell migration coupled to protease digestion and measures the ability of cells to invade through protein. Increases in cell migration and invasion are both associated with an increased propensity to form metastasis and are considered basic characteristics of the metastatic phenotype. As any drug that induces cell toxicity will give a false positive reading in a cell motility assay, all DABDs were tested for such, and none were found to reduce cell viability compared to control cells. All six DABDs significantly inhibited migration of human PCa PC3 cells (Fig. 2d). Compared to control, alendronate-based DABDs significantly decreased migration by 60-72% and zoledronate-based DABDs significantly decreased migration by 66-71%. Alendronate itself had no significant effect. While zoledronate significantly decreased migration by 40%, we found that under these assay conditions it was toxic to cells, and therefore, we did not consider resultant migration data meaningful. KBU2046, previously shown by us to inhibit cell migration and cell invasion (*30*), serves as a positive control and it significantly decreased migration by 55% compared to control.

Next, we examined cell invasion using collagen I and collagen IV (Fig. 2, e and f). Collagen I is a major extracellular matrix constituent of bone. It plays an important role in mediating binding of cancer cells, including PCa, to bone, and is in turn degraded during bone destruction (*31, 32*). Collagen IV is a central component of the basement membrane underlying epithelial cells. Invasion of *in situ* cancer cells through the basement membrane is a *sine qua non* characteristic for the clinical definition of cancer, and its presence across a wide array of pre-clinical models and clinical scenarios serves to portend the development of distant metastasis.

With collagen I, responses emulated the migration assay findings, with all six DABDs significantly inhibiting invasion. Compared to control, alendronate-based DABDs decreased invasion by 63-73% and zoledronate-based DABDs decreased invasion by 50-67%. With collagen IV, alendronate-based DABDs decreased invasion by 35-44%, while for zoledronate-based DABDs, only BTC-GN1 and BTC-IN1 were active, decreasing invasion by 29-36%. For migration and invasion assays cells were pre-treated in culture dishes, detached, and then placed into Boyden chambers. Under these assay conditions it was observed that zoledronate induced cell death and thus zoledronate data is not depicted. For both collagen I and IV, KBU2046 significantly decreased invasion and alendronate had no effect. Together, these findings demonstrate that DABDs inhibit cell motility, demonstrating activity across multiple different assays.

### Evaluation of DABDs’ efficacy and toxicity

We next examined whether DABDs were effective systemically. Movement through the circulation is required for cancer cells to form metastasis in a distant organ. PCa cells circulating in the blood of humans are known to portend the future development of bone metastasis (*33*). We have previously shown that such circulating cells have increased metastatic potential (*34*). To determine whether DABDs would inhibit circulating human PCa cells from metastasizing to and destroying bone, we used an intracardiac (IC) injection murine model wherein PC3-Luc cells metastasize to bone, grow and induce bone destruction (Supplementary Fig. 2). Their high propensity to metastasize to and destroy the jawbone ultimately leads to a moribund status, thereby providing a measure of the impact of PCa in humans. This model has been extensively used by us and others (*25, 26, 35, 36*).

We used this model to measure the toxicity and efficacy of three different DABDs. BTE-EN1 was selected to include as an alendronate-based DABD. It was efficacious in Osteo Assays and lacked the biphasic effects of other alendronate-based DABDs, it performed well across motility assays, and its associated synthetic route provided relatively high yields. BTC-HN1 and BTC-IN1 were selected because they were zoledronate-based compounds, exhibited relatively high efficacy in Osteo Assays, and exhibited relatively high efficacy in cell migration and collagen I invasion assays.

Beginning 3 days prior to injection of PC3-Luc cells, mice were treated with weekly intraperitoneal (IP) injections of vehicle (control), BTE-EN1, BTC-HN1 or BTC-IN1, with each compound given at 10, 100 and 1000 µg/kg (Fig. 3a). Mice given zoledronate-based DABDs, BTC-HN1 or BTC-IN1, experienced high rates of early death attributed to drug toxicity (Supplementary Fig. 3). In contrast, BTE-EN1 was well tolerated and highly efficacious (Fig. 3). There was no evidence of toxicity in mice given BTE-EN1 (Supplementary Fig. 4). As bone destruction increases, the ability of mice to consume food is impacted. This is reflected in the weight of control and 10 µg/kg cohort mice beginning to decline after 2 weeks post cell injection. In contrast, 100 and 1000 µg/kg cohorts continued to increase through week 3, thereafter declining. At 4 weeks, the weight of control mice was less than that of the 1000 µg/kg cohort, albeit not statistically significant, and was 27.1 ± 1.2 g (mean ± SEM) and 28.4 ± 1.2 g, respectively. As quantified by CT, BTE-EN1 treatment at 10, 100, and 1000 µg/kg significantly decreased mandibular bone destruction by 41 ± 3.5%, 58 ± 4.7%, and 56 ± 2.5% (mean ± SEM) compared to control (Fig. 3b). The further decrease in destruction with 100 and 1000 µg/kg doses was significantly different from the 10 µg/kg cohort, but not from each other, consistent with a plateau in therapeutic effect. Weekly In Vivo Imaging System (IVIS) imaging of the mandible demonstrates the expected growth kinetics in controls, and diminished growth in treatment cohorts (Supplementary Fig. 5). Of high importance, a significant dose-dependent increase in survival was observed (log rank P = 0.0071) with 100%, 80%, 70% and 50% of mice alive at 28 days treated with 1000, 100, 10, and 0 µg/kg BTE-EN1, respectively (Fig. 3c). To our knowledge non-cytotoxic bone-directed agents have never been reported to prolong life in this model. While the coupling of cytotoxic agents to bisphosphonates decreases bone destruction through induction of toxic effects on cancer cells, it also destroys bone marrow and is highly toxic (*37*). These are not features observed with BTE-EN1. Together, these studies identify BTE-EN1 as our lead agent. We demonstrate that it inhibits bone destruction by human PCa cells and prolongs life in a dose-dependent manner, and that it is well tolerated. All further investigations focused on BTE-EN1.

**Fig. 3.**
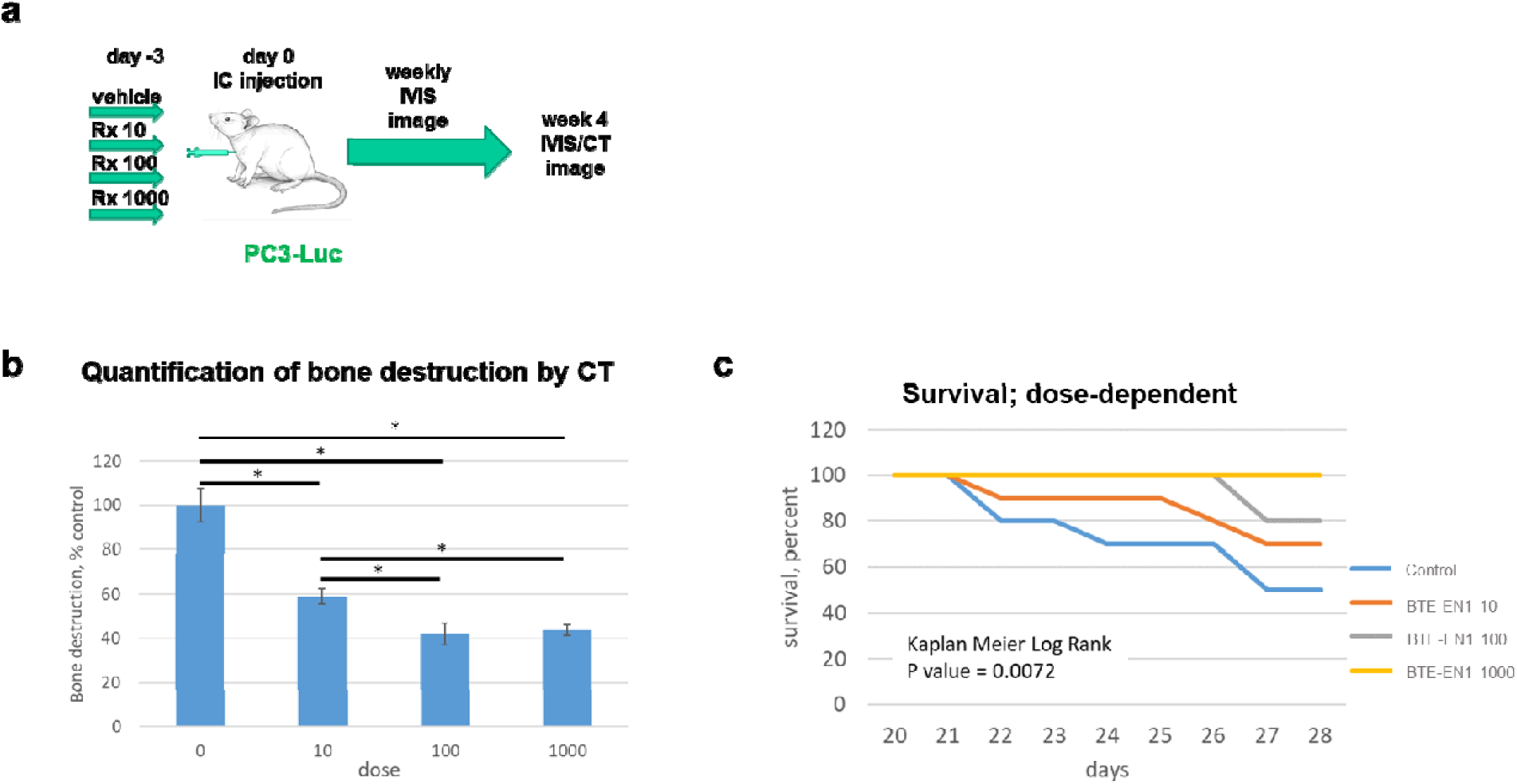
BTE-EN1 inhibits human PCa metastasis to bone and prolongs survival. (**a**) Experimental design of IC injection model. Three days prior to IC injection of PC3-Luc cells, N=10 mice/cohort began weekly IP treatment with 10, 100 or 1000 µg/kg BTE-EN1, with N=10 control mice receiving vehicle. (**b**) Mandibular bone destruction quantified by CT imaging. (**c**) Survival curve. Data are normalized to the percent of control, expressed as mean ± SEM; * denotes Student’s t test P ≤ 0.05, compared to control.

### BTE-EN1 inhibits bone destruction by established PCa metastasis

The IC model used above is well suited for examining whether BTE-EN1 would prevent circulating PCa cells from reaching and destroying bone, which it did. Its limitations relate to the fact that the biology of jawbone differs from that of other bones in the body (*38–41*), particularly as it relates to examining established PCa metastasis. Clinically, PCa almost never metastasizes to the jaw. Also, the jaw is uniquely susceptible to undergoing osteonecrosis in patients receiving bone-stabilizing agents (*42*). In contrast, PCa very commonly metastasizes to the long bones of the legs. With the intra-caudal artery (CA) injection model (*30*), cancer cells are injected into the caudal artery in the tail, and under the hydrostatic pressure of injection they travel retrograde into the pelvis, whereupon they enter the iliac arteries and selectively deposit in the long bones of the legs (Supplementary Fig. 6). These features model the clinical scenario in humans.

Here we used the CA injection model to examine whether BTE-EN1 has activity against established bone metastasis (Fig. 4, a to d). We demonstrated that IVIS signal of injected cells increased in the legs of mice over a 2-week period, indicating ongoing cell growth (Supplementary Fig. 7, a and b). After two weeks mice were randomized to weekly IP injection with 1000 µg/kg BTE-EN1 or vehicle (Supplementary Fig. 6, c and d). Quantification by CT demonstrated that BTE-EN1 significantly decreased bone destruction by 62 % compared to control mice (Fig. 4, b and c). The weight of control mice started to drop at 5 weeks, while the weight of BTE-EN1 mice continued to increase until 6 weeks and then gradually declined (Fig. 4d). Of interest, weekly IVIS imaging showed no significant difference between BTE-EN1 and control mice (Supplementary Fig. 7e). It is important to consider that IVIS imaging relies upon multiple factors impacting light transmission. Bone does not readily transmit light and bone undergoing degradation constitutes a highly variable situation. In contrast, CT imaging provides a direct quantitative gold-standard measure of bone destruction. Together, these data demonstrate that BTE-EN1 markedly decreases bone destruction in well-established bone destructive human PCa metastasis, and that it is well tolerated when administered systemically to mice.

**Fig. 4.**
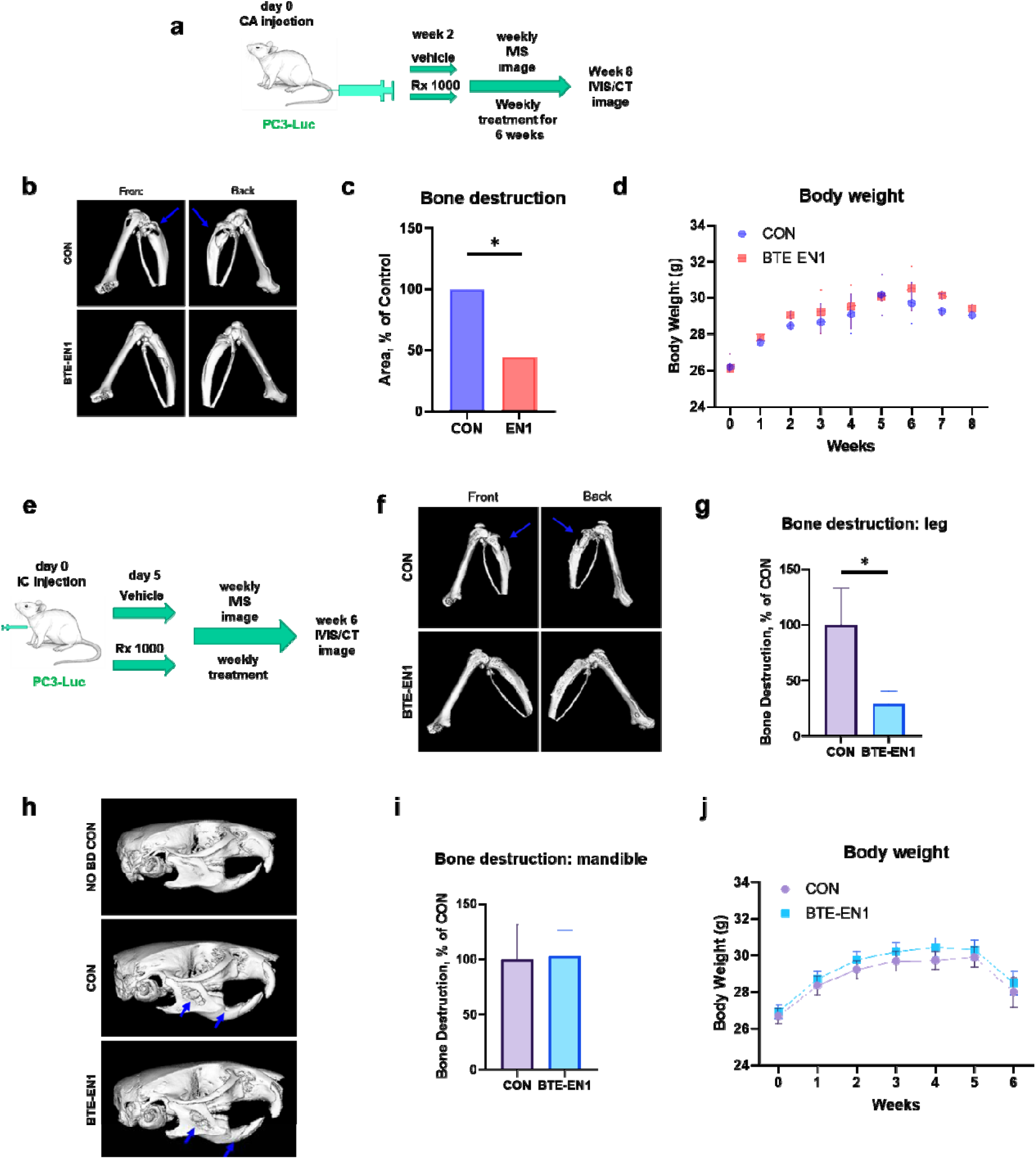
BTE-EN1 inhibits bone destruction in established PCa metastasis. (**a**) Experimental design of CA injection model of PCa bone metastasis (control group N=9; treatment group N=10). (**b**) Representative images of leg bone destruction in control and BTE-EN1 treated CA injection mice. Blue arrows denote large destructive lesions in controls. (**c**) Bone destruction in legs quantified by CT imaging. (**d**) Weekly body weight measurement of CA injection mice. (**e**) Experimental design of IC injection model of PCa bone metastasis. Five days post injection, 46 mice were randomized to control and treatment groups, N=28 per group. (**f**) Representative images of leg bone destruction in control and BTE-EN1 treated IC injection mice. Blue arrows denote large destructive lesions in controls. (**g**) Bone destruction of legs quantified by CT imaging. (**h**) Images depict bone destructive lesions in control and BTE-EN1 treated IC injection mice, compared to mice without metastasis to the jaw. (**i**) Bone destruction of mandible quantified by CT imaging. (**j**) Weekly body weight measurement of IC injection mice. Data are expressed as mean ± SEM; * denotes Student’s t test P ≤ 0.05, compared to control.

Many differences between the microenvironments of the jaw and the long bones of the legs have been identified (*43–45*). These differences have been linked to clinical findings that PCa metastasizes to bones of the legs and not the jaw, and that bone targeted therapeutics induce osteonecrosis selectively in the jaw (*46–48*). While the IC injection model is characterized by a propensity for PCa cells to localize to the jawbone, due to their widespread systemic distribution, we have found that ∼20% of mice metastasis can be detected in the long bones of the legs by IVIS imaging. We therefore sought to examine whether BTE-EN1 would exert differential therapeutic efficacy in bone of the jaw versus that of the legs using the IC injection model.

Treatment with 1000 µg/kg BTE-EN1 versus vehicle was begun 5 days post injection of PC3-Luc cells and continued for 6 weeks (Fig. 4e). Bone destruction measured by CT in the leg of BTE-EN1 treated mice was significantly decreased by 71% compared to control mice (Fig. 4, f and g), while there is no difference in the bone destruction of mandible between BTE-EN1 treated and control mice (Fig. 4, h and i). IVIS images yielded a similar pattern, with BTE-EN1 having no effect on the head but were significantly decreased in legs (Supplementary Fig. 8).

The body weights of both control and treatment groups began to decrease at 5 weeks after cell inoculation (Fig. 4j). These findings demonstrate that BTE-EN1 selectively decreases bone destruction by established metastasis in the leg but not in the mandible.

### BTE-EN1 retains the molecular mechanisms of action of its constituents

BTE-EN1 contains a bisphosphonate group connected by a linker to a KBU2046 group. We therefore hypothesized that it should exhibit mechanistic effects associated with each group. The two phosphate groups on bisphosphonates form ionic bonds with bone minerals. Bone mineral consists of calcium and phosphate, in the form of hydroxyapatite. We analyzed the ability of BTE-EN1 to bind hydroxyapatite using a fast protein liquid chromatography (FPLC) assay, as previously reported (*49*). As shown in Fig. 5a and Supplementary Fig. 9, BTE-EN1 binds hydroxyapatite with a retention of time of 28.15 ± 0.2 minutes (mean ± SEM). Zoledronate similarly bound hydroxyapatite, had a retention time of 25.26 ± 0.3 minutes (mean ± SEM). While it was in a similar range of that of BTE-EN1, it was distinct (P<0.001). Notably, KBU2046 does not bind.

**Fig. 5.**
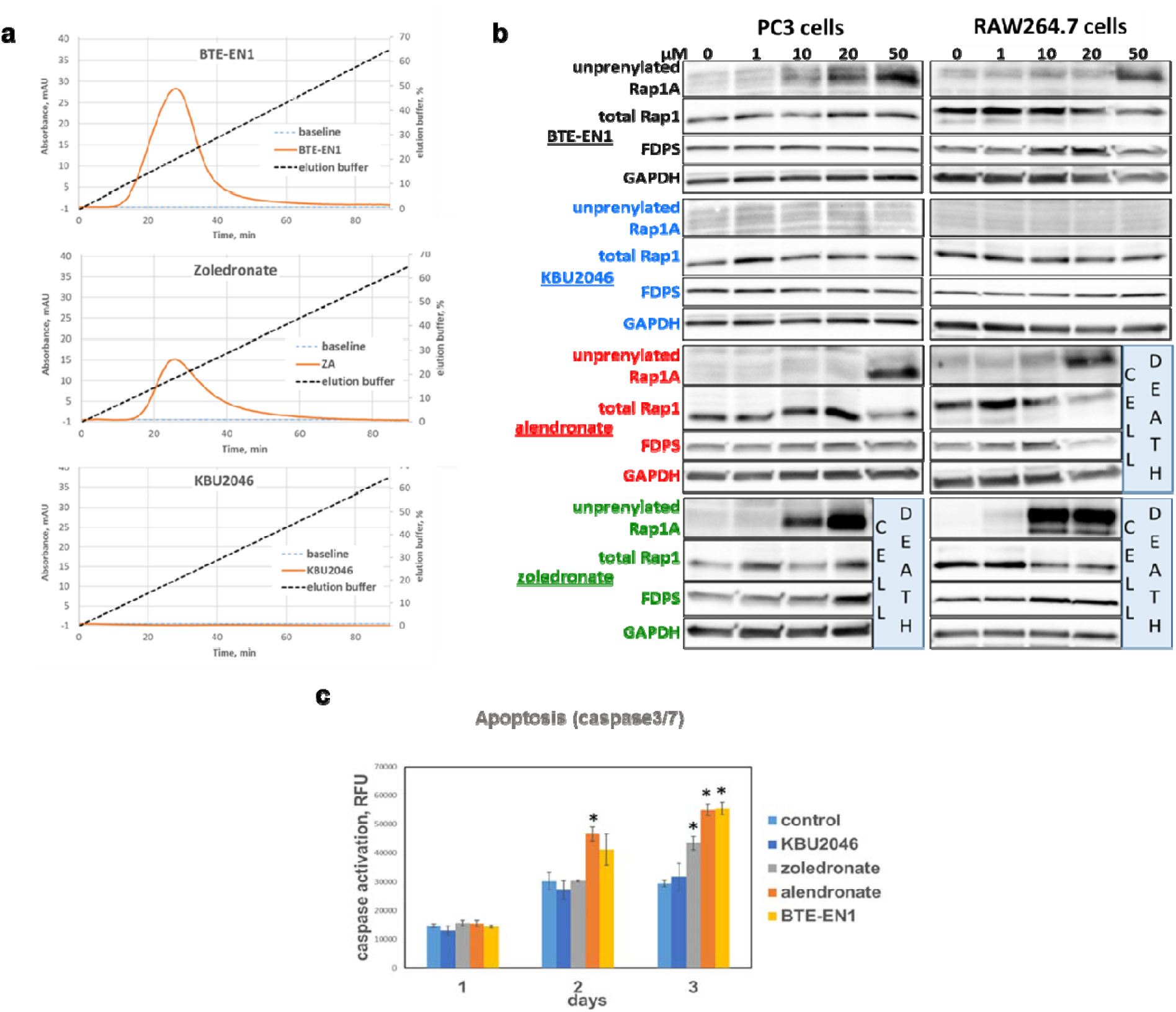
BTE-EN1 retains properties of bisphosphonates. (**a**) BTE-EN1 binds hydroxyapatite. Representative hydroxyapatite column chromatograms of BTE-EN1, zoledronate (ZA) and KBU2046 are depicted. (**b**) BTE-EN1 inhibits protein prenylation. PC3 and RAW264.7 cells were treated for 3 days at denoted concentrations and probed for the indicated proteins by Western blot. In instances where high drug concentrations induced cell death, lanes are so denoted. (**c**) BTE-EN1 induces apoptosis. RAW264.7 cells were treated with 10 µM drug or vehicle, as denoted, for 1-3 days, and then caspase activation was measured by ApoTox-Glo™ Triplex Assay. Data are the mean ± SEM (N=4) of a single experiment, with similar findings in a separate experiment (N=4). * Denotes Student’s t test P ≤ 0.05, compared to control.

After binding hydroxyapatite in bone, bisphosphonates are taken up into osteoclasts when they degrade bone. Once inside osteoclasts, nitrogen containing bisphosphonates, such as alendronate and zoledronate, inhibit farnesyl diphosphate synthase (FDPS) within the mevalonate pathway. This inhibits farnesyl pyrophosphate formation, in turn resulting in decreased prenylation of small GTPases, such as Rap1A (*50–55*). Evaluating the inhibition of prenylation is an accepted method for measuring bisphosphonate efficacy (*52, 54, 56*). Rap1A is prenylated on its carboxy terminus cysteine 181 residue. It is possible to detect changes in prenylation status by Western blot using an antibody that recognizes an epitope spanning amino acids 159-184 only when not prenylated (*57*). BTE-EN1 increased levels of unprenylated Rap1A in a concentration- and time-dependent fashion, and it did so in both osteoclasts (Fig. 5b) and PCa cells (Supplementary Fig. 10). KBU2046 had no effect. Zoledronate activity was observed at concentrations previously reported for this assay (*56*). It is about 100 folds more potent than alendronate and its greater potency is born out in this assay. At the highest concentrations zoledronate and alendronate both induced cell death (i.e., the missing lanes) (Fig. 5b). There appears to be a compensatory increase in FDPS with increases in unprenylated Rap1A, more pronounced in RAW264.7 cells and more pronounced with bisphosphonates, but evident with BTE-EN1 in RAW264.7 cells. Induction of apoptosis in osteoclast cells is a recognized and pharmacologically relevant effect of bisphosphonates (*50*). In Fig. 5c, we demonstrate that BTE-EN1 induces apoptosis in RAW264.7 cells, that effects increase with time, that both alendronate and zoledronate exhibit similar activity, but that KBU2046 lacks these effects. In Supplementary Fig. 11, we further confirmed that BTE-EN1, but not KBU2046, induces apoptosis in RAW264.7 cells by an orthogonal assay using Western blot to probe for cleaved-caspase 3. Together, these findings with BTE-EN1 are consistent with the characteristics of bisphosphonates, demonstrating its ability to bind bone mineral, inhibit Rap1A prenylation, and induce apoptosis in osteoclasts.

At the molecular level, KBU2046 inhibits phosphorylation of Ser338 on the activation motif of Raf1(*25*). We demonstrate in Fig. 6a that BTE-EN1 inhibits Raf1 Ser338 phosphorylation in PC3 cells. Raf1 is an actin binding protein that regulates actin function. Actin is involved in osteoclast-mediated bone degradation. In particular, it is necessary for the formation of the resorptive cavity in which bone degradation takes place. Under the influence of RANKL, RAW264.7 cells differentiate into mature osteoclasts characterized by formation of multinucleated cells that form well defined actin-ringed resorptive cavities, and expression of the maturation marker, tartrate resistant acid phosphatase (TRAP). As can be seen in Fig. 6, b to e and Supplementary Fig. 12, like KBU2046, BTE-EN1 inhibits formation of actin-ringed resorptive cavities, inhibits formation of multinucleated mature osteoclasts, and inhibits expression of TRAP. Together, these findings demonstrate that BTE-EN1 inhibits activation of Raf1 and inhibits maturation of osteoclasts, similar to the actions of KBU2046.

**Fig. 6.**
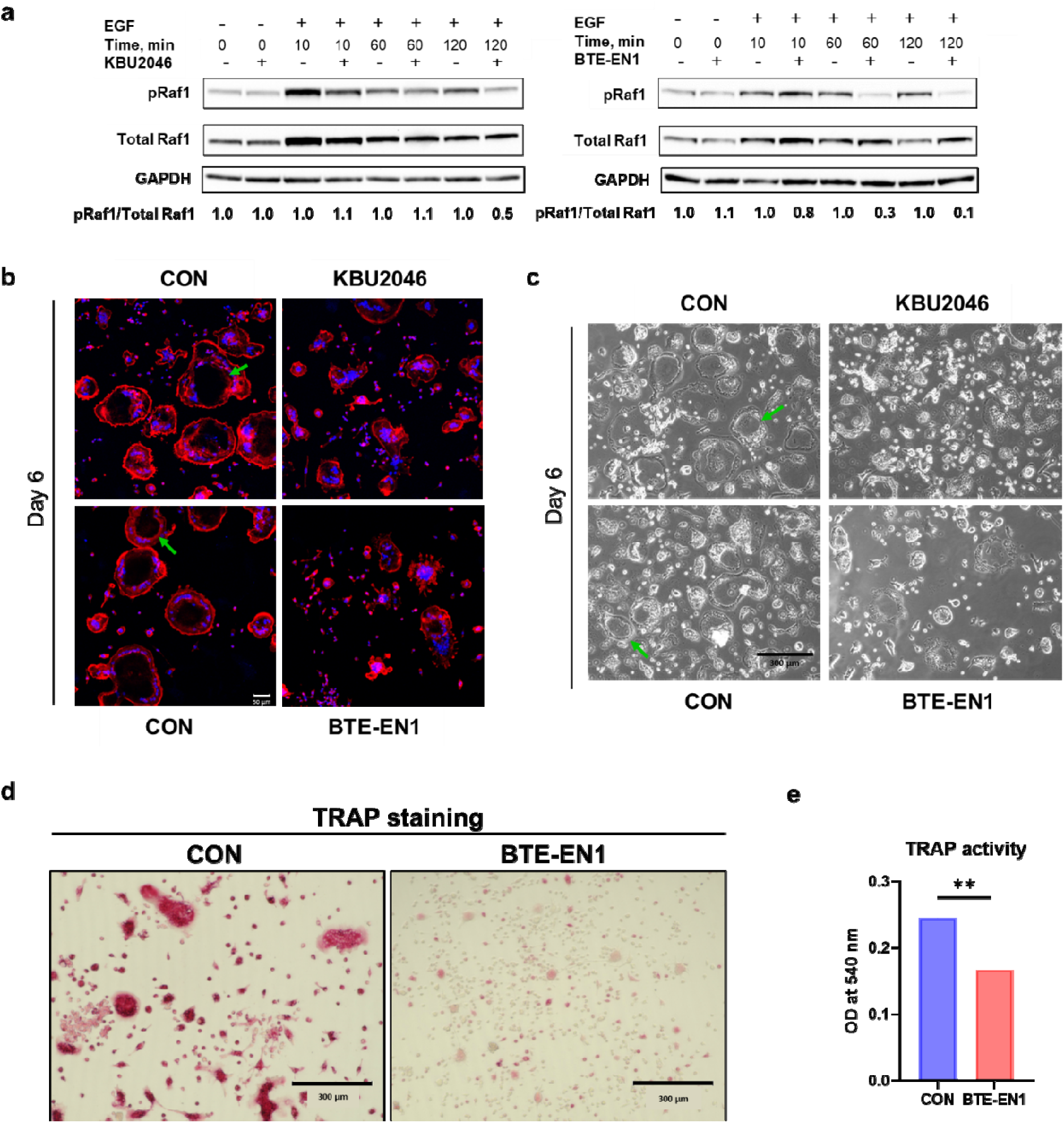
BTE-EN1 inhibits Raf1 phosphorylation and osteoclast maturation. (**a**) BTE-EN1 inhibits Raf1 phosphorylation of Ser338. PC3 cells were treated with 10 ng EGF, 10 µM BTE-EN1 or KBU2046, or vehicle for the denoted times, and phospho-Raf1, total Raf1 and GAPDH were detected by Western blot. (**b** and **c**) BTE-EN1 inhibits actin ring formation of osteoclasts. RAW264.7 cells were treated with 50 ng/mL RANKL and with 10 µM KBU2046 or BTE-EN1, or corresponding vehicle for 6 days, as denoted. Representative immunofluorescent images of cells were stained for actin (Rhodamine Phalloidin, red) and nuclei (DAPI, blue) are shown in (b), scale bar = 50 µm, while light microscopic images are shown in (c), scale bar = 300 µm. Green arrows denote actin-ringed resorptive cavities. (**d** and **e)** BTE-EN1 inhibits osteoclast maturation. (d) Representative images of TRAP staining are shown, scale bar = 300 µm. RAW264.7 cells were cultured on Osteo plates, treated with 50 ng/mL RANKL, 10 µM BTE-EN1 or vehicle for 6 days. TRAP was measured in cells (d) and media (e). Data are mean ± SEM (N=5) of optical density (OD); * Denotes Student’s t test P ≤ 0.05.

### Applicability of BTE-EN1 as a therapeutic

Low toxicity is important for a therapeutic. Clinically, BTE-EN1 was well tolerated. Treatment of BTE-EN1 is associated with maintaining body weight in the CA injection model (Fig. 4d) and prolongation of life in the IC injection model (Fig. 3c). To further examine BTE-EN1’s toxicity, non-tumor bearing CD1 mice were treated with 10,000 µg/kg of BTE-EN1 four times within a week. This dose was 4,000-fold higher than that required for efficacy (10 µg/kg in Fig. 3). Mice were evaluated for toxicity, inclusive of functional and microscopic analysis of critical organs/tissues. No evidence of toxicity was observed (Supplementary Fig. 13 and Supplementary Table. 2). Complementing the findings in CD1 mice, BTE-EN1 failed to demonstrate acute cytotoxic effects across the NCI-60 cell line screen (Supplementary Fig. 14).

BTE-EN1 was shown to be efficacious on human PCa PC3 cells, which induce osteolytic bone lesions, and on murine RAW264.7 osteoclast cells. If BTE-EN1 is to serve as an effective therapeutic, it will be important to demonstrate that it has broader efficacy. We demonstrate in Fig. 7a that BTE-EN1 inhibits cell motility across several human cancer cell types associated with bone destruction, including osteoblastic/osteolytic PCa C4-2B, lung cancer A549, and breast cancer MDA-MB-231 cells.

**Fig. 7.**
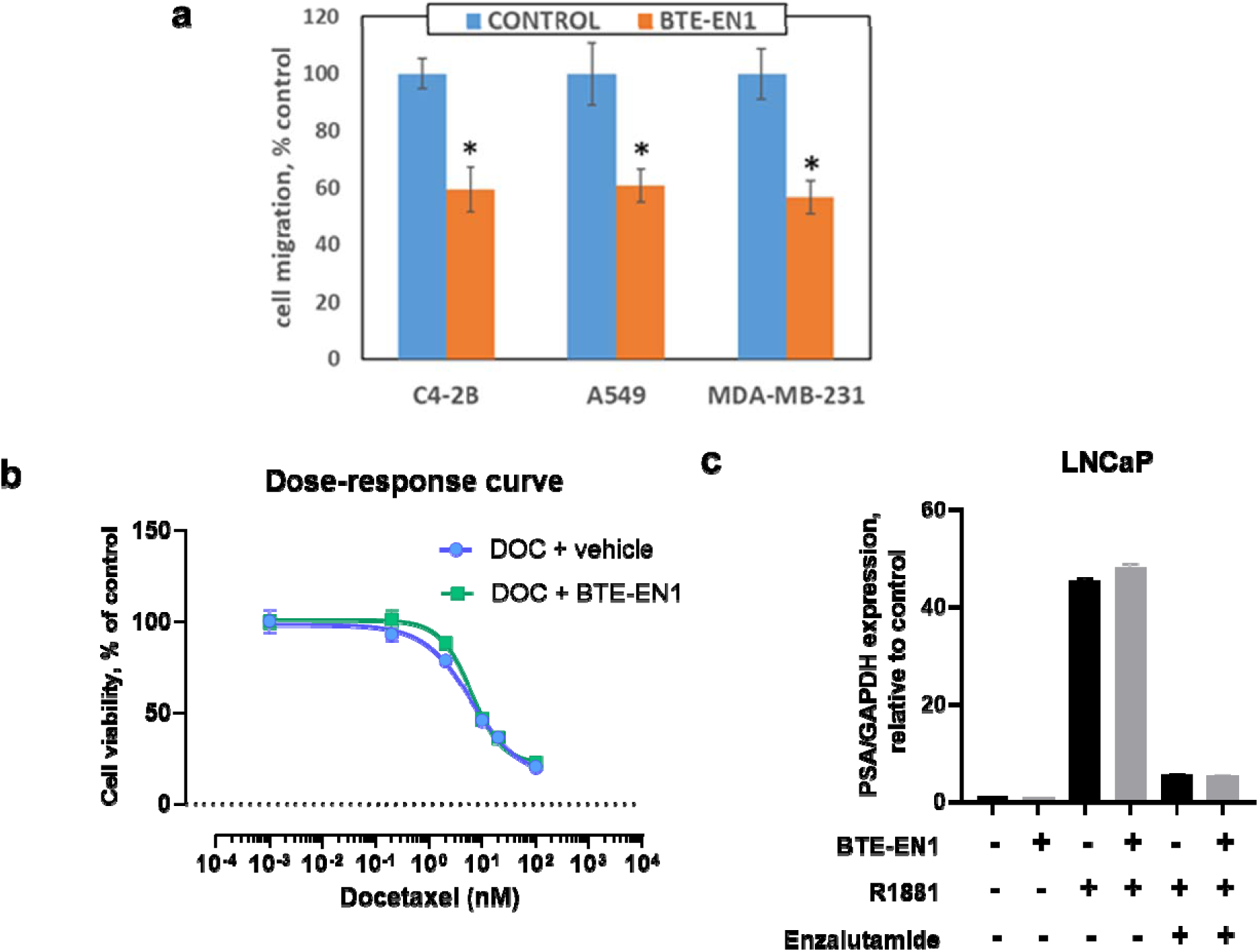
BTE-EN1 inhibits cell motility across a panel of cancer cell types and does not affect PCa therapies used in the clinic. (**a**) BTE-EN1 inhibits cell migration of multiple human cancer cell types. C4-2B prostate, A549 lung and MDA-MB-231 breast cancer cells were treated with 10 µM BTE-EN1 or vehicle control for 3 days and cell migration was measured by Boyden chamber migration assay. Data are the mean ± SEM, N=4. (**b**) BTE-EN1 does not interfere with cytotoxic agents. PC3 cells were treated with docetaxel (DOC) in the presence of 10 µM BTE-EN1 or vehicle for 3 days, and cell viability was measured by MTT assay. Data are the mean ± SD, N=8. (**c**) BTE-EN1 does not interfere with therapeutic alteration of the androgen receptor. LNCaP cells were cultured under hormone-free conditions, treated with 1 nM R1881 (synthetic androgen), 10 µM enzalutamide, 10 µM BTE-EN1, or respective vehicle for 24 hours as denoted. Expression of PSA was measured by qRT-PCR, and the ratio of PSA to GAPDH was expressed as the mean ± SEM % compared to vehicle only control. Data are the mean ± SEM, N=2; similar findings were found in two separate experiments (N=2). * Denotes Student’s t test, P ≤ 0.05.

BTE-EN1 exerts functions distinct from that of more conventional anti-cancer agents and in practice would be given along with them. Thus, it is important to ensure that BTE-EN1 does not inhibit their activity. Docetaxel is a cytotoxic chemotherapeutic agent used to treat patients with PCa and has been shown to prolong life. To investigate whether BTE-EN1 will interfere with its anti-cancer growth effect, we performed 3-day growth inhibition assays. In Fig. 7b, it is shown that docetaxel alone inhibited the growth of PC3 cells with an IC50 of 6.191 ± 0.3 nM (mean ± SEM), and that docetaxel plus BTE-EN1 had an IC50 of 6.087 ± 0.1 nM, which was not significantly different from that of docetaxel alone. Similar findings were observed with C4-2B cells (Supplementary Fig. 15). Docetaxel inhibits microtubule function, through inhibition of disassembly, thereby inhibiting cell cycle. In Supplementary Fig. 16 it is demonstrated that the cell cycle blockade caused by docetaxel is not altered by the addition of BTE-EN1.

Inhibiting the ability of androgen to drive PCa growth is a mainstay of treatment in the clinic. Androgen (R1881) induces upregulation of the androgen receptor (AR) responsive gene, prostate specific antigen (PSA) in human PCa AR positive LNCaP cells and this effect is blocked by the AR antagonist enzalutamide. We demonstrate that none of these actions are significantly impacted by the addition of BTE-EN1 (Fig. 7c). These experimental conditions emulate clinical treatment of PCa where androgen production is inhibited through medical or surgical castration and androgen receptor antagonists, such as enzalutamide, and therefore inhibit receptor activation. Collectively, BTE-EN1 was shown to lack toxicity both *in vivo* and *in vitro* and inhibit cell motility across different cancer cell types, and did not interfere with therapies used in the clinic to treat advanced PCa, all suggesting its high potential to be used as a therapeutic clinically.

## DISCUSSION

Cancer-mediated bone destruction is a common and longstanding clinical problem. Its clinical consequences serve to destroy and shorten lives. Current bone targeted therapeutics have limited efficacy and significant toxicity. Despite being available for several decades, they have not notably otherwise evolved. By focusing on inhibiting cell motility, we undertook an innovative and rational approach. It yielded high success.

Here, we describe the synthesis of heterobifunctional new chemical entities called, dual-acting bone defenders (DABDs). They represent first-in-class compounds. We demonstrate that they inhibit human PCa cell migration and invasion, as well as osteoclast-mediated bone destruction *in vitro*. Our lead compound, BTE-EN1, inhibits human PCa mediated bone destruction, demonstrating activity across multiple systemic models. In the IC injection mouse model BTE-EN1 is so active it prolongs life. This is an unprecedented achievement for non-cytotoxic bone targeted agents compared to previous reports (*58–60*). The CA injection mouse model represents a state-of-the-art model that emulates the scenario in humans (*30*). We refined it to create well-established PCa metastasis in the long bones of the legs, going on to demonstrate the high efficacy of BTE-EN1 against established bone-destructive PCa metastasis.

A series of assays demonstrate that BTE-EN1 retains the pharmacologic properties of its two constituent functional moieties. These assays involve measures of activity at the molecular level, as well as those that measure efficacy at the cellular level, together providing orthogonal and comprehensive assessments. Together our findings support the model depicted in Fig. 8.

**Fig. 8.**
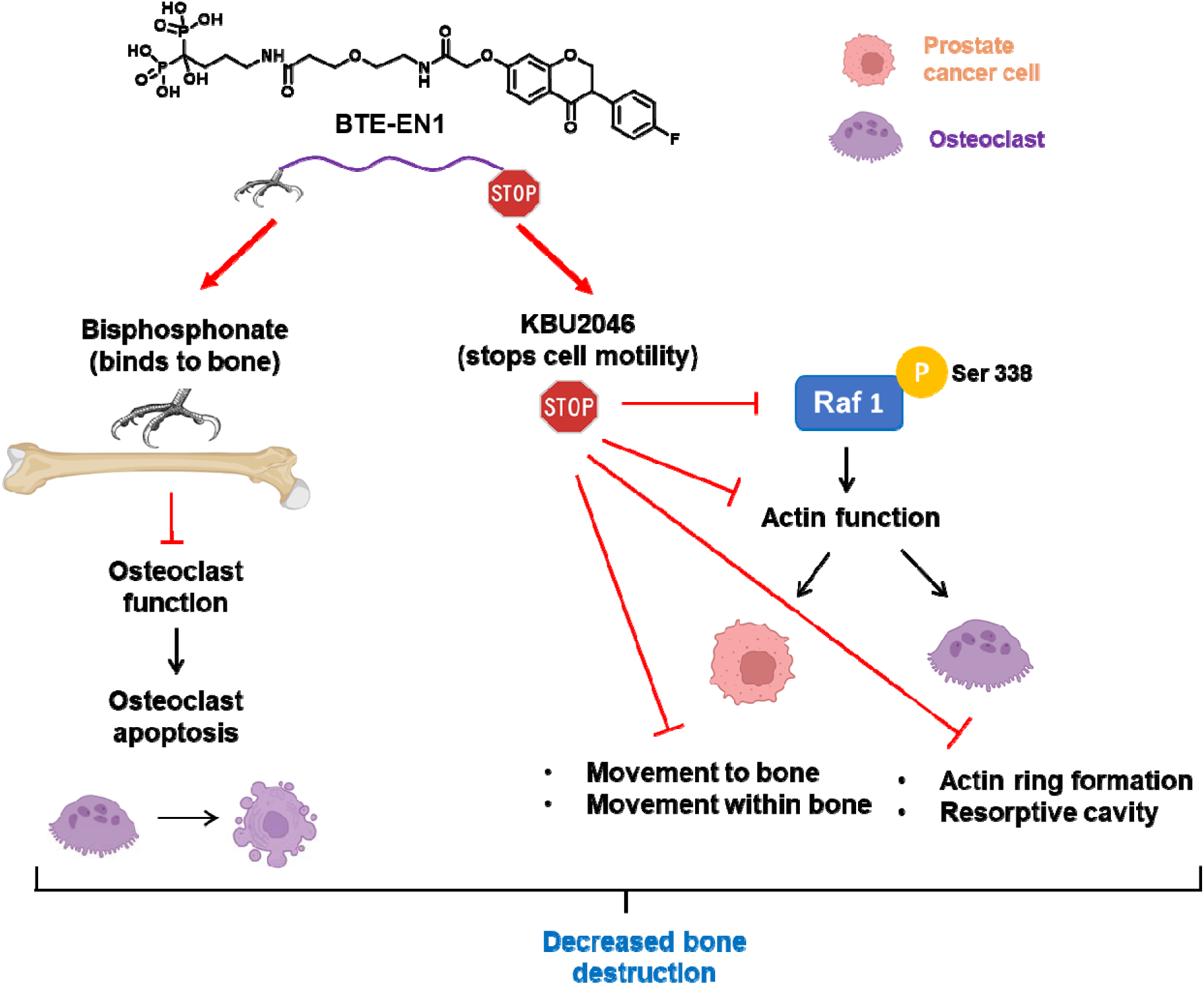
Graphic summary of BTE-EN1’s mechanisms of action. BTE-EN1 is composed of a bisphosphonate group, a linker, and a KBU2046 group. The bisphosphonate group binds to bone and induces apoptosis of osteoclasts. The KBU2046 group stops cell motility via downregulating phosphorylation of Ser338 on Raf1, thereby affecting actin function, resulting in inhibiting the movement of both PCa cancer cells and osteoclasts as well as inhibiting osteoclast resorptive cavity formation. Thus, BTE-EN1 treatment leads to decreased bone destruction via targeting these different mechanisms.

Consistent with the heterobifunctional chemical nature of BTE-EN1, it exerts multiple effects on different cell types and different functions that constitute basic elements of the vicious cycle. We have previously described the concept of combined “multi-functional” therapy and have demonstrated its therapeutic advantages (*26*). With BTE-EN1, we incorporate those functions into a single chemical entity and demonstrate high efficacy.

Our studies address several fundamental concepts important for considering translation into the clinic. First, we demonstrate lack of clinical toxicity across three systemic models. Two of them involved tumor-bearing mice, and in each clinical improvement was demonstrated, inclusive of prolongation of life (Fig. 3) and delay in cancer-associated weight loss (Fig. 4). In the third model, comprehensive tissue analysis demonstrated lack of toxicity at doses 4,000-fold higher than that associated with efficacy (Supplementary Fig. 13). Lack of toxicity across the 60 cell lines of the NCI-60 cell line screen provides another orthogonal measure of absence of toxicity (Supplementary Fig. 14). A second concept relates to our demonstrating efficacy across multiple cancer types, inclusive prostate, breast and lung cancer (Fig. 7a). While the current study focused on PCa, many other cancers similarly destroy bone, inclusive of very common ones such as lung and breast cancer. Thus, BTE-EN1 has broad impact potential. We also demonstrated efficacy on an osteoblastic human PCa cell line (Fig. 7a). While PCa is considered to cause osteoblastic lesions clinically, the actual bone destruction in these lesions is driven by osteolysis, and this was the focus of our systemic models. A third concept relates to whether BTE-EN1 can be administered with other clinically used agents. This is important practically, demonstrating that BTE-EN1 can be administered at the same time with cytotoxic and hormonal agents, both of which are known to prolong life in patients with advanced PCa. It is also important conceptionally, as BTE-EN1’s mechanism of action differs from these agents, setting up the possibility of their intentional combination to target multiple functions, thereby improving clinical efficacy.

Inhibiting the motility of PCa cells after they have already trafficked to bone may seem counterintuitive. It is important to consider that cells engaged in the vicious cycle are not docile. In order to degrade bone and remain in close proximity, they must move across the surface of the bone. Our findings support the notion that targeting movement is both pathogenically and therapeutically important. The current study does not prove that the movement of PCa cells in the bone is necessary for driving destruction. Future studies will pursue this fundamental question. We had previously reported that the administration of the anti-motility agent KBU2046 effectively prevented circulating PCa cells from moving into bone, but was inactive after cells had already established in bone (*25*). This would seem to support the notion that inhibiting motility after cells had arrived in bone was not effective. However, KBU2046 has a half-life of only a few hours and its anti-motility activity was similarly limited. These considerations led us to couple KBU2046 to bisphosphonates, which are known to selectively target and persist in bone.

We identified differential findings between gold standard CT-based imaging and that of IVIS imaging to measure bone lesions. This provides a cautionary finding for other investigators who use IVIS imaging to quantify tumors residing in bone. Light does not readily transmit through bone, and bone destruction adds a high degree of variability in transmitted light.

It was expected that zoledronate-based DABDs were more potent at inhibiting bone destruction *in vitro*. It was surprising to find that they were toxic in animals. Separate studies by us confirmed their purity and lack of toxicity *in vitro*. Possible explanations include an undetected impurity, and/or altered pharmacologic action potentially due to heretofore undefined metabolism.

In summary, we have synthesized a new class of therapeutics that consist of heterobifunctional chemical entities that bind bone mineral, inhibit cell motility, and exhibit high efficacy. Their ability to inhibit bone destruction by established metastatic bone lesions provides a novel therapeutic approach to a longstanding, common and very problematic clinical challenge. Our findings demonstrate that BTE-EN1 has little toxicity and is active against different types of cancer that frequently destroy bone, suggesting a high potential for its impactful translation into the clinic.

## METHODS

### Cell culture and reagents

Prostate cancer cell lines (PC3 and C4-2B), breast cancer cell line (MDA-MB-231), lung cancer cell line (A549), and osteoclast precursor mouse macrophage cell line (RAW264.7) were obtained from American Type Culture Collection (ATCC). The PC3-Luc cell line was obtained from PerkinElmer (#BW133416). All cell lines were cultured following ATCC’s handling guidelines and maintained at 37°C in humidified 5% CO_2_ incubators. All cell lines were drawn from stored stock cells and replenished on a standardized periodic basis and were routinely monitored for Mycoplasma (PlasmoTest™, #rep-pt1, InvivoGen or Mycoplasma Detection Kit, #13100-01, SouthernBiotech). PC3 and PC3-Luc cells were cultured in RPMI1640 medium containing 10% FBS and 1% Antibiotic-Antimycotic (#15240062, Gibco). RAW264.7 cells were maintained in DMEM medium containing 10% FBS and 1% Antibiotic-Antimycotic. MDA-MB-231 cells were cultured in RPMI1640 medium with 10% FBS and 1% Antibiotic-Antimycotic. A549 cells were cultured in F-12K medium containing 10% FBS and 1% Antibiotic-Antimycotic. Zoledronic acid (#S1314), alendronate (#S1624), and docetaxel (#1148) were purchased from Selleckchem.com and reconstituted per the manufacturer’s recommendations. EGF was purchased from Sigma (#E9644).

Cell lysis and Western blots were performed as described by us (2). Antibodies recognizing Rap1A/Rap1B (#2399), c-Raf (#53745), phospho-c-Raf (Ser338) (#9427), Cleaved-Caspase 3 (9664T), GAPDH (#2118), anti-mouse IgG-HRP linked secondary (#7076), and anti-rabbit IgG-HRP linked secondary (#7074) antibodies were purchased from Cell Signaling Technology. Anti-FDPS/FPS (#ab189874) antibody was purchased from Abcam. Antibodies recognizing Rap1A (sc-373968), β-Actin (# SC-47778) were purchased from Santa Cruz Biotechnology. Pierce ECL Western blotting substrate (#32106) and SuperSignal West Femto maximum sensitivity substrate (#34096) were purchased from Thermo Scientific.

### Bone degradation assay

The bone degradation assay was conducted as described previously by us (*26*). Briefly, a 50,000 cell/mL suspension of an early passage number of RAW264.7 cells in osteoclast differentiation media (MEM-alpha (Gibco), 10% FBS, 1% Antibiotic-Antimycotic, and 50 ng/mL RANKL (EMD Millipore, R0525)) was distributed into tubes. Calculated amounts for the indicated concentrations of DABDs (BTE-EN1, BTE-EN2, BTE-EN3, BTC-GN1, BTC-HN1, BTC-IN1), alendronate, zoledronate, or KBU2046, were mixed in the tubes prior to pipetting 100 uL cell suspension into the wells of the 96-well Osteo Assay Surface plates (Corning). Control wells only contain vehicle. No RAW 264.7 cells were added to blank wells. Around day 3 or 4, 75% of the media was aspirated and replenished with fresh media containing RANKL and treatments. After 6-day incubation, the wells were aspirated, and 100 uL of 10% bleach solution was added. After 10 minutes incubation, the wells were aspirated and washed thrice with ultrapure water. The plate was patted on paper towels to expel water. In order for enhancing the contrast of the pits for digital imaging and downstream analyses, SilverQuest Silver Staining Kit protocol (ThermoFisher Scientific) was adapted to mimic the Von Kossa staining. Briefly, following the manufacturer’s recommendations, 100 uL of a freshly prepared staining solution was pipetted into the wells and incubated for 1 hour in the tissue culture incubator. Then, the wells were aspirated and washed thrice with ultrapure water, followed by addition of 100 uL of developing solution (containing Developer and Developer Enhancer) for 30 minutes at room temperature, and 100 uL stopper solution for 15 minutes. For determining the area of the remaining bone matrix, each well was imaged in parts as 4 overlapping quadrants using a 4X objective, or as 2 overlapping semi-circles with a 2X objective on an EVOS™ M5000 Imaging System. An ImageJ macro was developed for automated pairwise stitching of the images for each well, cropping off outer regions, contrasting, threshold adjustments, and bone surface area determination. All experiments repeated, each treatment condition had N=4 replicates.

### Cell migration assay

Boyden chamber cell migration assays were performed as described previously by us (*61*) in the conventional 24-well format and verified in 96-well high throughput screening plate formats such as CULTREX (3465-096-K) and Millipore (MAMIC8S10). PC3, C4-2B, A549, and MDA-MB-231 cells were cultured for 2 days in their complete growth media with the denoted compounds at 10 µM. The cells were harvested, and 50,000 cells were added in the Boyden insert in media containing 0.5% FBS, 0.2% BSA, with or without the same compounds of interest. The lower chambers (or wells) received serum-free media supplemented with 20 ng/mL EGF as a chemoattractant. After incubating for 24 hours, the top and the bottom wells were aspirated carefully. The bottom wells were filled with a cell dissociation solution containing Calcein AM and incubated for 1 hour with occasional tapings on the sides of the plate to dislodge the cells that had crossed the membrane. The inserts were removed, and the fluorescence intensities were recorded on a BioTek Synergy plate reader. The intensities were converted to cell numbers by comparing them to a standard curve that was separately generated from the same type of cells by incubating 0-100,000 cells in Calcein AM/Dissociation buffer for 1 hour. All experiments repeated, each treatment had N=4 replicates.

### Cell invasion assay

Boyden chamber cell invasion assays (*61*) were performed similarly as in the cell migration assays with a slight difference that the inner membranes of the Boyden chambers were optimally coated overnight (at 37 °C in humidified 5% CO2 incubator) with rat Collagen I (BD Biosciences) at 50 mg/mL, or with 0.125 mg/mL human type IV Collagen (BD Biosciences) prior to adding the PC3 cells into the Boyden chambers. All experiments repeated, each in N=4 replicates.

### Binding capacity to hydroxyapatite columns

Bio-Gel HT hydroxyapatite matrix (Bio-Rad # 1300150) in its original bottle was shaken well and a small volume was pipetted into a container and allowed to settle for 5 min. The supernatant was discarded and copious amounts of Buffer A (1 mM potassium phosphate, pH 6.8) were added to resuspend the precipitate. The precipitation and washing process was repeated thrice to eliminate fine particulates. A Diba BenchMark™ Microbore Chromatography Column with 2 Fixed Endpieces (3 mm x 25 mm) (Cole-Parmer #EW-11941-06) was freshly packed with the matrix under similar flowrate and pressure conditions for every column run on a GE AKTA Pure FPLC system. The packed bed volume was approximately 0.18 mL. After washing the column with Buffer A, about 125 nmol of compound BTE-EN1, diluted in Buffer A, was injected to the column from a capillary loop at a rate of 0.2 mL/min. The column was equilibrated with Buffer A for 10 min at 1 mL/min, followed by a shallow gradient elution from 0 to 65% with Buffer B (1500 mM potassium phosphate, pH 6.8) over 90 minutes at 1 mL/min. Similarly, the experiment was repeated with other loadings of BTE-EN1 and ZA (125, 250, 375, 500 and 750 nmol). The chromatograms were detected at 280 nm and 222 nm for BTE-EN1 and ZA, respectively. Also, a baseline curve was generated (without BTE-EN1 or ZA) using similar gradient and flowrate for 280 nm and 222 nm.

Chromatograms were analyzed with Cytiva’s UNICORN Evaluation Classic 7 package. The corresponding baseline curve was subtracted from each of the BTE-EN1 and ZA chromatographs, smoothed with moving average algorithm for 0.2 minutes length, and compared as an overlay. Separately, the peak retention time and area under the curve were determined for each chromatogram.

### Adsorption to hydroxyapatite

A 5 mL, 100 µM BTE-EN1 and ZA stock was prepared in 10 µM potassium phosphate buffer (pH 6.8). 200 µL of hydroxyapatite slurry (Bio-Gel HT Hydroxyapatite matrix, Bio-Rad # 1300150) was pipetted in six separate low binding Eppendorf 1.5 mL microcentrifuge tubes, centrifuged and buffer discarded. The matrix was washed twice with 1 mL of the phosphate buffer. 800 µL of the drug stocks (∼80 nmol) was dispensed in tubes in triplicates. Samples were incubated at room temperature for 30 minutes on a tube rotator. Tubes were centrifuged and the supernatants were pipetted into fresh tubes. The flow paths of a GE AKTA Pure FPLC system were equilibrated with the potassium buffer. A characteristic wavelength of 215 nm was selected to detect BTE-EN1 and ZA. A 500 µL sample injection capillary loop was attached, and 700 µL of untreated stocks were pumped through the system at 1 mL/min followed by additional 3 mL buffer for cleaning. This was repeated twice. Similarly, the hydroxyapatite treated BTE-EN1 and ZA supernatants were read. Finally, the phosphate buffer was read thrice for blank or baseline. Cytiva’s UNICORN Evaluation 7 package was used to calculate the area under the curves. The blank values were subtracted from the -/+ hydroxyapatite treated BTE-EN1 and ZA values. The amount of drug (BTE-EN1 or ZA) not adsorbed by hydroxyapatite was depicted as a percentage of the untreated stock solution.

### Apoptosis Assay

ApoTox-Glo™ Triplex Assay (Promega) was used to measure caspase-3/7 activation per manufacturer’s instructions. RAW264.7 or PC3 cells were seeded in 3000 cells/well in 96-well plates and cultured in osteoclast differentiation media for 3 or 4 days, then the media was replaced with fresh differentiation media containing 10 µM denoted drugs or vehicle for 1-3 days. Caspase activation was assessed at day 1, 2, and 3, respectively, by ApoTox-Glo™ Triplex Assay.

### Immunofluorescence

RAW264.7 cells (2 x 10^4^) were seeded in 15 mm diameter glass coverslips in 12 well plates and cultured in maintenance media (DMEM, 10% FBS, and 1% Antibiotic-Antimycotic). One day later media were replaced with osteoclast differentiation media (MEM-alpha, 10% FBS, 1% Antibiotic-Antimycotic, and 50 ng/mL RANKL (Peprotech, #315-11)) containing DMSO (vehicle control of KBU2046), 10 µM KBU2046, dH_2_O (vehicle control of BTE-EN1) or 10 µM BTE-EN1 for 4 or 6 days. The differentiation media was changed every 2 days, along with replacement compounds. Cells were fixed at day 4 or at day 6. Before staining, fixed cells were imaged on an EVOS™ M5000 Imaging System using the transmitted light channel. Cells were washed twice with PBS, followed by 15 minutes fixation in 4% paraformaldehyde, permeabilized with 0.1% Triton X-100 in PBS for 15 minutes, and blocked with 1% BSA in PBS for 1 hour at room temperature. Cells were incubated with rhodamine phalloidin (Invitorgen, #R415) diluted (1:400) in 1% BSA in PBS for 30 minutes at room temperature. Cells were washed with PBS three times between the above-mentioned steps. Subsequently, coverslips were mounted with ProLong™ Gold Antifade Reagent with DAPI (Invitrogen™, P36931). The slides were cured overnight and imaged on a Confocal Microscope (Zeiss LSM 710).

### TRAP staining

Tartrate-resistant acid phosphatase (TRAP) staining and TRAP activity in culture media were measured by TRAP staining kit with osteoplates (#, KT-651, Kamiya Biomedical Company), per manufacturer’s instructions. Briefly, RAW264.7 cells were treated with 50 ng/mL RANKL (#315-11, Peprotech) and with vehicle or 10 µM BTE-EN1 for 4 days, and media was replenished once after 2 days. Culture supernatants were collected for TRAP activity measurement and cells were fixed for TRAP staining.

### Cell growth inhibition assay

Three-day MTT growth inhibition assays were performed as previously described by us (*26*). Briefly, PC3 or C4-2B cells (5000 cell/well) were seeded into 96 well black/clear bottom plates, treated with 0, 0.2, 2, 10, 20, 100 nM of docetaxel for PC3 cells and 0, 0.2, 1, 1.5, 2, 10 nM of docetaxel for C4-2B cells, in the presence of either 10 µM BTE-EN1 or vehicle, and at 72 hours cell viability was assessed using the CyQUANT^TM^ MTT cell proliferation assay kit (Invitrogen #V13154) per manufactures instructions. The IC50 values were calculated using nonlinear regression analysis, as described by us (*62*). Data were log-transformed and fitted to a dose– response curve using the four-parameter logistic model of GraphPad Prism 10 software.

### Cell cycle analysis

A total of 3 x 10^5^ PC3 cells were seeded into 10 cm culture dishes, 24 hours later cells were treated in replicates of N=3, with 8 nM docetaxel with addition of vehicle or 10 µM BTE-EN1 for 24 or 72 hours. Cells were collected and filtered to obtain single-cell suspensions. Equal numbers of cells were fixed in 1 mL of cold 70% ethanol at 4 °C for 1 h, washed with PBS, and resuspended in 1 mL of Telford reagent (1 mM EDTA disodium salt, 2.5 U/mL RNase A, 75 µM Propidium Iodide, and 0.1% Triton X-100 in PBS), and then incubated at 4 °C overnight. Cells were analyzed by flow cytometry (FACSCalibur analyzer, BD Biosciences). Cell cycle analysis was performed using Modfit LT 4.0.5.

### Reverse transcription and qPCR

Cells were cultured in hormone-free conditions and qRT-PCR was performed as previously described by us (*26*). Briefly, LNCaP cells were precultured under hormone-free condition in charcoal-stripped phenol red-free RPMI1640 media for 48 hours and then treated with vehicle, 10 µM BTE-EN1, 1 nM R1881, 10 µM BTE-EN1 + 1 nM R1881, 1 nM R1881 + 10 µM enzalutamide (androgen receptor antagonist), and 10 µM BTE-EN1 + 1 nM R1881 + 10 µM enzalutamide for 24 hours. R1881 was added 1 hour after other treatments. RNA was isolated using RNeasy Plus Mini Kit (Qiagen) and reverse transcription was conducted with SuperScript III First-Strand Synthesis SuperMix (Thermo Fisher Scientific). The mRNA expression level of prostate-specific antigen (PSA) was measured by qPCR using QuantStudio^TM^ 6 Flex Real-Time PCR System (Applied Biosystems™). The primer/probe sets (TaqMan Gene Expression Assay) for PSA (prostate-specific antigen) and GPADH were Hs02576345_m1 and Hs99999905_m1, respectively. The data of qPCR was analyzed using double delta Ct method. Assays were performed in duplicate and were repeated 2 times.

### *In vivo* experiments

All animal studies adhered to the NIH Guide for the Care and Use of Laboratory Animals, and were treated under Institutional Animal Care and Use Committee–approved protocols by Oregon Health and Sciences University (OHSU) and University of Nebraska Medical Center (UNMC), complied with all federal, state, and local ethical regulations. Mice were maintained in a temperature- and humidity-controlled environment with a 12 h/12 h light/dark cycle and given soy-free food and water ad libitum. 6-to 8-week-old male athymic nude mice (Charles River) were used for IC and CA injection experiments. Body weight and IVIS imaging were conducted weekly until end points. CT imaging was performed at the end of the experiments to measure bone destruction. The toxicity of BTE-EN1 was evaluated by Calvert Laboratories, Inc. A comprehensive and detailed description of all experimental procedures can be found in the Supplementary Information.

### Statistical analysis

Data analysis was performed with Microsoft Excel and GraphPad Prism Statistical Software version 10. Statistical comparisons between two groups were evaluated with the unpaired two-tailed Student’s t-test. P-values ≤0.05 are considered significant. The values were presented as the means ± SEM, unless otherwise stated. The survival of mice was evaluated by the log-rank (Mantel-Cox) test.

## Data availability

The data that support the findings of this study are available from the corresponding author upon reasonable request.

## Acknowledgments

Financial support for this research was provided to R.B. by the United States Veterans Administration (IBX002842A) and the United States Department of Defense (W81XWH-15-1-0527). We would like to thank William Packwood for his assistance in performing the high-resolution CT imaging. We would like to thank Kanak Majumdar and Subrata Ghoshal for guidance related to chemical synthesis while at TCG Lifesciences Pvt. Ltd. in Kolkata, India. We acknowledge use of the UNMC Advanced Microscopy Core Facility, Echo Facility, In Vivo Imaging Core Facility and the University of Nebraska-Lincoln (UNL) Institutional Animal Care Program (IACP) for CT imaging. We additionally thank Bryan Hackfort of the UNMC Echo Facility and Cedric Wooledge of the UNL IACP for their assistance.

## Author contributions

Conceptualization and design: FQ, AP, RG, RB. Development of Methodology: FQ, AP, RG, RB. Investigation: FQ, AP, WL, RG, JK, KO, WC. Visualization: FQ AP, RG, PM, AN, RB. Funding acquisition: RB. Project administration: FQ, RG, RB. Supervision: FQ, RG, RB. Writing – original draft: FQ, AP, RG, RB. Writing – review & editing: FQ, AP, WL, RG, JK, KO, WC, PM, AN, RB.

