## Supplementary Information for "A New Class of Precision Therapeutics that Inhibit Prostate Cancer Mediated Bone Destruction"

**The PDF file includes:**

Supplementary Figures. 1 to 17

Supplementary Tables. 1 to 2

Supplementary Methods

- Chemistry
- Animal Models
- *In vivo* toxicity evaluation

Legend for Supplementary Movie 1

Legend for Supplementary Data file 1

Supplementary References

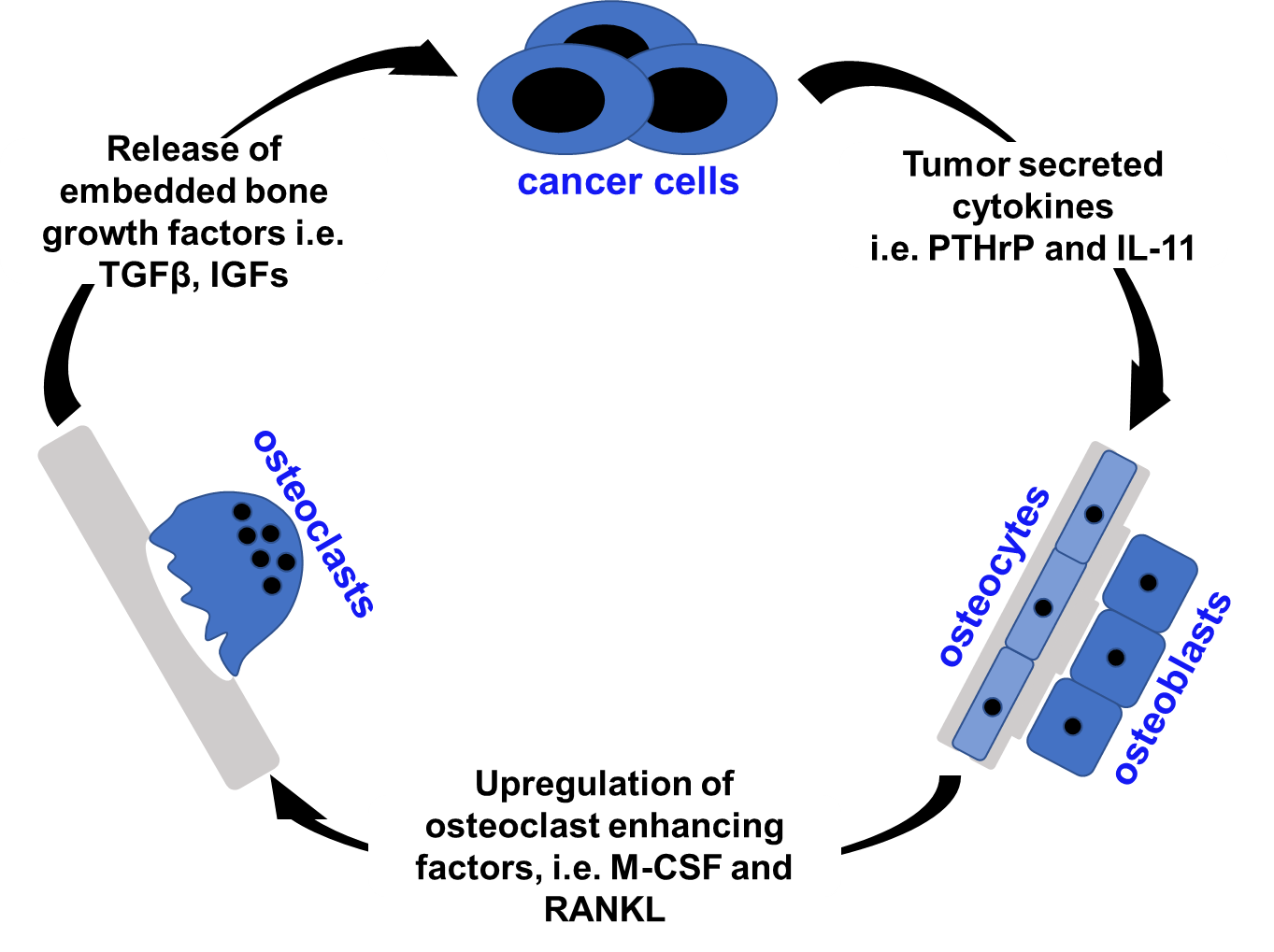

**Supplementary Figure 1.** **The vicious cycle.** Cancer cells secrete factors that stimulate osteoblasts and osteocytes to in turn increase osteoclast mediated matrix destruction, which releases cytokines from the matrix that stimulate cancer cell growth and movement. PTHrP = Parathyroid hormone-related protein; IL-11 = Interleukin 11; M-CSF = Macrophage colony-stimulating factor; RANKL = Receptor activator of nuclear factor-κB ligand; TGFβ = Transforming growth factor beta; IGF = Insulin-like growth factor.

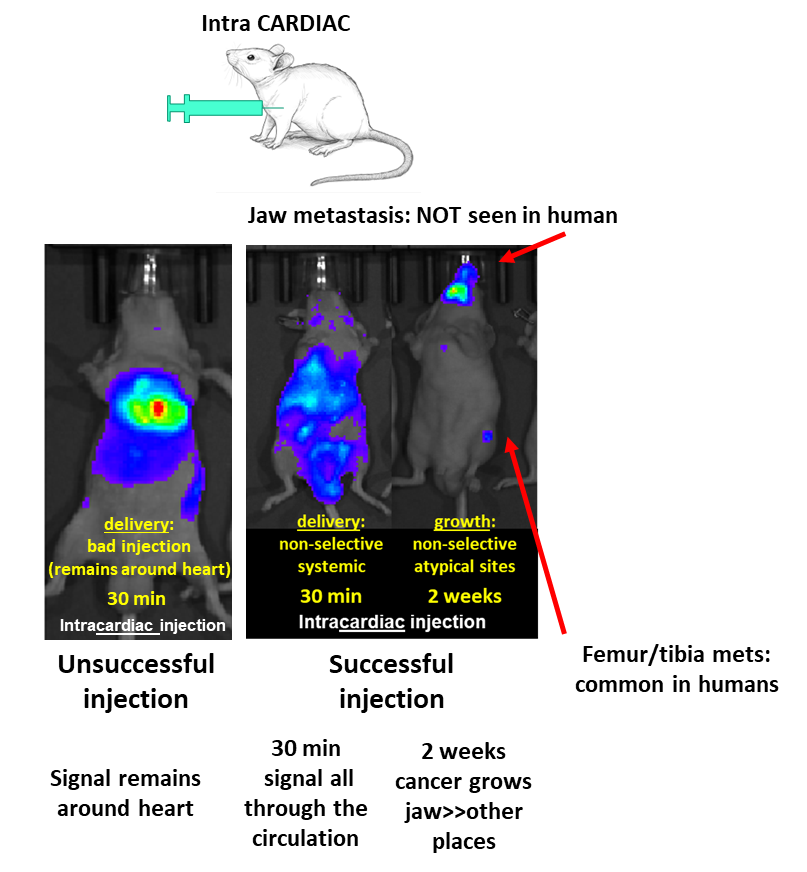

**Supplementary Figure 2**. **Intracardiac (IC) injection model.** 30 min after successful injection cells are evenly dispersed across the circulatory system. Over time cells implant in distant organs, typically bone, form metastatic deposits, and induce bone destruction, emulating the clinical scenario in humans. With a non-successful injection, typically due to puncture of the heart, cells collect in the pericardiac space (left panel).

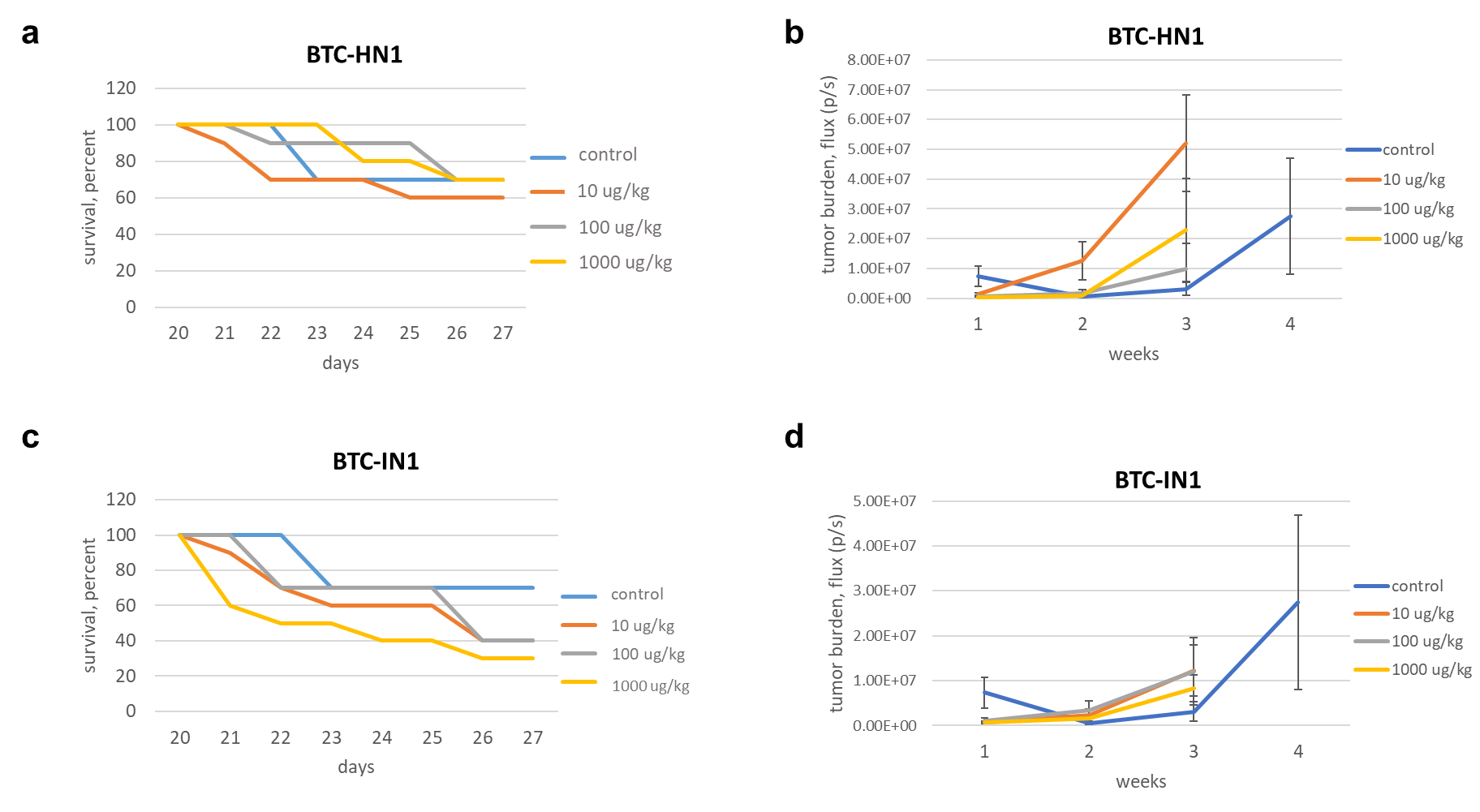

**Supplementary Figure 3**. **Performance of BTC-HN1 and BTC-IN1 compounds in the IC injection model.** Three days prior to IC injection of PC3-Luc cells, N=10 mice/cohort began weekly treatment with 10, 100 or 1000 µg/kg BTC-HN1 or BTC-IN1 intraperitoneally, with N=10 control mice receiving vehicle. (**a and c**) Survival curves of compounds BTC-HN1 and BTC-IN1, respectively. (**b and d**) Weekly mandibular IVIS imaging of compounds BTC-HN1 and BTC-IN1, respectively. IVIS imaging values for week 4 are not shown for treatment mice because rapid tumor growth led to death, thereby skewing selection for small tumors, and thus a spurious decrease in IVIS signal.

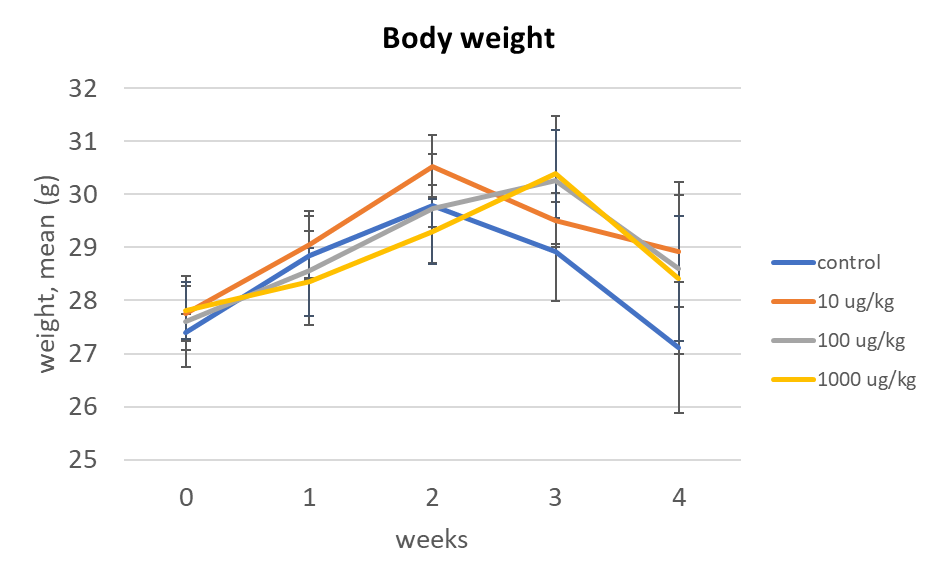

**Supplementary Figure 4**. **There was no evidence of toxicity with BTE-EN1.** As described in **Fig. 3a**, mice were dosed with weekly intraperitoneal injections of BTE-EN1 at 0 (vehicle), 10, 100 or 1000 µg/kg beginning 3 days before IC injection of PC3-Luc cells. Depicted above are the mean weights of mice. There was no statistical difference in weights between cohorts at any time point. Additional assessments of toxicity consisted of close observations of behavior and physical appearance, including skin color and turgor, activity level, body habitus, gait, and interaction with other animals. No differences between cohorts were noted.

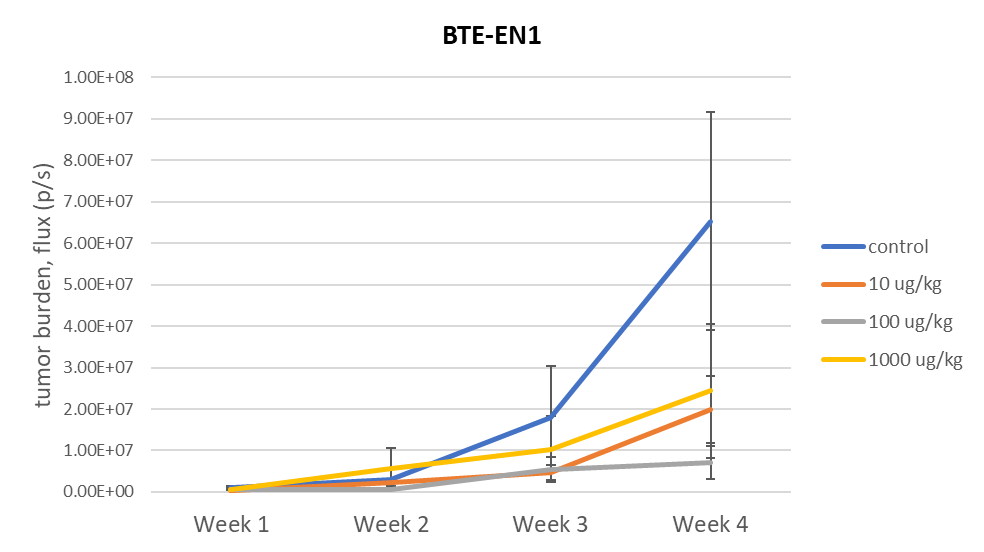

**Supplementary Figure 5**. **Weekly IVIS imaging of the mandible.** As described in **Fig. 3a**, mice were dosed with weekly intraperitoneal injections of BTE-EN1 at 0 (vehicle), 10, 100 or 1000 µg/kg beginning 3 days before IC injection of PC3-Luc cells. Depicted above are the weekly IVIS imaging results of mice. Weekly IVIS imaging of the mandible demonstrates the expected growth kinetics in controls, and diminished growth in treatment cohorts.

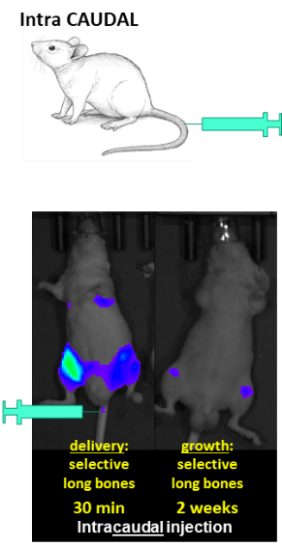

**Supplementary Figure 6. Intra-caudal artery injection model.** Representative 30 min and 2 weeks post-injection images demonstrate that PC3-Luc cells selectively traffic to clinically relevant sites, specifically to bones of the hind limbs.

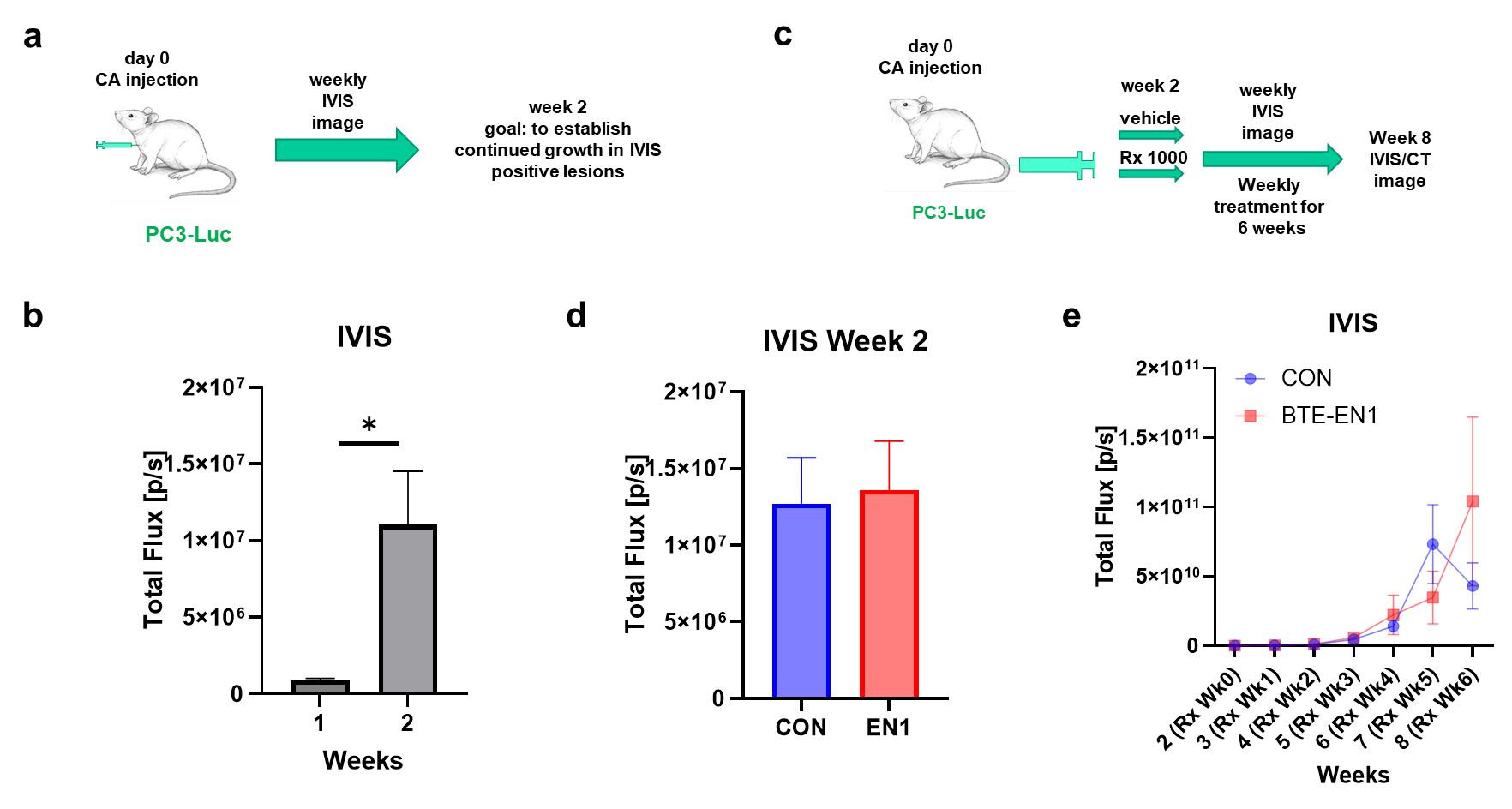

**Supplementary Figure 7.** **Weekly IVIS imaging of hind limbs.** (**a** and **b**) Tumor growth in hind limbs. (a) Experimental design of a CA injection model experiment. (b) Increase of IVIS signal from week 1 to week 2 in the hind limbs of mice, indicating tumor growth, N=9 mice. (**c** and **d**) Supplemental data for that described in **Fig. 4 a** **and c**. (c) Experimental design of the CA model. A total of 50 mice underwent CA injection on day 0, yielding 19 mice with established lesions in long bones of the legs, which were then randomized to BTE-EN1 treatment (N=10) or vehicle control (N=9). (d) IVIS signal at 2 weeks post injection, i.e., the time at which treatment began. Control and BTE-EN1 groups exhibited similar IVIS signal intensity. (**e**) Weekly IVIS imaging showed a progressive increase of tumor size in the hind limbs over 6 weeks of the treatment portion of the experiment, but no significant difference was observed. Note, the reverse of IVIS imaging values was associated with animals reaching to termination criteria in the control group (2 mice were sacrificed at week 8).

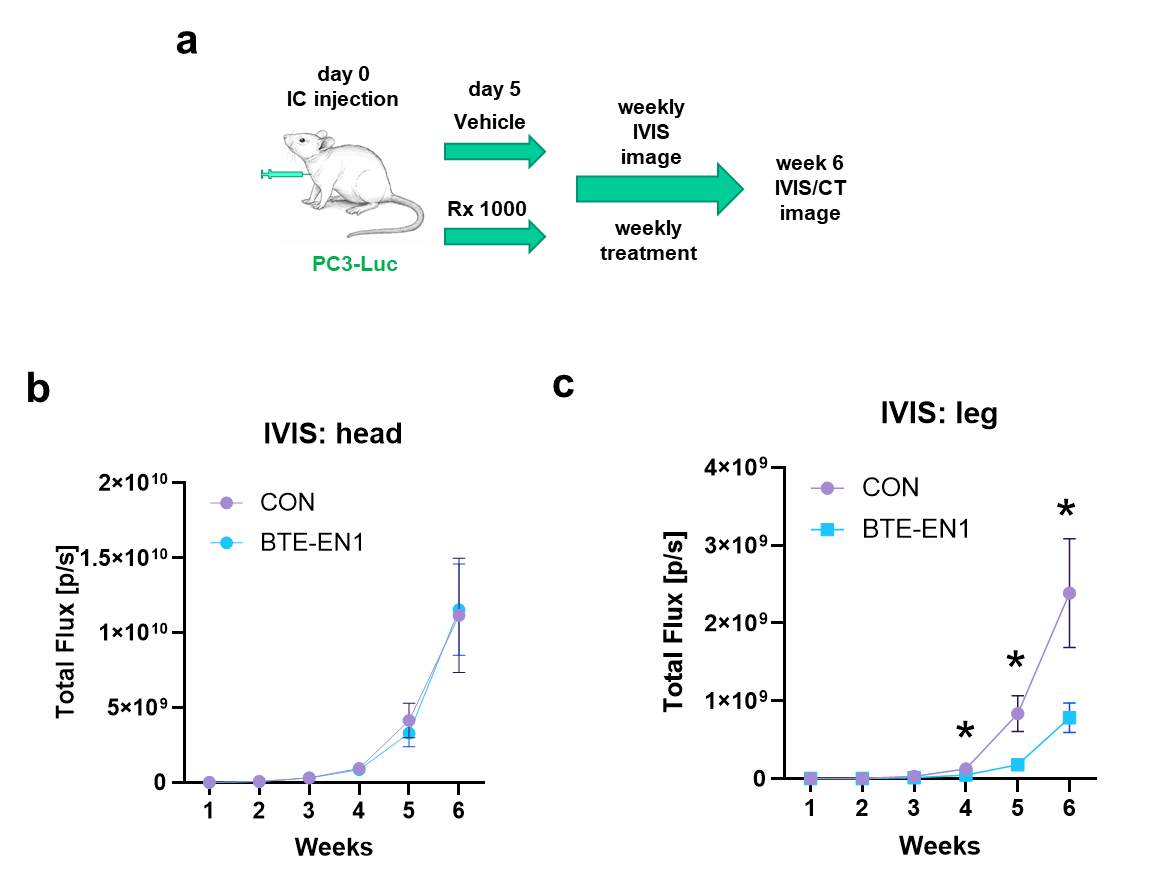

**Supplementary Figure 8. Weekly IVIS imaging of mouse head and legs.** (**a**) Five days post IC injection of PC3-Luc cells, mice were randomized to reflect equal numbers of mice with IVIS signal in the jawbone and leg bones in each cohort. mice began treatment with weekly 1000 µg/kg BTE-EN1 or vehicle, with N=24/cohort. (**b**) Weekly IVIS imaging of mouse head. (**c**) Weekly IVIS imaging of mouse legs. * Denotes Student’s t test P ≤ 0.05.

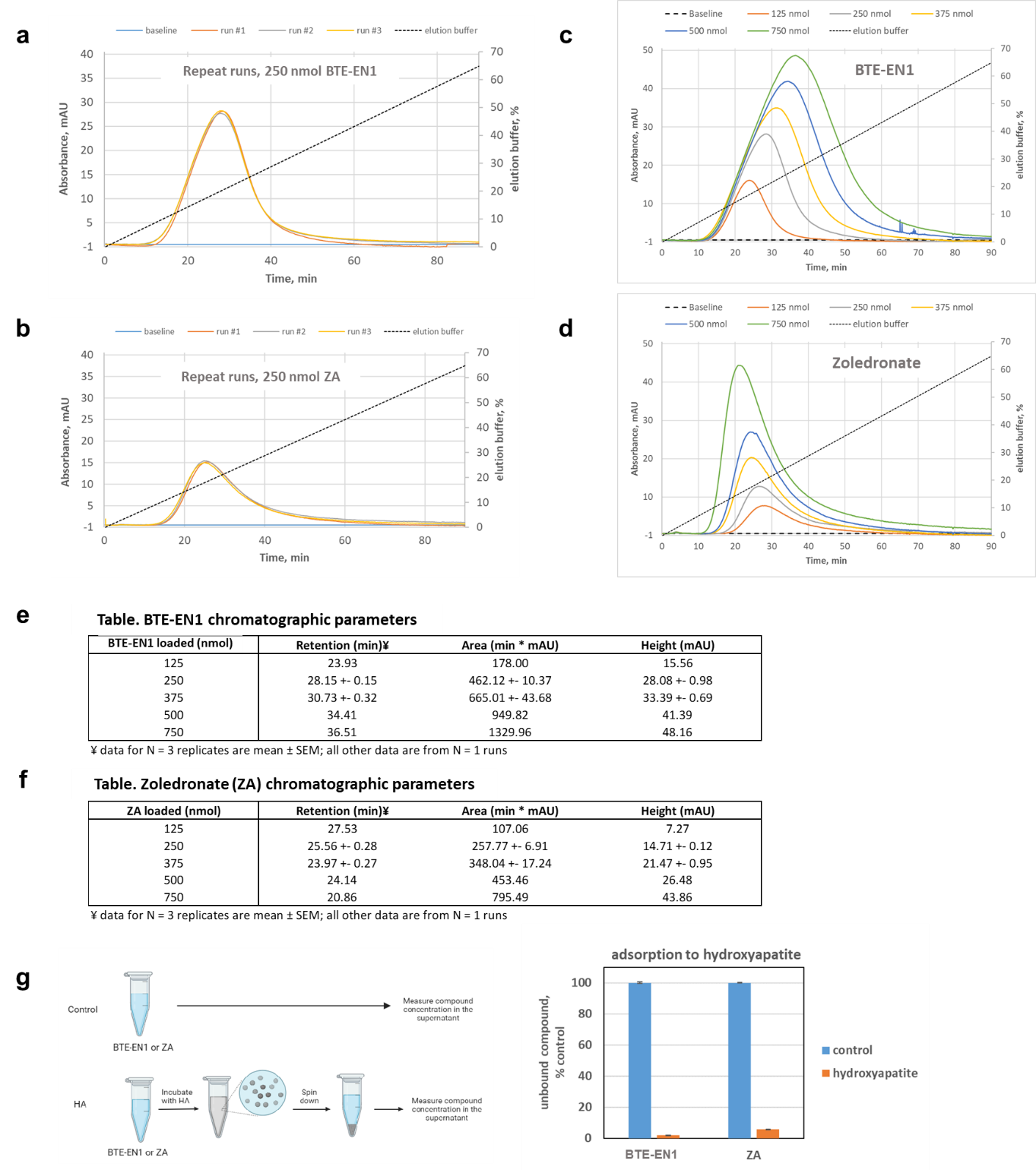

**Supplementary Figure 9. BTE-EN1 binds hydroxyapatite.** (**a-f**) The performance characteristics of hydroxyapatite column chromatographic assays. Assay repeatability was examined by loading equal amounts of BTE-EN1 (a) or zoledronate (b) onto hydroxyapatite columns and eluting under identical running conditions. Assay performance across a range of analytes was examined by loading differing amounts of BTE-EN1 (c) or zoledronate (d) onto hydroxyapatite columns and eluting under identical running conditions. The resultant chromatographic parameters for BTE-EN1 (e) and zoledronate (f) are presented in tabular form. Changes in chromatographic parameters as a function of amount loaded has been previously described (*6*) and needs to be taken into consideration when constructing standard curves to measure an unknown quantity. (**g**) BTE-EN1 and zoledronate exhibit a similar binding capacity to hydroxyapatite. Compounds were incubated with hydroxyapatite, excess washed off, remaining in solution quantified, and expressed as percent of control solution. HA: hydroxyapatite. Data are the mean ± SEM of N=3 separate experiments.

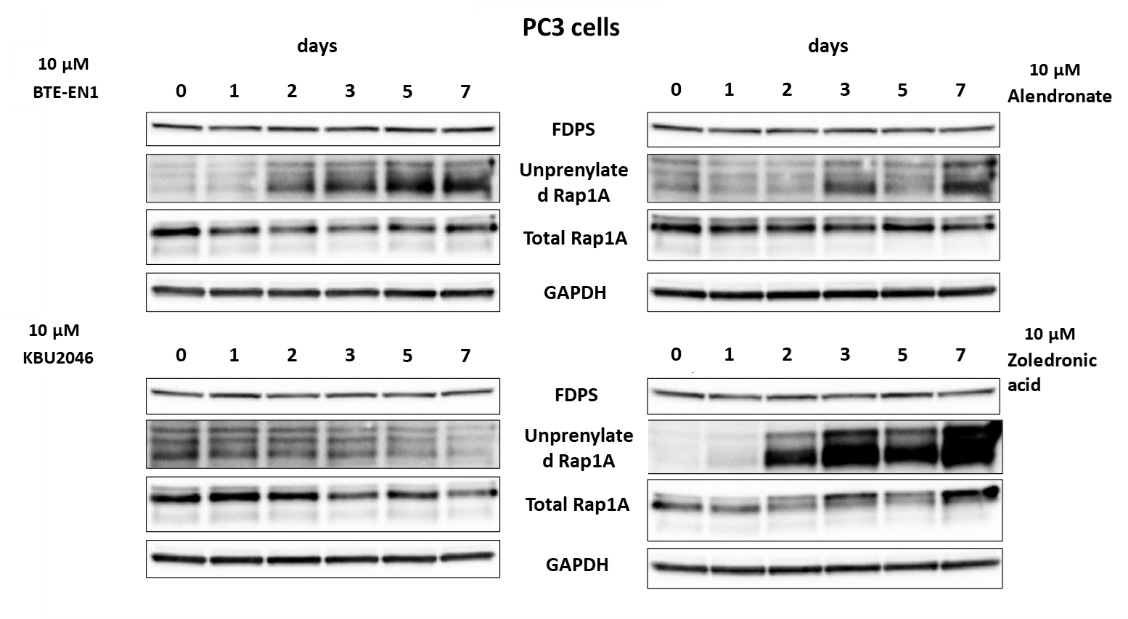

**Supplementary Figure 10. BTE-EN1 inhibits protein prenylation in a time-dependent fashion.** PC3 cells were treated with denoted drug at 10 µM for 1, 2, 3, 5, and 7 days, or with vehicle (0), and probed for the indicated proteins by Western blot.

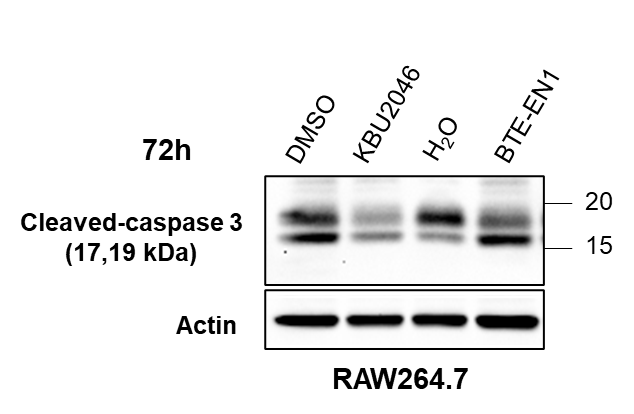

**Supplementary Figure 11. BTE-EN1 but not KBU2046, induces apoptosis in RAW264.7 cells.** After pre-treatment of RAW264.7 cells with 50 ng/mL RANKL for 2 days, they were then treated with 10 µM KBU2046, 10 µM BTE-EN1 or respective vehicle controls for 72 hours, along with continued RANKL treatment. Resultant Western blot depicts cleaved-caspase 3. Note that at 5-7 days post RANKL treatment, cells are terminally differentiated and are undergoing spontaneous apoptosis, as evident in DMSO and water vehicle controls. KBU2046 inhibits the differentiation of osteoclasts and therefore delays apoptotic process compared to the vehicle control which is demonstrated by reduced activation of cleaved-caspase 3. In contrast, compared to water vehicle control, BTE-EN1 increases cleaved-caspase 3 generation.

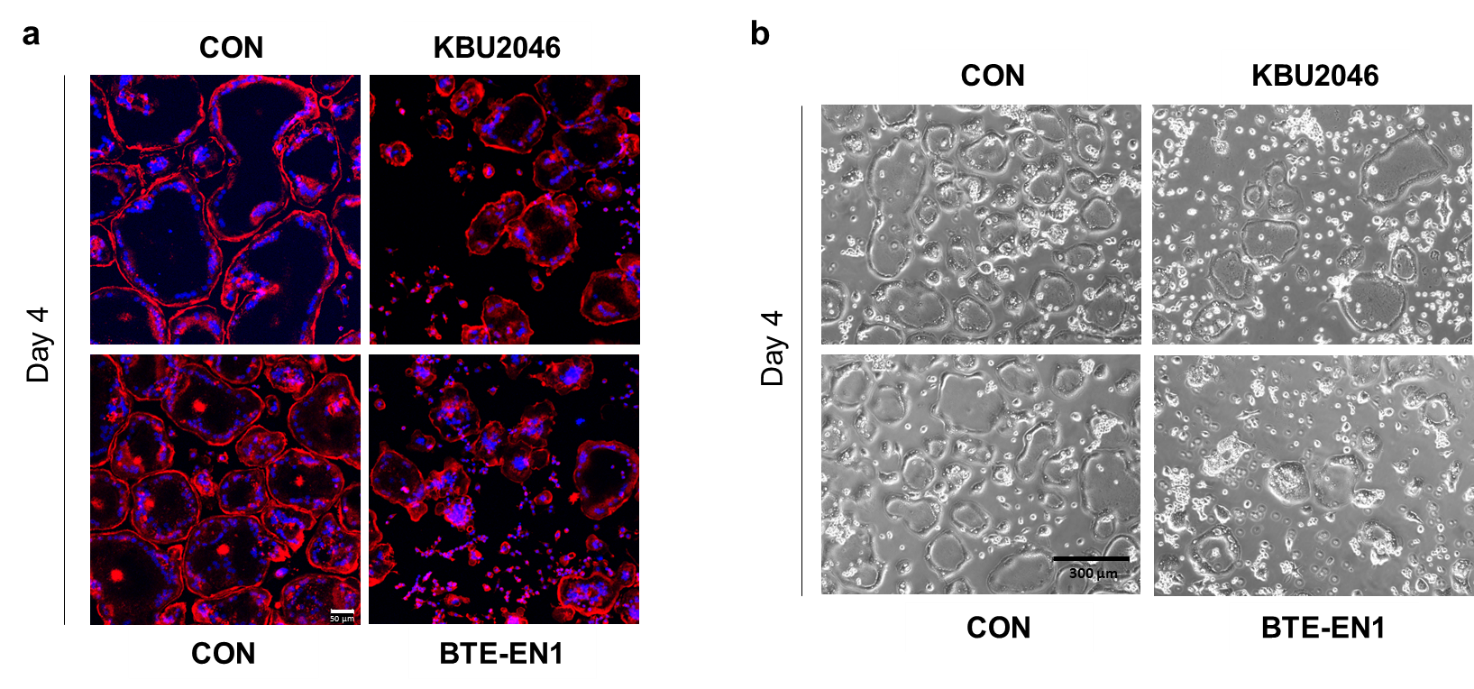

**Supplementary Figure 12.** **BTE-EN1 Inhibits actin ring formation. Fig. 6** depicts RAW264.7 cells at 6 days of maturation, just prior to their terminal/death phase. This supplemental figure depicts findings at an earlier time point. RAW264.7 cells were treated with 50 ng/mL RANKL and with 10 µM KBU2046, 10 µM BTE-EN1, or vehicle controls for 4 days, and stained for actin (Rhodamine Phalloidin, red) and nuclei (DAPI, blue). Representative immunofluorescent images of cells were stained for actin (Rhodamine Phalloidin, red) and nuclei (DAPI, blue) are shown in (**a**), scale bar=50 µm, while light microscopic images are shown in (**b**), scale bar=300 µm.

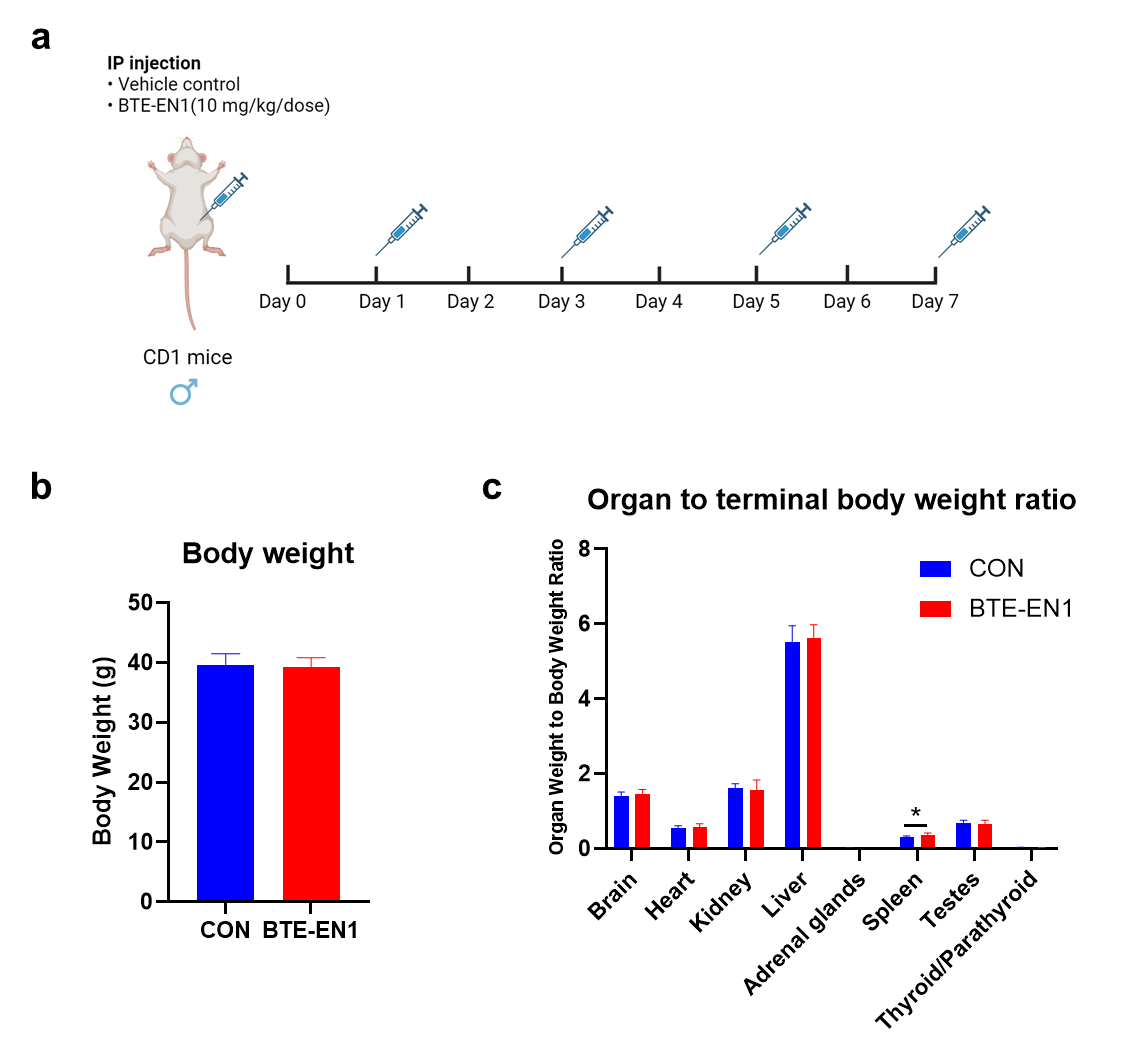

**Supplementary Figure 13. *In vivo* toxicity evaluation of BTE-EN1.** Toxicological analysis was performed as described in Methods. (**a**) Experimental design of 7-Day Repeat-Dose Study. BTE-EN1 (10,000 mcg/kg/dose) and vehicle control were administered by intraperitoneal injection to groups of 6 male 8-week-old CD1 mice on Days 1, 3, 5, and 7. (**b**) Body weights (Mean ± SD) at Day 7. (**c**) Organ weight to terminal body weight ratio. * Denotes P ≤ 0.05 (Dunnett LSD Test). Note, the increased spleen size was associated with intraperitoneal administration of high doses of test article, i.e., BTE-EN1. The final report from Calvert Laboratories stated, “All BTE-EN1-related findings were considered non-adverse.”

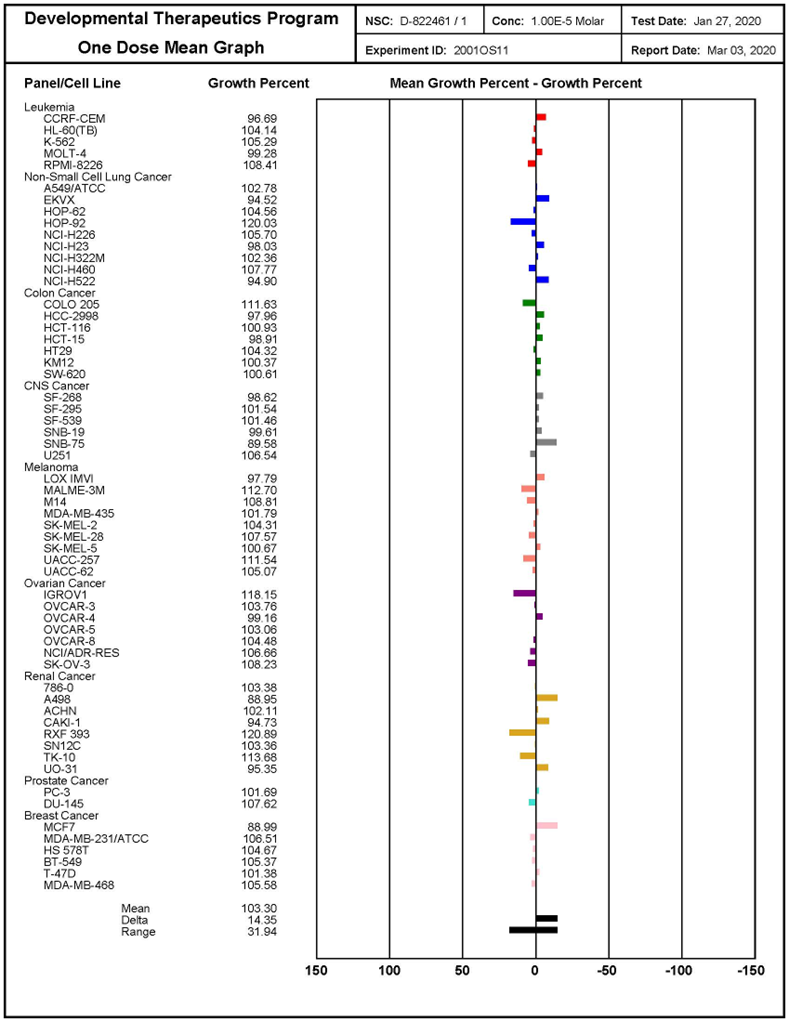

**Supplementary Figure 14**. **BTE-EN1 is not toxic to the cells in the NCI-60 cell line panel.** BTE-EN1 was tested by NCI in the NCI-60 cell line screen. The associated COMPARE plot is depicted.

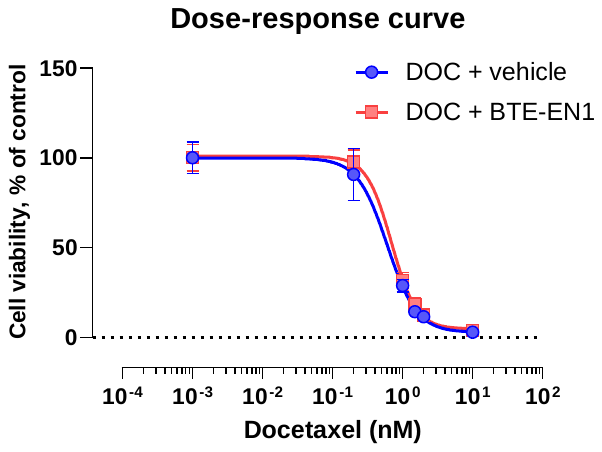

**Supplementary Figure 15**. **BTE-EN1 does not interfere with cytotoxic agent effect in PCa cell line C4-2B.** C4-2B cells were treated with docetaxel (DOC) in the presence of 10 µM BTE-EN1 or vehicle for 3 days, and cell viability was measured by MTT assay. IC50 [DOC + vehicle]=1.186 ± 0.28 nM, IC50 [DOC + BTE-EN1]=1.699 ± 0.02 nM (mean ± SEM), calculated using N=4 replicates. Student’s t-test. Data are the mean ± SD of single experiments conducted in replicates of N=8.

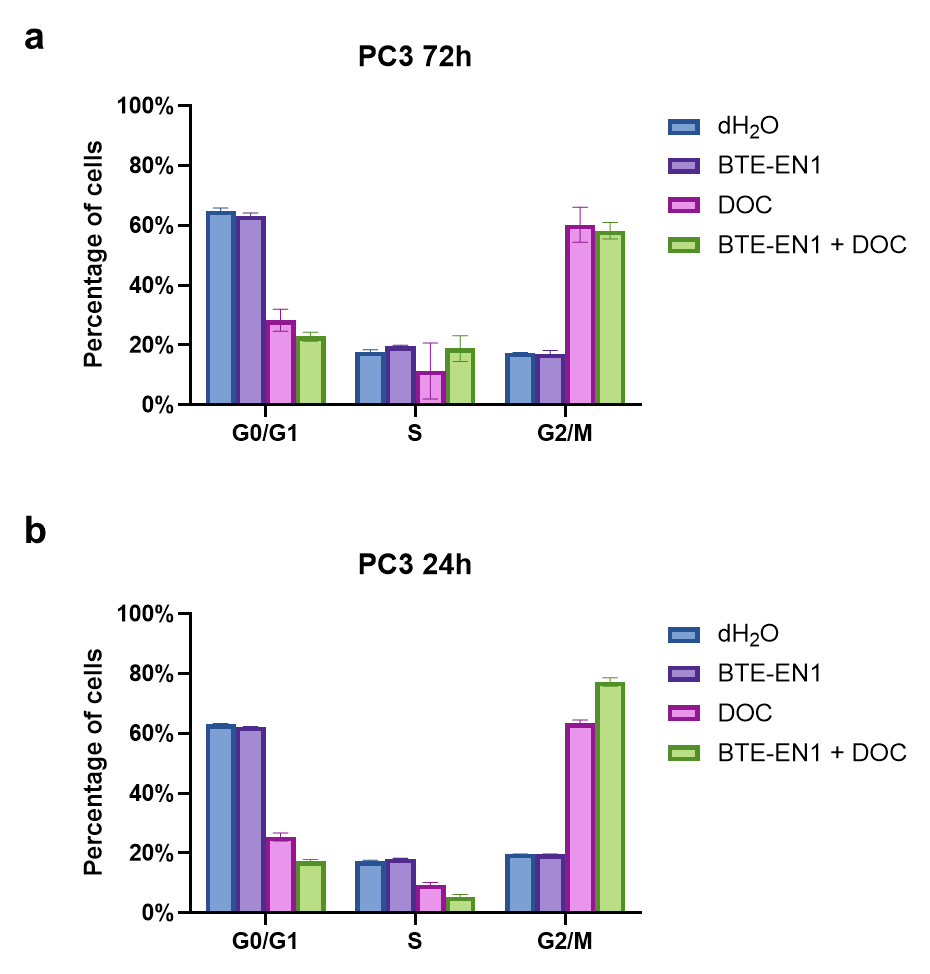

**Supplementary Figure 16. BTE-EN1 does not interfere with docetaxel’s effect on cell cycle inhibition in PC3 cells.** Cell cycle analysis of PC3 cells. PC3 cells were treated with 10 µM BTE-EN1, 8 nM docetaxel, the combination or vehicle for 72 hours (**a**) and 24 hours (**b**). Data are the mean ± SEM (N=3); data were analyzed by Student’s t-test.

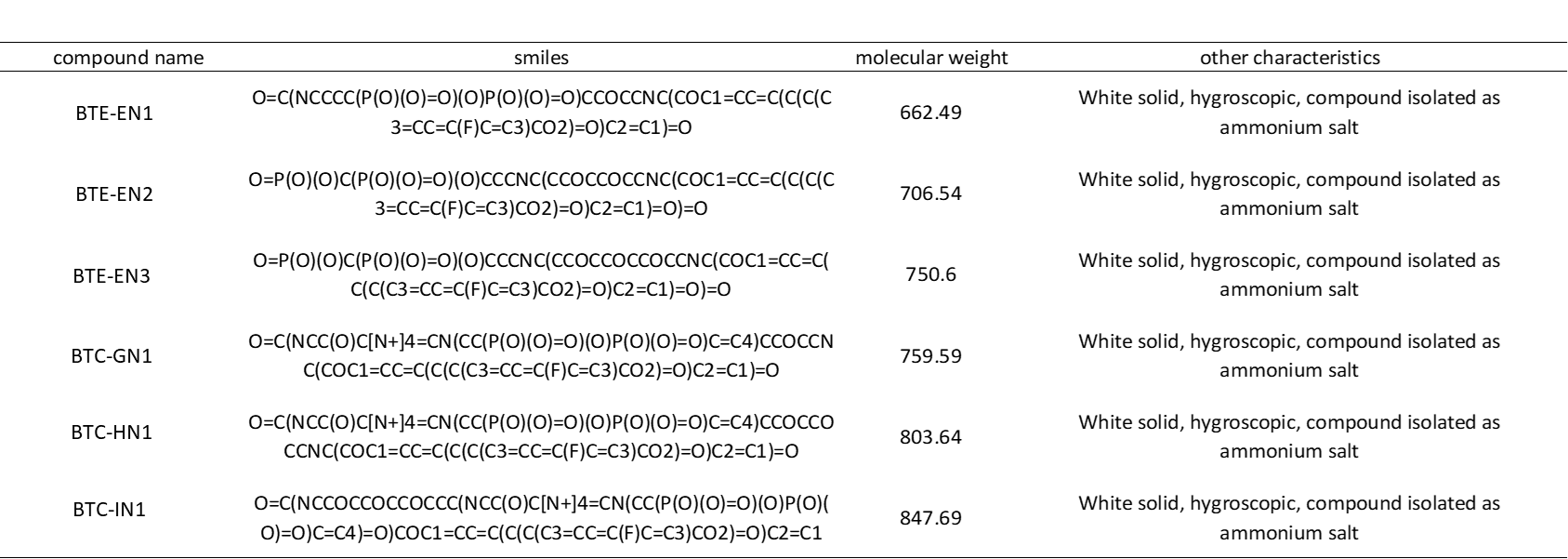

**Supplementary Table 1**. **Chemical characteristics of synthesized compounds.**

| Table 2. Tissues evaluated for toxicity | |
| --- | --- |
| Adrenal glands | Aorta |
| Brain | Epididymides |
| Esophagus | Eye |
| Femur w/ articular surface | Gallbladder |
| Heart | Cecum |
| Colon | Rectum |
| Duodenum | Jejunum |
| Ileum | Kidneys |
| Lacrimal gland(s) | Larynx |
| Liver | Lung w/ mainstem bronchus |
| Lymph node, mandibular | Lymph node, mesenteric |
| Mammary gland | Nerve - sciatic |
| Pancreas | Pituitary gland |
| Prostate gland | Salivary glands |
| Seminal vesicles | Skeletal muscle (thigh) |
| Skin | Spinal cord, cervical |
| Spinal cord, lumbar | Spinal cord, midthoracic |
| Spleen | Sternum w/ bone marrow |
| Stomach | Testes |
| Thymus | Thyroid/Parathyroid |
| Tongue | Trachea |
| Urinary bladder |  |

**Supplementary Table 2. Tissues analyzed at necropsy.** Male 8-week-old CD1 mice were treated with BTE-EN1 at the dose and schedule described in **supplementary** **fig. 13**. At day 8 mice underwent necropsy, and the above organs were examined at the microscopic and macroscopic levels.

**Supplementary Methods**

**Chemistry**

All chemicals and solvents were commercially purchased and used without further purification unless otherwise specified. ^1^H Nuclear magnetic resonance (NMR) (400 MHz) spectra were recorded in MeOD-*d*4 on a Bruker-400 spectrometer, with tetramethyl silane (Me₄Si) as the internal standard. Chemical shifts are reported in δ values (ppm), and the spectral data are presented in the format: δ chemical shift (multiplicity, J values in Hz, integration). The following abbreviations are used: s = singlet, d = doublet, t = triplet, and m = multiplet. Mass spectrometry (MS) analysis was conducted using ultrahigh-pressure liquid chromatography (UPLC) mass spectrometer.

***General Synthetic route for the synthesis of Dual Acting Bone Defenders (DABDs)***

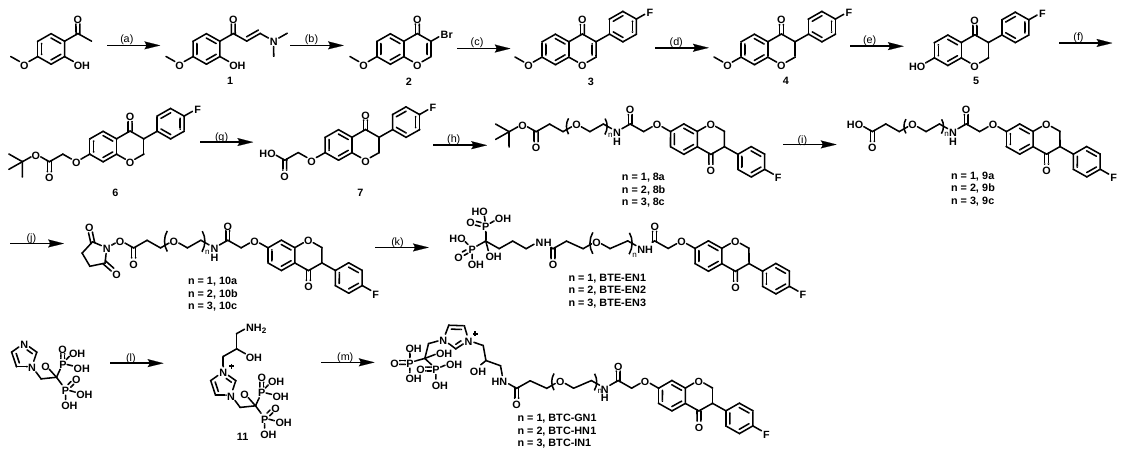

Reagents and conditions: (a) DMF-DMA, DMF, 1h, 86.2%; (b) NBS, DCM, 0-5 °C, 0.5 h, 76.4%; (c) 4-Fluoro phenyl boronic acid, Pd(PPh3)4, aq.Na2CO3, dioxane, 80 °C, 6h, 63.1%; (d) L-Selectride, THF, -78 °C, 1h, 62.1%; (e) AlCl3, 1-dodecanthiol, DCM, rt, 8h, 74.6%; (f) tert. butyl bromo acetate, DIPEA, DMF, rt, 16h, 83.1%; (g) TFA, DCM, rt, 0.5 h, 80.7%; (h) PEG-Amines, TBTU, DIPEA, DMF, rt, 16h, 65.4-75.9%; (i) TFA, DCM, rt, 2h, 84.1-94.0%; (j) N-hydroxy succinimide, EDC, DMF, rt, 16h; (k) alendronic acid, KHCO3, DMF, rt, 16 h, 16.0-19.0%; (l) Oxiranylmethyl-carbamic acid tert-butyl ester, Na2CO3, MeOH, 50 °C, 3days followed by TFA, rt, 4h; (m) 10, KHCO3, DMF, rt, 16 h, 1.8-5.8%.

***Chemical synthesis***

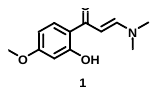

**Procedure for the Synthesis of (E)-3-Dimethylamino-1-(2-hydroxy-4-methoxy-phenyl)-propenone (1):** To a stirred solution of 4-methoxy-2-hydroxyacetophenone (602.4 mmol, 100 g) in dimethylformamide (DMF) (1320 mL) was added DMF-DMA (1807.23 mmol, 287.8 g) and resultant reaction mixture was allowed to stir at 80 °C for 1 h. After completion [confirmed by thin layer chromatography (TLC) analysis, 20% EtOAc-hexane, Rf-0.2] reaction mixture was quenched with cold water [1500 mL] and resultant light orange precipitate was filtered through glass sintered, dried under vacuum to afford (E)-3-Dimethylamino-1-(2-hydroxy-4-methoxy-phenyl)-propenone (115 g, 86.2%) as light orange solid.

**Analytical data for (E)-3-Dimethylamino-1-(2-hydroxy-4-methoxy-phenyl)-propenone (1)**: ^1^H NMR (400 MHz, CDCl_3_) δ 14.45 (s, 1H), 7.80 (d, *J* = 12.1 Hz, 1H), 7.58 (d, *J* = 8.0 Hz, 1H), 6.39 (d, *J* = 2.0 Hz, 1H), 6.35 (dd, *J* = 8.8, 2.2 Hz, 1H), 5.65 (d, *J* = 12.1 Hz, 1H), 3.8 (s, 3H), 3.15 (s, 3H), 2.93 (s, 1H). Ultra performance liquid chromatography/mass spectrometry (UPLCMS): Mass calculated for C_12_H_16_NO_3_, [M+H]^+^, 222. Found 222.

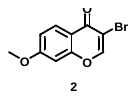

**Procedure for the Synthesis of 3-Bromo-7-methoxy-chromen-4-one (2):** To a stirred solution of (E)-3-Dimethylamino-1-(2-hydroxy-4-methoxy-phenyl)-propenone **(1)** (496.6 mmol, 110 g) in dichloromethane (DCM) (2200 mL) was added N-Bromosuccinimide (NBS) (521.44 mmol, 92.8 g) maintaining the external temperature under 0-5 °C and stirred for 30 minutes at same temperature. Then the reaction mixture was quenched with water [2000 mL]. Organic layer was separated, concentrated under reduced pressure. Resultant crude was triturated with Pentane [500 mL] to afford 3-Bromo-7-methoxy-chromen-4-one (96.8 g, 76.4%) as off white solid.

**Analytical Data for 3-Bromo-7-methoxy-chromen-4-one (2)**: ^1^H NMR (400 MHz, CDCl_3_) δ 8.13-8.15 (m, 2H), 6.98 (dd, *J* = 8.8, 1.6 Hz, 1H), 6.83 (d, *J* = 1.4 Hz, 1H), 3.89 (s, 3H) UPLCMS: Mass calculated for C_10_H_8_BrO_3_, [M+H]^+^, 255. Found 255.

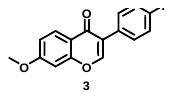

**Procedure for the Synthesis of 3-(4-Fluoro-phenyl)-7-methoxy-chromen-4-one (3):** To a well stirred solution of 3-Bromo-7-methoxy-chromen-4-one **(2)** (351.85 mmol, 95 g) in Dioxane (1520 mL) was added 4-Fluoro Phenyl boronic acid (351.85 mmol, 49.26 g) and aqueous solution of sodium carbonate (703.7 mmol, 74.59 g in 980 mL water). Then the heterogeneous reaction mixture was purged with N_2_ for 10 minutes followed by addition of Pd (PPh_3_)_4_ (17.6 mmol, 20.3 g) and allowed to stir at 80 °C for 6 h. After completion reaction mixture was brought to room temperature and partitioned with EtOAc (1500 mL). Organic layer was separated, washed with brine (500 mL) and concentrated under reduced pressure. Resultant crude was triturated with Et_2_O (200 mL) to afford 3-(4-Fluoro-phenyl)-7-methoxy-chromen-4-one (60 g, 63.1 %) as off white solid.

**Analytical Data for 3-(4-Fluoro-phenyl)-7-methoxy-chromen-4-one (3)**: ^1^H NMR (400 MHz, CDCl_3_) δ 8.18 (d, *J* = 8.9 Hz, 1H), 7.92 (s, 1H), 7.51 – 7.54 (m, 2H), 7.09 (t, *J* = 8.6 Hz, 2H), 6.98 (d, *J* = 8.6 Hz, 1H), 6.85 (s, 1H), 3.91 (s, 3H) UPLCMS: Mass calculated for C_16_H_12_FO_3_, [M+H]^+^, 271. Found 271.

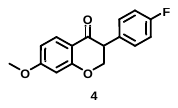

**Procedure for the Synthesis of 3-(4-Fluoro-phenyl)-7-methoxy-chroman-4-one (4):** Solution of 3-(4-Fluoro-phenyl)-7-methoxy-chromen-4-one **(3)** (111.04 mmol, 30 g) in THF (750 mL) was allowed to cooled to -78 °C and 1(M) solution of L-Selectride (240 mmol, 240 mL ) was added over a period of 30-40 minutes via a dropping funnel and allowed to stir at same temperature for additional 30 minutes. Then the reaction mixture was slowly quenched with MeOH (50 mL) and was brought to room temperature. Reaction mixture was poured into ice cold water (500 mL) and pH was adjusted to ~6-7 using 1(N) aqueous HCl. Then reaction mixture was extracted with EtOAc (1000 mL), organic layer was separated, washed with brine (200 mL) and concentrated under reduced pressure. Resultant crude was purified by column chromatography under gradient elution of [2-5 % EtOAc-hexane] to provide 3-(4-Fluoro-phenyl)-7-methoxy-chroman-4-one (18.5 g, 62.12%) as off white solid. **Analytical data of 3-(4-Fluoro-phenyl)-7-methoxy-chroman-4-one (4)**: (18.5 g, 62.12%) ^1^H NMR (400 MHz, DMSO-*d*_6_) δ 7.73 (d, *J* = 8.4 Hz, 1H), 7.29 – 7.33 (m, 2H), 7.15 (t, *J* = 8.4 Hz, 2H), 6.66 (dd, *J* = 8.8, 2.0 Hz, 1H), 6.59 (d, *J* = 2.0 Hz, 1H), 4.61 – 4.73 (m, 2H), 4.13 – 4.16 (m, 1H), 3.83 (s, 3H) UPLCMS: Mass calculated for C_16_H_14_FO_3_, [M+H]^+^, 273. Found 273.

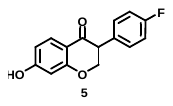

**Procedure for the Synthesis of 3-(4-Fluoro-phenyl)-7-hydroxy-chroman-4-one (5):** AlCl_3_ (352.94 mmol, 46.94 g) and 1-dodecanthiol (176.47 mmol, 42.43 mL) were allowed to stir with DCM (100 mL) for 30 minutes at 0-5 °C. Then solution of 3-(4-Fluoro-phenyl)-7-methoxy-chroman-4-one **(4)** (44.11 mmol, 12 g) in DCM (60 mL) was slowly added to the reaction mixture and stirred at room temperature for 8 h. After completion reaction mixture was quenched with water (200 mL) then pH was adjusted to ~6-7 using 1(N) aqueous HCl and the resultant reaction mixture was extracted with EtOAc (500 mL). Organic layer was separated, washed with brine, dried over sodium sulphate and concentrated under reduced pressure to afford light brown solid. The crude compound was further triturated with n-Pentane (200 mL) to provide 3-(4-Fluoro-phenyl)-7-hydroxy-chroman-4-one (8.5 g, 74.6%) as light brown solid.

**Analytical data of 3-(4-Fluoro-phenyl)-7-hydroxy-chroman-4-one (5):** ^1^H NMR (400 MHz, DMSO-*d*_6_) δ 10.63 (s, 1H), 7.65 (d, *J* = 8.6 Hz, 1H), 7.28 (t, *J* = 5.6 Hz, 2H), 7.14 (t, *J* = 8.6 Hz, 2H), 6.51 (d, *J* = 8.6 Hz, 1H), 6.35 (s, 1H), 4.56 – 4.66 (m, 2H), 4.08 – 4.11 (m, 1H) UPLCMS: Mass calculated for C_15_H_12_FO_3_, [M+H]^+^, 259. Found 259.

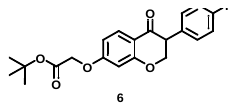

**Procedure for the Synthesis of [3-(4-Fluoro-phenyl)-4-oxo-chroman-7-yloxy]-acetic acid tert-butyl ester (6)**: To a stirred solution of 3-(4-Fluoro-phenyl)-7-hydroxy-chroman-4-one **(5)** (19.38 mmol, 5 g) in DMF (50 mL) was added tert butyl bromo acetate (29.07 mmol, 5.64 g), DIPEA (58.14 mmol, 7.5 g) and resultant reaction mixture was allowed to stir at room temperature for 16 h. Then reaction mixture was partitioned between EtOAc (500 mL) and water (300 mL). Organic layer was separated, drier over sodium sulphate and concentrated under reduced pressure, to afford [3-(4-Fluoro-phenyl)-4-oxo-chroman-7-yloxy]-acetic acid tert-butyl ester (6 g, 83.14%) as off white solid.

**Analytical data of [3-(4-Fluoro-phenyl)-4-oxo-chroman-7-yloxy]-acetic acid tert-butyl ester (6):** ^1^H NMR (400 MHz, DMSO) δ 7.73 (d, *J* = 8.8 Hz, 1H), 7.29 – 7.33 (m, 2H), 7.15 (t, *J* = 8.6 Hz, 2H), 6.66 (dd, *J* = 8.6, 1.6 Hz, 1H), 6.54 (s, 1H), 4.78 (s, 2H), 4.61 – 4.7 (m, 2H), 4.15 – 4.19 (m, 1H), 1.42 (s, 9H) UPLCMS: Mass calculated for C_21_H_22_FO_5_, [M+H]^+^, 373. Found 373.

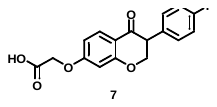

**Procedure for the Synthesis of [3-(4-Fluoro-phenyl)-4-oxo-chroman-7-yloxy]-acetic acid (7)**: To a well stirred solution of [3-(4-Fluoro-phenyl)-4-oxo-chroman-7-yloxy]-acetic acid tert-butyl ester **(6)** (6 g, 16.13 mmol) in DCM (80 mL) was added TFA (10 mL) drop-wise maintaining the external temperature at 0-5 °C and resultant reaction mixture was allowed to stir at same temperature for 30 minutes. After completion [Monitored with TLC] reaction mixture was concentrated under reduced pressure and azeotroped with toluene to provide [3-(4-Fluoro-phenyl)-4-oxo-chroman-7-yloxy]-acetic acid (4.12 g, 80.7%) as off white solid.

**Analytical data of [3-(4-Fluoro-phenyl)-4-oxo-chroman-7-yloxy]-acetic acid (7)**: ^1^H NMR (400 MHz, DMSO-*d*_6_) δ 13.13 (s, 1H), 7.73 (d, *J* = 8.8 Hz, 1H), 7.29 – 7.33 (m, 2H), 7.15 (t, *J* = 8.8 Hz, 2H), 6.67 (dd, *J* = 8.8, 2 Hz, 1H), 6.55 (d, *J* = 1.8 Hz, 1H), 4.8 (s 2H), 4.61 – 4.73 (m, 2H), 4.14 – 4.18 (m, 1H) UPLCMS: Mass calculated for C_17_H_14_FO_5_, [M+H]^+^, 317. Found 317.

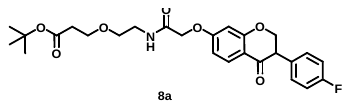

**Procedure for the Synthesis of 3-(2-{2-[3-(4-Fluoro-phenyl)-4-oxo-chroman-7-yloxy]-acetylamino}-ethoxy)-propionic acid tert-butyl ester (8a)**: Solution of [3-(4-Fluoro-phenyl)-4-oxo-chroman-7-yloxy]-acetic acid (3.16 mmol, 1 g), TBTU (4.74 mmol, 1.52 g) and DIPEA (9.49 mmol, 1.22 g) in DMF (15 mL) was allowed to stir at room temperature for 10 minutes. Followed by addition of PEG 1-Amine (3.16 mmol, 0.6 g) and the resultant reaction mixture was allowed to stir at room temperature for 16 h. Then the reaction mixture was partitioned between EtOAc (300 mL) and water (200 mL). Organic layer was separated, dried over sodium sulphate and concentrated under reduced pressure. Crude residue thus obtained was purified by column chromatography using silica gel 100-200 mesh under gradient elution of 40-50 % EtOAc-hexane to afford 3-(2-{2-[3-(4-Fluoro-phenyl)-4-oxo-chroman-7-yloxy]-acetylamino}-ethoxy)-propionic acid tert-butyl ester (1.1 g, 65.39%) as pale brown liquid.

**Analytical data of 3-(2-{2-[3-(4-Fluoro-phenyl)-4-oxo-chroman-7-yloxy]-acetylamino}-ethoxy)-propionic acid tert-butyl ester (8a)**: ^1^H NMR (400 MHz, DMSO-*d*_6_) δ 8.11 (t, *J* = 5.6 Hz, 1H), 7.74 (d, *J* = 8.8 Hz, 1H), 7.29 – 7.33 (m, 2H), 7.15 (t, *J* = 8.8 Hz, 2H), 6.7 (dd, *J* = 8.8, 2.2 Hz, 1H), 6.57 (d, *J* = 2.2 Hz, 1H), 4.61 – 4.73 (m, 2H), 4.58 (s, 2H), 4.14 – 4.18 (m, 1H), 3.56 (t, *J* = 6.2 Hz, 2H) , 3.41 (t, *J* = 5.8 Hz, 2H), 3.25 – 3.31 (m, 2H), 2.4 (t, *J* = 6.2 Hz, 2H), 1.38 (s, 9H) UPLCMS: Mass calculated for C_26_H_31_FNO_7_, [M+H]^+^, 488. Found 488.

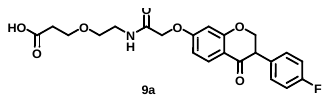

**Procedure for the Synthesis of 3-(2-{2-[3-(4-Fluoro-phenyl)-4-oxo-chroman-7-yloxy]-acetylamino}-ethoxy)-propionic acid (9a):** Solution of 3-(2-{2-[3-(4-Fluoro-phenyl)-4-oxo-chroman-7-yloxy]-acetylamino}-ethoxy)-propionic acid tert-butyl ester **(8a)** (1.1 g, 2.25 mmol) in DCM (10 mL) was treated with TFA (4 mL) and resultant reaction mixture was stirred at room temperature for 2 h. Then reaction mixture was concentrated under reduced pressure and triturated with pentane (50 mL) to afford 3-(2-{2-[3-(4-Fluoro-phenyl)-4-oxo-chroman-7-yloxy]-acetylamino}-ethoxy)-propionic acid (900 mg, 84.1%) as off white solid.

**Analytical data of 3-(2-{2-[3-(4-Fluoro-phenyl)-4-oxo-chroman-7-yloxy]-acetylamino}-ethoxy)-propionic acid (9a)**: ^1^H NMR (400 MHz, DMSO-*d*_6_) δ 12.2 (brs, 1H), 8.13 (t, *J* = 5.7 Hz, 1H), 7.74 (d, *J* = 8.8 Hz, 1H), 7.29 – 7.33 (m, *2*H), 7.23 – 7.27 (m, 2H), 6.71 (dd, *J* = 8.8, 2.2 Hz, 1H), 6.57 (d, *J* = 2.2 Hz, 1H), 4.61 – 4.73 (m, 2H), 4.58 (s, 2H), 4.14 – 4.18 (m, 1H), 3.57 (t, *J* = 6.2 Hz, 2 H), 3.41 (t, *J* = 5.7 Hz, 2H), 3.27 (t, *J* = 5.7 Hz,, 2H), 2.42 – 2.46 (m, 2H), UPLCMS: Mass calculated for C_22_H_23_FNO_7_, [M+H]^+^, 432. Found 432.

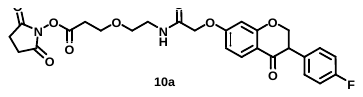

**Procedure for the Synthesis of 2,5-dioxopyrrolidin-1-yl 3-(2-(2-((3-(4-fluorophenyl)-4-oxochroman-7-yl)oxy)acetamido)ethoxy)propanoate (10a):** To a stirred solution of 3-(2-{2-[3-(4-Fluoro-phenyl)-4-oxo-chroman-7-yloxy]-acetylamino}-ethoxy)-propionic acid **(9a)** (0.46 mmol, 0.2 g) in DMF (2 mL) was added EDC (1.39 mmol, 0.26 g), N-hydroxy succinimide (0.69 mmol, 0.08 g) and allowed to stir at room temperature for 16 h. Then reaction mixture was partitioned between EtOAc (200 mL) and water (200 mL). Organic layer was separated, dried over sodium sulphate and concentrated under reduced pressure to provide 3-(2-{2-[3-(4-Fluoro-phenyl)-4-oxo-chroman-7-yloxy]-acetylamino}-ethoxy)-propionic acid 2,5-dioxo-pyrrolidin-1-yl ester (0.15 g, crude compound) as crude compound. This compound was used for the next step without further purification.

**Analytical data of 2,5-dioxopyrrolidin-1-yl 3-(2-(2-((3-(4-fluorophenyl)-4-oxochroman-7-yl)oxy)acetamido)ethoxy)propanoate (10a)**: UPLCMS: Mass calculated for C_26_H_26_FN_2_O_9_, [M+H]^+^, 529. Found 529.

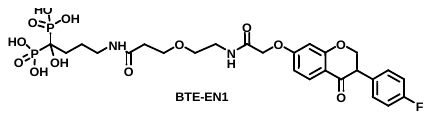

**Procedure for the Synthesis of {4-[3-(2-{2-[3-(4-Fluoro-phenyl)-4-oxo-chroman-7-yloxy]-acetylamino}-ethoxy)-propionylamino]-1-hydroxy-1-phosphono-butyl}-phosphonic acid (BTE-EN1):** To a stirred solution of 3-(2-{2-[3-(4-Fluoro-phenyl)-4-oxo-chroman-7-yloxy]-acetylamino}-ethoxy)-propionic acid 2,5-dioxo-pyrrolidin-1-yl ester **(10a)** (0.57 mmol, 0.3 g) in DMF (0.5 mL) was added aqueous (3 mL) solution of alendronic acid (0.96 mmol, 0.24 g) & KHCO_3_ (1.14 mmol, 0.11 g) and the resultant suspension was allowed to stir at room temperature for 16 h. Then reaction mixture was filtered through glass sintered and filtrate part was purified by RP column to afford {4-[3-(2-{2-[3-(4-Fluoro-phenyl)-4-oxo-chroman-7-yloxy]-acetylamino}-ethoxy)-propionylamino]-1-hydroxy-1-phosphono-butyl}-phosphonic acid (60 mg, 16%) as white solid. Analytical data of {4-[3-(2-{2-[3-(4-Fluoro-phenyl)-4-oxo-chroman-7-ylamino]-acetylamino}-ethoxy)-propionylamino]-1-hydroxy-1-phosphono-butyl}-phosphonic acid.

**Analytical data of {4-[3-(2-{2-[3-(4-Fluoro-phenyl)-4-oxo-chroman-7-yloxy]-acetylamino}-ethoxy)-propionylamino]-1-hydroxy-1-phosphono-butyl}-phosphonic acid** (**BTE-EN1):** ^1^H NMR (400 MHz, D_2_O) δ (ppm) 7.93 (d, *J* = 8.9 Hz, 1H), 7.39 – 7.31 (m, 2H), 7.19 (t, *J* = 8.8 Hz, 2H), 6.84 (dd, *J* = 9.0, 2.4 Hz, 1H), 6.65 (d, *J* = 2.4 Hz, 1H), 4.80 – 4.72 (m, 4H), 4.24 (t, *J* = 7.4 Hz, 1H), 3.72 (t, *J* = 6.2 Hz, 2H), 3.63 (t, *J* = 5.2 Hz, 2H), 3.51 (t, *J* = 5.3 Hz, 2H), 3.22 (t, *J* = 6.8 Hz, 2H), 2.46 (t, *J* = 6.2 Hz, 2H), 1.99 – 1.95 (m, 2H), 1.85 – 1.83 (m, 2H). UPLCMS: Mass calculated for C_26_H_34_FN_2_O_13_P_2_, [M+H]^+^, 663. Found 663.

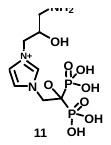

**Procedure for the Synthesis of 1-(3-Amino-2-hydroxy-propyl)-3-(2-hydroxy-2,2-bis-phosphono-ethyl)-3H-imidazol-1-ium (11):** To a stirred solution of zoledronic acid (19.38 mmol, 5 g) in MeOH (18 mL) & water (50 mL) was added Na_2_CO_3_ (21.38 mmol, 2.26 g), Oxiranylmethyl-carbamic acid tert-butyl ester (29.07 mmol, 5.03 g) and resultant reaction mixture was allowed to stir at 50 °C for 3 days. Reaction mixture was concentrated under reduced pressure. Crude residue thus obtained was allowed to stir with TFA for 4 h at RT. Then reaction mixture was concentrated and triturated with Et_2_O [200 mL] to afford 5 g of crude 1-(3-Amino-2-hydroxy-propyl)-3-(2-hydroxy-2,2-bis-phosphono-ethyl)-3H-imidazol-1-ium as white solid. This compound was used for the next step without further purification.

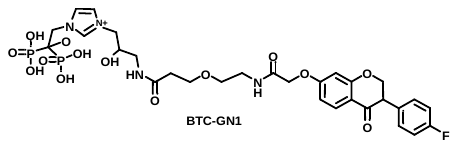

**Procedure for the Synthesis of 1-{3-[3-(2-{2-[3-(4-Fluoro-phenyl)-4-oxo-chroman-7-yloxy]-acetylamino}-ethoxy)-propionylamino]-2-hydroxy-propyl}-3-(2-hydroxy-2,2-bis-phosphono-ethyl)-3H-imidazol-1-ium (BTC-GN1):** To a stirred solution of 3-(2-{2-[3-(4-Fluoro-phenyl)-4-oxo-chroman-7-yloxy]-acetylamino}-ethoxy)-propionic acid 2,5-dioxo-pyrrolidin-1-yl ester **(10a)** (0.32 mmol, 0.17 g) in DMF (0.5 mL) was added aqueous (3 mL) solution of 1-(3-Amino-2-hydroxy-propyl)-3-(2-hydroxy-2,2-bis-phosphono-ethyl)-3H-imidazol-1-ium ( 0.32 mmol, 0.11 g) & KHCO_3_ (1.27 mmol, 0.13 g) then resultant suspension was allowed to stir at room temperature for 16 h. Reaction mixture was filtered through glass sintered and filtrate part was purified by RP column to afford 1-{3-[3-(2-{2-[3-(4-Fluoro-phenyl)-4-oxo-chroman-7-yloxy]-acetylamino}-ethoxy)-propionylamino]-2-hydroxy-propyl}-3-(2-hydroxy-2,2-bis-phosphono-ethyl)-3H-imidazol-1-ium (9 mg, 3.73%) as white solid. Analytical data of 1-{3-[3-(2-{2-[3-(4-Fluoro-phenyl)-4-oxo-chroman-7-yloxy]-acetylamino}-ethoxy)-propionylamino]-2-hydroxy-propyl}-3-(2-hydroxy-2,2-bis-phosphono-ethyl)-3H-imidazol-1-ium.

**Analytical data of 1-{3-[3-(2-{2-[3-(4-Fluoro-phenyl)-4-oxo-chroman-7-yloxy]-acetylamino}-ethoxy)-propionylamino]-2-hydroxy-propyl}-3-(2-hydroxy-2,2-bis-phosphono-ethyl)-3H-imidazol-1-ium (BTC-GN1):** ^1^H NMR (400 MHz, D_2_O) δ (ppm) 8.82 (s, 1H), 7.93 (d, *J* = 8.9 Hz, 1H), 7.58 (t, *J* = 1.8 Hz, 1H), 7.41 – 7.31 (m, 3H), 7.18 (t, *J* = 8.8 Hz, 2H), 6.84 (dd, *J* = 9.0, 2.5 Hz, 1H), 6.66 (d, *J* = 2.4 Hz, 1H), 4.72 – 4.70 (m, 2H), 4.22-4.07 (m, 4H), 3.73 (t, *J* = 6.1 Hz, 2H), 3.64 (t, *J* = 5.1 Hz, 2H), 3.50 (t, *J* = 5.2 Hz, 2H), 3.41 – 3.39 (m, 1H), 3.29 – 3.23 (m, 1H), 2.50 (t, *J* = 6.1 Hz, 2H). UPLCMS: Mass calculated for C_30_H_38_FN_4_O_14_P_2_^+^, [M]^+^, 759. Found 759.

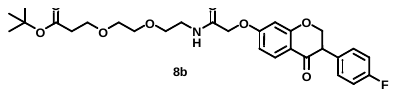

**Procedure for the Synthesis of 3-[2-(2-{2-[3-(4-Fluoro-phenyl)-4-oxo-chroman-7-yloxy]-acetylamino}-ethoxy)-ethoxy]-propionic acid tert-butyl ester (8b):** [3-(4-Fluoro-phenyl)-4-oxo-chroman-7-yloxy]-acetic acid **(7)** (3.16 mmol, 1 g), TBTU (4.74 mmol, 1.52 g) and DIPEA (9.49 mmol, 1.22 g) in DMF (5 mL) was allowed to stir at room temperature for 10 minutes. Followed by addition of PEG 2-Amine (3.16 mmol, 0.74g) and reaction mixture was allowed to stir at room temperature for 16 h. Then the reaction mixture was partitioned between EtOAc (300 mL) and water (200 mL). Organic layer was separated, dried over sodium sulphate and concentrated under reduced pressure. Crude residue thus obtained was purified by column chromatography using silica gel 100-200 mesh under gradient elution of 40-50 % EtOAc-hexane to afford 3-[2-(2-{2-[3-(4-Fluoro-phenyl)-4-oxo-chroman-7-yloxy]-acetylamino}-ethoxy)-ethoxy]-propionic acid tert-butyl ester (1.2 g, 71.34 %).

**Analytical data of 3-[2-(2-{2-[3-(4-Fluoro-phenyl)-4-oxo-chroman-7-yloxy]-acetylamino}-ethoxy)-ethoxy]-propionic acid tert-butyl ester (8b)**: ^1^H NMR (400 MHz, DMSO-*d*_6_) δ 8.14 (t, *J* = 5.5 Hz, 1H), 7.74 (d, *J* = 8.8 Hz, 1H), 7.29 – 7.33 (m, 2H), 7.15 (t, *J* = 8.8 Hz, 2H), 6.7 (dd, *J* = 8.8, 2.3 Hz, 1H), 6.57 (d, *J* = 2.2 Hz, 1H), 4.61 – 4.73 (m, 2H), 4.58 (s, 2H), 4.05 – 4.14 (m, 1H), 3.56 (t, *J* = 6.2 Hz, 2H) , 3.43 – 3.48 (m, 6H), 3.26 (t, *J* = 5.7 Hz, 2H), 2.39 – 2.49 (m, 2H), 1.38 (s, 9H) UPLCMS: Mass calculated for C_28_H_35_FNO_8_, [M+H]^+^, 532. Found 532.

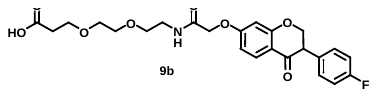

**Procedure for the Synthesis of 3-[2-(2-{2-[3-(4-Fluoro-phenyl)-4-oxo-chroman-7-yloxy]-acetylamino}-ethoxy)-ethoxy]-propionic acid (9b):** Solution of 3-(2-{2-[3-(4-Fluoro-phenyl)-4-oxo-chroman-7-yloxy]-acetylamino}-ethoxy)-propionic acid tert-butyl ester **(8b)** (2.26 mmol, 1.2 g) in DCM (10 mL) was treated with TFA (4 mL) and stirred at room temperature for 2 h. Then reaction mixture was concentrated under reduced pressure and triturated with pentane (50 mL) to afford 3-[2-(2-{2-[3-(4-Fluoro-phenyl)-4-oxo-chroman-7-yloxy]-acetylamino}-ethoxy)-ethoxy]-propionic acid (1.01 g, 94%) as off white solid.

**Analytical data of 3-[2-(2-{2-[3-(4-Fluoro-phenyl)-4-oxo-chroman-7-yloxy]-acetylamino}-ethoxy)-ethoxy]-propionic acid (9b):** ^1^H NMR (400 MHz, DMSO-*d*_6_) δ 12.1 (brs, 1H), 8.15 (t, *J* = 5.7 Hz, 1H), 7.74 (d, *J* = 8.8 Hz, 1H), 7.29 – 7.33 (m, *2*H), 7.23 – 7.27 (m, 2H), 6.71 (dd, *J* = 8.8, 2.1 Hz, 1H), 6.57 (d, *J* = 2.1 Hz, 1H), 4.61 – 4.73 (m, 2H), 4.59 (s, 2H), 4.14 – 4.18 (m, 1H), 3.43 – 3.49 (m, 6 H), 3.28 (t, *J* = 5.6 Hz, 2H), 3.28 (t, *J* = 5.6 Hz, 2H), 2.43 – 2.5 (m, 2H), UPLCMS: Mass calculated for C_24_H_27_FNO_8_, [M+H]^+^, 476. Found 476.

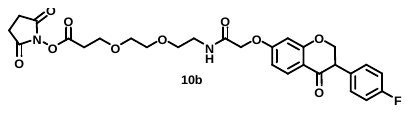

**Procedure for the Synthesis of 3-[2-(2-{2-[3-(4-Fluoro-phenyl)-4-oxo-chroman-7-yloxy]-acetylamino}-ethoxy)-ethoxy]-propionic acid 2,5-dioxo-pyrrolidin-1-yl ester (10b):** To a stirred solution of 3-[2-(2-{2-[3-(4-Fluoro-phenyl)-4-oxo-chroman-7-yloxy]-acetylamino}-ethoxy)-ethoxy]-propionic acid **(9b)** (0.42 mmol, 0.2 g) in DMF (2 mL) was added EDC (1.26 mmol, 0.24 g), N-hydroxy succinimide (0.63 mmol, 0.07 g) and allowed to stir at room temperature for 16 h. Then the reaction mixture was partitioned between EtOAc (100 mL) and water (200 mL). Organic layer was separated, dried over sodium sulphate and concentrated under reduced pressure to provide 3-[2-(2-{2-[3-(4-Fluoro-phenyl)-4-oxo-chroman-7-yloxy]-acetylamino}-ethoxy)-ethoxy]-propionic acid 2,5-dioxo-pyrrolidin-1-yl ester (0.15 g, crude). This compound was used for the next step without further purification.

**Analytical data of 3-[2-(2-{2-[3-(4-Fluoro-phenyl)-4-oxo-chroman-7-yloxy]-acetylamino}-ethoxy)-ethoxy]-propionic acid 2,5-dioxo-pyrrolidin-1-yl ester (10b):** UPLCMS: Mass calculated for C_28_H_30_FN_2_O_10_, [M+H]^+^, 573. Found 573.

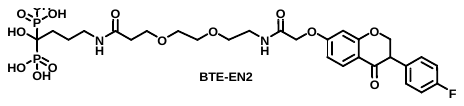

**Procedure for the Synthesis of (4-{3-[2-(2-{2-[3-(4-Fluoro-phenyl)-4-oxo-chroman-7-yloxy]-acetylamino}-ethoxy)-ethoxy]-propionylamino}-1-hydroxy-1-phosphono-butyl)-phosphonic acid (BTE-EN2):** To a stirred solution of 3-{2-[2-(2-{2-[3-(4-Fluoro-phenyl)-4-oxo-chroman-7-yloxy]acetylamino}-ethoxy)-ethoxy]-ethoxy}-propionic acid 2,5-dioxo-pyrrolidin-1-yl ester **(10b)** (0.35 mmol, 0.2 g) in DMF (0.5 mL) was added aqueous (3 mL) solution of alendronic acid (0.59 mmol, 0.15 g), KHCO_3_ (0.69 mmol, 0.07 g) and stirred at room temperature for 16 h. The heterogeneous reaction mixture was filtered through glass sintered and filtrate part was purified by RP column to afford (4-{3-[2-(2-{2-[3-(4-Fluoro-phenyl)-4-oxo-chroman-7-yloxy]-acetylamino}-ethoxy)-ethoxy]-propionylamino}-1-hydroxy-1-phosphono-butyl)-phosphonic acid (48 mg, 19%) as white solid. Analytical data of {4-[3(4-{3-[2-(2-{2-[3-(4-Fluoro-phenyl)-4-oxo-chroman-7-yloxy]-acetylamino}-ethoxy)-ethoxy]-propionylamino}-1-hydroxy-1-phosphono-butyl)-phosphonic acid.

**Analytical data of (4-{3-[2-(2-{2-[3-(4-Fluoro-phenyl)-4-oxo-chroman-7-yloxy]-acetylamino}-ethoxy)-ethoxy]-propionylamino}-1-hydroxy-1-phosphono-butyl)-phosphonic acid (BTE-EN2):** ^1^H NMR (400 MHz, D_2_O) δ (ppm) 7.90 (d, *J* = 8.8 Hz, 1H), 7.34 (t, *J* = 6.8 Hz, 2H), 7.18 (t, *J* = 8.6 Hz, 2H), 6.81 (d, *J* = 9.0 Hz, 1H), 6.63 (s, 1H), 4.76 – 4.72 (m, 4H), 4.21 (t, *J* = 7.4 Hz, 1H), 3.75 – 3.72 (m, 2H), 3.64 – 3.50 (m, 8H), 3.21 (t, *J* = 6.8 Hz, 2H), 2.50 (t, *J* = 6.2 Hz, 2H), 1.98 – 1.95 (m, 2H), 1.85 – 1.83 (m, 2H). UPLCMS: Mass calculated for C_28_H_38_FN_2_O_14_P_2_, [M+H]^+^, 707. Found 707.

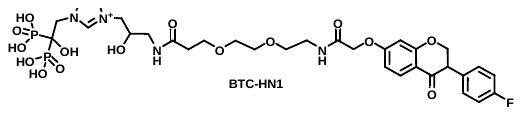

**Procedure for the Synthesis of 1-(3-{3-[2-(2-{2-[3-(4-Fluoro-phenyl)-4-oxo-chroman-7-yloxy]-acetylamino}-ethoxy)-ethoxy]-propionylamino}-2-hydroxy-propyl)-3-(2-hydroxy-2,2-bis-phosphono-ethyl)-3H-imidazol-1-ium (BTC-HN1):** To a stirred solution of 3-{2-[2-(2-{2-[3-(4-Fluoro-phenyl)-4-oxo-chroman-7-yloxy]acetylamino}-ethoxy)-ethoxy]-ethoxy}-propionic acid 2,5-dioxo-pyrrolidin-1-yl ester **(10b)** (0.37 mmol, 0.21 g) in DMF (0.7 mL) was added aqueous (3 mL) solution of 1-(3-Amino-2-hydroxy-propyl)-3-(2-hydroxy-2,2-bis-phosphono-ethyl)-3H-imidazol-1-ium (0.62 mmol, 0.21 g) & KHCO_3_ (1.47 mmol, 0.15 g), then resultant suspension was allowed to stir at room temperature for 16 h. Reaction mixture was filtered through glass sintered and filtrate part was purified by RP column to afford 1-(3-{3-[2-(2-{2-[3-(4-Fluoro-phenyl)-4-oxo-chroman-7-yloxy]-acetylamino}-ethoxy)-ethoxy]-propionylamino}-2-hydroxy-propyl)-3-(2-hydroxy-2,2-bis-phosphono-ethyl)-3H-imidazol-1-ium (17 mg, 5.8%) as white solid. Analytical data of 1-(3-{3-[2-(2-{2-[3-(4-Fluoro-phenyl)-4-oxo-chroman-7-yloxy]-acetylamino}-ethoxy)-ethoxy]-propionylamino}-2-hydroxy-propyl)-3-(2-hydroxy-2,2-bis-phosphono-ethyl)-3H-imidazol-1-ium.

**Analytical data of 1-(3-{3-[2-(2-{2-[3-(4-Fluoro-phenyl)-4-oxo-chroman-7-yloxy]-acetylamino}-ethoxy)-ethoxy]-propionylamino}-2-hydroxy-propyl)-3-(2-hydroxy-2,2-bis-phosphono-ethyl)-3H-imidazol-1-ium (BTC-HN1)**: ^1^H NMR (400 MHz, D_2_O) δ (ppm) 8.83 (s, 1H), 7.92 (d, *J* = 8.9 Hz, 1H), 7.58 (s, 1H), 7.43 (s, 1H), 7.35 (t, *J* = 7.0 Hz, 2H), 7.19 (t, *J* = 8.7 Hz, 2H), 6.82 (d, *J* = 9.0 Hz, 1H), 6.65 (s, 1H), 4.68 – 4.65 (m, 2H), 4.36 – 4.32 (m, 1H), 4.22 (t, *J* = 7.2 Hz, 1H), 4.13 – 4.10 (m, 2H), 3.76 (t, *J* = 6.0 Hz, 2H), 3.74 – 3.64 (m, 6H), 3.63 – 3.59 (m, 2H), 3.50 – 3.45 (m, 1H), 3.41 – 3.38 (m, 1H), 2.56 (t, *J* = 5.9 Hz, 2H). UPLCMS: Mass calculated for C_32_H_42_FN_4_O_15_P_2_^+^, [M]^+^, 803. Found 803.

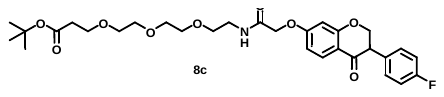

**Procedure for the Synthesis of 3-{2-[2-(2-{2-[3-(4-Fluoro-phenyl)-4-oxo-chroman-7-yloxy]-acetylamino}-ethoxy)-ethoxy]-ethoxy}-propionic acid tert-butyl ester (8c):** [3-(4-Fluoro-phenyl)-4-oxo-chroman-7-yloxy]-acetic acid (3.16 mmol, 1 g), TBTU (4.74 mmol, 1.52 g) and DIPEA (9.49 mmol, 1.22 g) in DMF (5 mL) was allowed to stir at room temperature for 10 minutes. Followed by addition of PEG 3-Amine (3.16 mmol, 0.87 g) and reaction was stirred at room temperature for 16 h. Then the reaction mixture was partitioned between EtOAc (300 mL) and water (200 mL). Organic layer was separated, dried over sodium sulphate and concentrated under reduced pressure. Crude residue thus obtained was purified by column chromatography using silica gel 100-200 mesh under gradient elution of 40-50 % EtOAc-hexane to afford 3-{2-[2-(2-{2-[3-(4-Fluoro-phenyl)-4-oxo-chroman-7-yloxy]-acetylamino}-ethoxy)-ethoxy]-ethoxy}-propionic acid tert-butyl ester (1.2 g, 75.88%).

**Analytical data of 3-{2-[2-(2-{2-[3-(4-Fluoro-phenyl)-4-oxo-chroman-7-yloxy]-acetylamino}-ethoxy)-ethoxy]-ethoxy}-propionic acid tert-butyl ester (8c):** ^1^H NMR (400 MHz, DMSO) δ 8.16 (t, *J* = 5.3 Hz, 1H), 7.74 (d, *J* = 8.8 Hz, 1H), 7.29 – 7.33 (m 2H), 7.15 (t, *J* = 8.8 Hz, 2H), 6.7 (dd, *J* = 8.8, 2 Hz, 1H), 6.57 (d, *J* = 2 Hz, 1H), 4.61 – 4.73 (m, 2H), 4.58 (s, 2H), 4.14 – 4.18 (m, 1H), 3.55 (t, *J* = 6.2 Hz, 2H) , 3.32 – 3.49 (m, 10H), 3.26 (t, *J* = 5.4 Hz, 2H), 2.38 (t, *J* = 6.2 Hz, 2H), 1.38 (s, 9H) UPLCMS: Mass calculated for C_30_H_39_FNO_9_, [M+H]^+^, 576. Found 576.

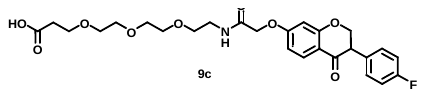

**Procedure for the Synthesis of 3-{2-[2-(2-{2-[3-(4-Fluoro-phenyl)-4-oxo-chroman-7-yloxy]-acetylamino}-ethoxy)-ethoxy]-ethoxy}-propionic acid (9c):** Solution of 3-(2-{2-[3-(4-Fluoro-phenyl)-4-oxo-chroman-7-yloxy]-acetylamino}-ethoxy)-propionic acid tert-butyl ester **(8c)** (1.2 g, 2.26 mmol) in DCM (15 mL) was treated with TFA (4 mL) and resultant reaction mixture was stirred at room temperature for 2 h. Then reaction mixture was concentrated under reduced pressure and triturated with pentane (50 mL) to afford 3-{2-[2-(2-{2-[3-(4-Fluoro-phenyl)-4-oxo-chroman-7-yloxy]-acetylamino}-ethoxy)-ethoxy]-ethoxy}-propionic acid (1.01 g, 94.03%) as off white solid.

**Analytical data of 3-{2-[2-(2-{2-[3-(4-Fluoro-phenyl)-4-oxo-chroman-7-yloxy]-acetylamino}-ethoxy)-ethoxy]-ethoxy}-propionic acid (9c)**: ^1^H NMR (400 MHz, DMSO-*d*_6_) δ 12.1 (brs, 1H), 8.16 (t, *J* = 5.5 Hz, 1H), 7.74 (d, *J* = 8.8 Hz, 1H), 7.29 – 7.33 (m, *2*H), 7.15 (t, *J* = 8.8 Hz, 2H), 6.71 (dd, *J* = 8.8, 2.1 Hz, 1H), 6.57 (d, *J* = 2.1 Hz, 1H), 4.61 – 4.73 (m, 2H), 4.59 (s, 2H), 4.14 – 4.18 (m, 1H), 3.56 (t, *J* = 6.4 Hz, 2H), 3.43 – 3.50 (m, 10H), 3.26 – 3.31 (m, 2H), 2.41 (t, *J* = 6.2 Hz, 2H), UPLCMS: Mass calculated for C_26_H_29_FNO_9_, [M-H]^+^, 518. Found 518.

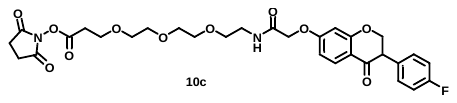

**Procedure for the Synthesis of 3-{2-[2-(2-{2-[3-(4-Fluoro-phenyl)-4-oxo-chroman-7-yloxy]-acetylamino}-ethoxy)-ethoxy]-ethoxy}-propionic acid 2,5-dioxo-pyrrolidin-1-yl ester (10c):** To a stirred solution of 3-{2-[2-(2-{2-[3-(4-Fluoro-phenyl)-4-oxo-chroman-7-yloxy]-acetylamino}-ethoxy)-ethoxy]-ethoxy}-propionic acid **(9c)** (0.42 mmol, 0.2 g) in DMF (2 mL) was added EDC (1.26 mmol, 0.24 g), N-hydroxy succinimide (0.63 mmol, 0.07 g) and allowed to stir at room temperature for 16 h. Then the reaction mixture was partitioned between EtOAc (100 mL) and water (200 mL). Organic layer was separated, dried over sodium sulphate and concentrated under reduced pressure to provide 3-{2-[2-(2-{2-[3-(4-Fluoro-phenyl)-4-oxo-chroman-7-yloxy]-acetylamino}-ethoxy)-ethoxy]-ethoxy}-propionic acid 2,5-dioxo-pyrrolidin-1-yl ester (0.24 g, crude compound). **Analytical data of 3-{2-[2-(2-{2-[3-(4-Fluoro-phenyl)-4-oxo-chroman-7-yloxy]acetylamino}-ethoxy)-ethoxy]-ethoxy}-propionic acid 2,5-dioxo-pyrrolidin-1-yl ester (10c)**: UPLCMS: Mass calculated for C_30_H_34_FN_2_O_11_, [M+H]^+^ 617. Found 617.

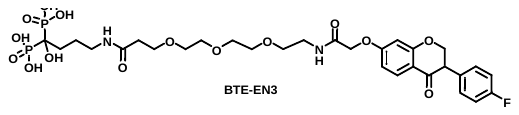

**Procedure for the Synthesis of [4-(3-{2-[2-(2-{2-[3-(4-Fluoro-phenyl)-4-oxo-chroman-7-yloxy]-acetylamino}-ethoxy)-ethoxy]-ethoxy}-propionylamino)-1-hydroxy-1-phosphono-butyl]-phosphonic acid (BTE-EN3):** To a stirred solution of 3-{2-[2-(2-{2-[3-(4-Fluoro-phenyl)-4-oxo-chroman-7-yloxy]acetylamino}-ethoxy)-ethoxy]-ethoxy}-propionic acid 2,5-dioxo-pyrrolidin-1-yl ester **(10c)** (0.44 mmol, 0.11 g) in DMF (0.5 mL) was added aqueous (3 mL) solution of alendronic acid (0.44 mmol, 0.27 g) & KHCO_3_ (0.88 mmol, 0.09 g), then resultant suspension was allowed to stir at room temperature for 16 h. Reaction mixture was filtered through glass sintered and filtrate part was purified by RP column to afford [4-(3-{2-[2-(2-{2-[3-(4-Fluoro-phenyl)-4-oxo-chroman-7-yloxy]-acetylamino}-ethoxy)-ethoxy]-ethoxy}-propionylamino)-1-hydroxy-1-phosphono-butyl]-phosphonic acid (62 mg, 19%) as white solid. Analytical data of [4-(3-{2-[2-(2-{2-[3-(4-Fluoro-phenyl)-4-oxo-chroman-7-yloxy]-acetylamino}-ethoxy)-ethoxy]-ethoxy}-propionylamino)-1-hydroxy-1-phosphono-butyl]-phosphonic acid.

**Analytical data of [4-(3-{2-[2-(2-{2-[3-(4-Fluoro-phenyl)-4-oxo-chroman-7-yloxy]-acetylamino}-ethoxy)-ethoxy]-ethoxy}-propionylamino)-1-hydroxy-1-phosphono-butyl]-phosphonic acid (BTE-EN3):** ^1^H NMR (400 MHz, D_2_O) δ (ppm) 7.78 (d, *J* = 8.8 Hz, 1H), 7.22 (t, *J* = 7.0 Hz, 2H), 7.05 (t, *J* = 8.8 Hz, 2H), 6.69 (d, *J* = 9.7 Hz, 1H), 6.52 (d, *J* = 2.5 Hz, 1H), 4.63 – 4.61 (m, 4H), 3.62 (t, *J* = 6.5 Hz, 2H), 3.53 – 3.48 (m, 9H), 3.39 (d, *J* = 5.3 Hz, 2H), 3.09 (t, *J* = 6.7 Hz, 2H), 2.38 (t, *J* = 6.4 Hz, 2H), 1.82 – 1.78 (m, 2H), 1.69 – 1.68 (m, 2H), UPLCMS: Mass calculated for C_30_H_42_FN_2_O_15_P_2_, [M+H]^+^, 751. Found 751.

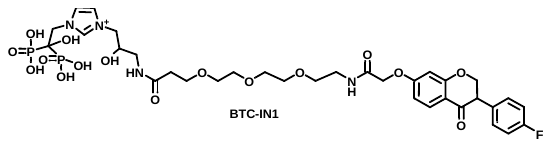

**Procedure for the Synthesis of 1-[3-(3-{2-[2-(2-{2-[3-(4-Fluoro-phenyl)-4-oxo-chroman-7-yloxy]-acetylamino}-ethoxy)-ethoxy]-ethoxy}-propionylamino)-2-hydroxy-propyl]-3-(2-hydroxy-2,2-bis-phosphono-ethyl)-3H-imidazol-1-ium (BTC-IN1):** To a stirred solution of 3-{2-[2-(2-{2-[3-(4-Fluoro-phenyl)-4-oxo-chroman-7-yloxy]acetylamino}-ethoxy)-ethoxy]-ethoxy}-propionic acid 2,5-dioxo-pyrrolidin-1-yl ester **(10c)** (0.32 mmol, 0.2 g) in DMF (1 mL) was added aqueous (3 mL) solution of 1-(3-Amino-2-hydroxy-propyl)-3-(2-hydroxy-2,2-bis-phosphono-ethyl)-3H-imidazol-1-ium (0.55 mmol, 0.19 g), KHCO_3_ (0.97 mmol, 0.097 g) and the resultant suspension was allowed to stir at room temperature for 16 h. Then reaction mixture was filtered through glass sintered and filtrate part was purified by RP column to afford 1-[3-(3-{2-[2-(2-{2-[3-(4-Fluoro-phenyl)-4-oxo-chroman-7-yloxy]-acetylamino}-ethoxy)-ethoxy]-ethoxy}-propionylamino)-2-hydroxy-propyl]-3-(2-hydroxy-2,2-bis-phosphono-ethyl)-3H-imidazol-1-ium (5 mg, 1.8%) as white solid. Analytical data of 1-[3-(3-{2-[2-(2-{2-[3-(4-Fluoro-phenyl)-4-oxo-chroman-7-yloxy]-acetylamino}-ethoxy)-ethoxy]-ethoxy}-propionylamino)-2-hydroxy-propyl]-3-(2-hydroxy-2,2-bis-phosphono-ethyl)-3H-imidazol-1-ium.

**Analytical data of 1-[3-(3-{2-[2-(2-{2-[3-(4-Fluoro-phenyl)-4-oxo-chroman-7-yloxy]-acetylamino}-ethoxy)-ethoxy]-ethoxy}-propionylamino)-2-hydroxy-propyl]-3-(2-hydroxy-2,2-bis-phosphono-ethyl)-3H-imidazol-1-ium (BTC-IN1):** ^1^H NMR (400 MHz, D_2_O) δ (ppm) 8.84 (s, 1H), 7.92 (d, *J* = 8.8 Hz, 1H), 7.58 (s, 1H), 7.44 (s, 1H), 7.35 (t, *J* = 7.1 Hz, 2H), 7.19 (t, *J* = 8.8 Hz, 2H), 6.83 (d, *J* = 9.0 Hz, 1H), 6.66 (d, *J* = 2.4 Hz, 1H), 4.81 – 4.74 (m, 6H), 4.72 – 4.69 (m, 2H), 4.38 (d, *J* = 11.2 Hz, 1H), 4.26 – 4.14 (m, 1H), 4.13 – 4.11 (m, 2H), 3.76 (t, *J* = 6.1 Hz, 2H), 3.69 – 3.58 (m, 8H), 3.53 (d, *J* = 5.3 Hz, 2H), 3.46 – 3.42 (m, 1H), 3.35 – 3.30 (m, 1H), 2.55 (t, *J* = 6.0 Hz, 2H). UPLCMS: Mass calculated for C_34_H_46_FN_4_O_16_P_2_^+^, [M]^+^, 847 Found 847.

***Purification of DABDs***

All six final compounds (BTE-EN1, BTE-EN2, BTE-EN3, BTC-GN1, BTE-HN1, and BTC-IN1) were purified by reverse phase high-performance liquid chromatography (RP-HPLC) using a C18 column with water-acetonitrile and ammonium bicarbonate buffer, yielding ammonium salts that are hygroscopic and readily water soluble.

**Animal models**

***Intracardiac (IC) injection pre-treatment model***

Beginning 3 days prior to cell injection, mice were treated with weekly intraperitoneal (IP) injections of vehicle (control), BTE-EN1, BTC-HN1 or BTC-IN1, with each compound given at 10, 100 and 1000 µg/kg. PC3-Luc cells (4 x 10^5^) were inoculated into the left ventricle of 6-8 weeks male athymic nude mice (Charles River) via IC injection under ultrasound guidance (VEVO 3100) as previously described (*1, 3*). In Vivo Imaging System (IVIS) imaging was performed 30 min after IC injection to confirm systemic distribution of cells. Mice were monitored hourly and then daily for toxicity, underwent body weight measurement and IVIS imaging weekly for 4 weeks. Animal survival was monitored for 4 weeks and then euthanized. After termination of the experiment, mandibular bone destruction was quantified by microCT imaging (Inveon, siemens) as previously described (*1*).

***Caudal artery (CA) injection model of established bone metastasis***

PC3-Luc cells (1.5 x 10^5^) were suspended in 100 uL PBS and were injected into the caudal artery of 8-9 weeks male athymic nude mice (Charles River) to establish metastasis in the femur and tibia as described previously (*4*). IVIS imaging was performed at 30 minutes post cell injection on 50 mice, confirming regional distribution of cells in the lower body of 28 mice. Body weight was measured weekly. After 2 weeks post-injection of cells IVIS imaging was performed, yielding 19 mice demonstrating signal localized to the lower limbs. Mice were randomized into 2 groups based on the IVIS signal, giving N=9 mice in the control group and N=10 in the treatment group. The control group and treatment group received weekly IP injection of PBS (vehicle) and 1000 µg/kg BTE-EN1 for 6 weeks, respectively. Mice were sacrificed at 8 weeks post-injection based on weight loss in controls and underwent microCT imaging using Quantum GX2 (PerkinElmer).

To determine the leg bone destruction, the low body region of each mouse was scanned using the following parameters: 70 kVp, 114 uA, 14 min exposure time, 0.5 mm aluminum filter, 36 µm field of view (FOV) and 72 µm voxel size. The region of interest (ROI) of leg bone was defined as the whole femur and the epiphysis of the tibia down to the fibula insertion point. Image segmentation was performed using the “Segment” tool. Segmentation masks of the defined ROI were generated with a combination of semi-automatic (Threshold Volume) and manual (Manual Trace) techniques and saved as object maps. The bone destruction area in the leg was calculated using “Measure” tool after loading the saved object map.

***Intracardiac (IC) injection model of established bone metastasis***

PC3-Luc cells (4 x 10^5^) were injected into the left ventricle of 8-10 weeks male athymic nude mice (Charles River) via IC injection with ultrasound guidance (VEVO 3100) (*1, 3*). IVIS imaging was performed 30 min after IC injection. Systemic distribution of cells was confirmed in 56 out of 60 injected mice. Five days post cell injection, mice were randomized into two groups (control, N=28; treatment, N=28), and received weekly IP injection of PBS (vehicle) and 1000 µg/kg BTE-EN1 for 6 weeks, respectively. Body weight and IVIS imaging were monitored weekly. With time 5 mice in the control group and 3 mice in the treatment group demonstrated concentration and increase of IVIS signal over the heart, indicative of leakage of cells into the pericardial space during injection. Thus, these mice were excluded. At the completion of the study, microCT imaging was performed on all remaining animals using the Quantum GX2 microCT imaging system (PerkinElmer).

Whole body CT images were acquired at 70 kVp, 114 uA, 14 min exposure time, with 0.5 mm aluminum filter, 72 mm field of view (FOV), and 144 µm voxel size. IC injections of PC3-Luc cells produce metastases to both mandible and leg (femur and tibia). To determine the bone destruction in mandible and leg, sub volume reconstructions were carried out for head (FOV=23.04 mm) and low body (FOV=36.864 mm) to achieve the resolution of 45 µm and 72 µm, respectively, using Quantum GX2 microCT software. The microCT data were then imported into Analyze 14.0 software (AnalyzeDirect, Inc.) for quantitative analysis of bone destruction. The ROI of leg bone was defined as the whole femur and the epiphysis of the tibia down to the fibula insertion point. The skull and leg image segmentations were performed using the “Segment” tool. Segmentation masks of the defined ROI were generated with a combination of semi-automatic (Threshold Volume) and manual (Manual Trace) techniques and saved as object maps. The mandibular bone destruction was calculated by measuring ratio of the bone destruction area to the whole mandible area using “Measure” tool. The leg bone destruction area was measured and calculated using “Measure” tool.

***In vivo* toxicity evaluation**

The toxicity of BTE-EN1 was evaluated by Calvert Laboratories, Inc. Specifically, cohorts of N=6 male CD-1 mice were administered vehicle control or 10 mg BTE-EN1/kg by IP injection on Days 1, 3, 5, and 7. On the days of dosing, animals were observed prior to each dose administration, immediately post-dose, and approximately 1-2 hours post-dose. On non-dosing days, animals were observed once daily. Animals were observed once prior to scheduled sacrifice on Day 8. Body weights were recorded at the time of randomization, prior to dose administration on Day 1, and on Day 7. A fasting terminal body weight was recorded prior to sacrifice on Day 8. Within each cohort, terminal blood was collected from 3 mice for evaluation of hematology parameters and from a separate 3 mice for evaluation of clinical chemistry parameters. Selected tissues were harvested at necropsy, selected organs weighed, and selected tissues from all mice were evaluated microscopically.

**Supplementary Movie 1. Intracardiac injection with ultrasound guidance**. PC3-Luc cells were inoculated in the left ventricle of nude mice via intracardiac injection under ultrasound guidance.

**Supplementary Data file 1. ^1^H NMR spectra and HPLC trace of representative compounds BTE-EN1, BTE-EN2, BTE-EN3, BTC-GN1, BTC-HN1, and BTE-IN1.**
