## Supplementary Data file 1 for "A New Class of Precision Therapeutics that Inhibit Prostate Cancer Mediated Bone Destruction"

**^1^H NMR spectra and HPLC trace of representative compounds BTE-EN1, BTE-EN2, BTE-EN3, BTC-GN1, BTC-HN1, and BTE-IN1**

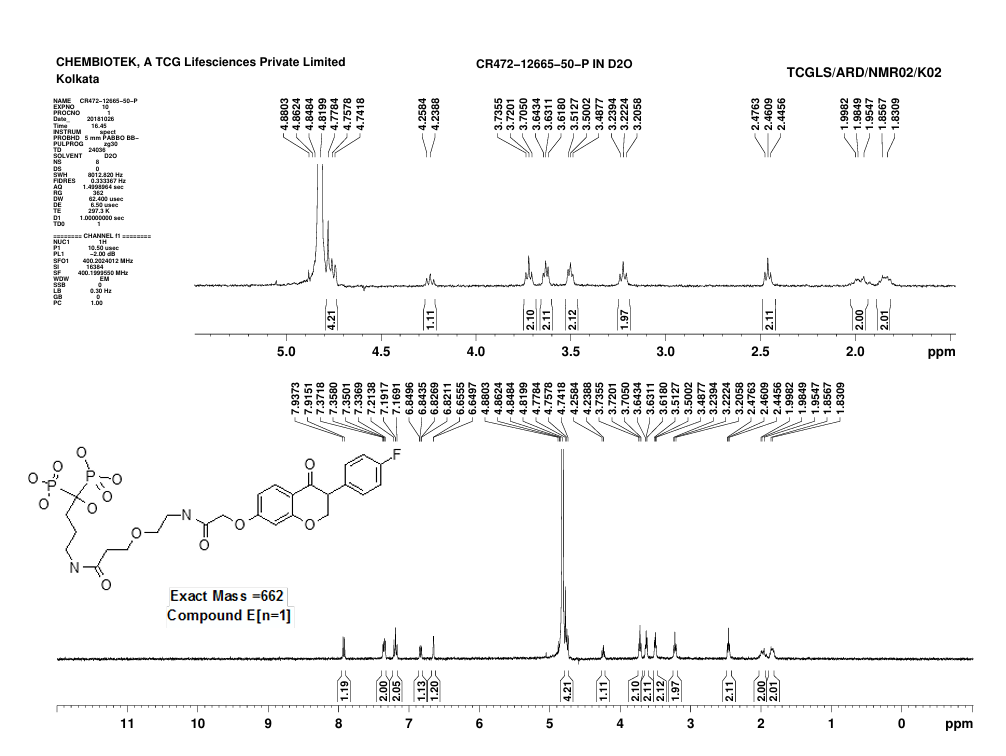

**^1^H NMR spectra of compound BTE-EN1**

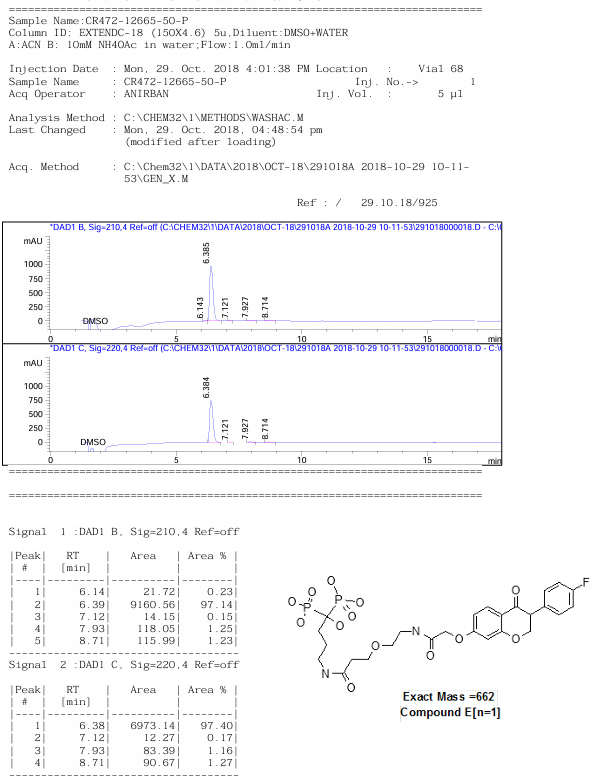

**HPLC trace of compound BTE-EN1**

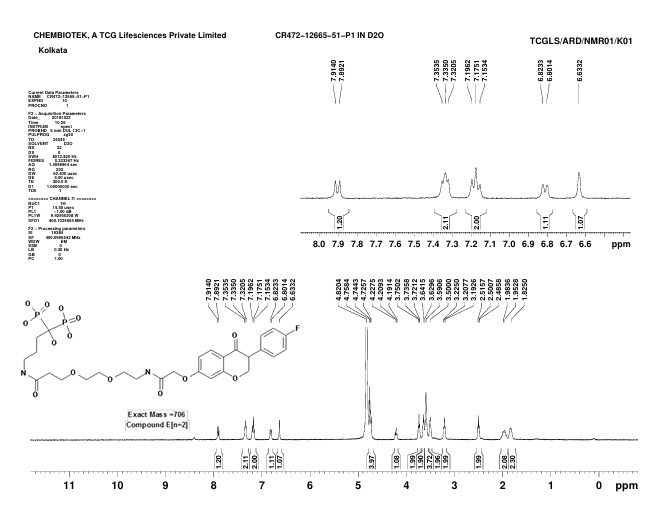

**^1^H NMR spectra of compound BTE-EN2**

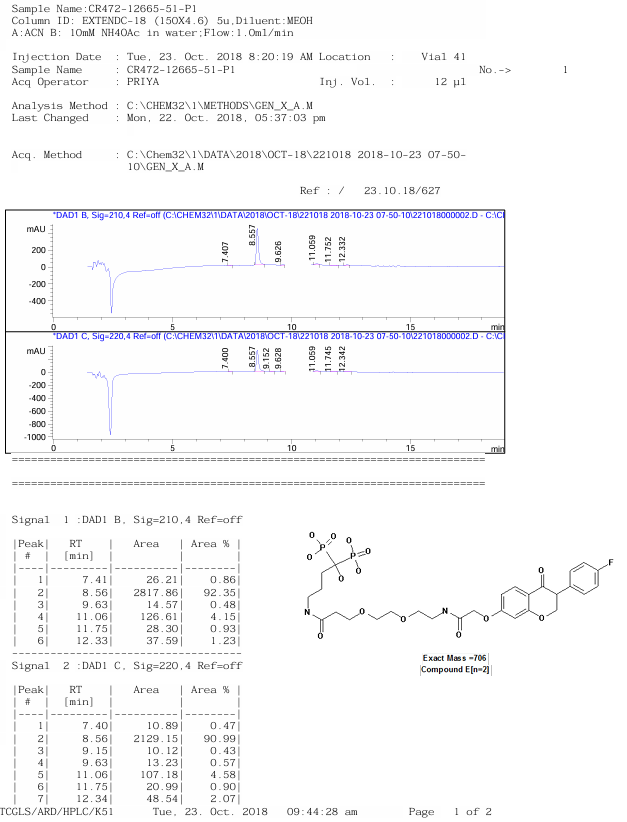

**HPLC trace of compound BTE-EN2**

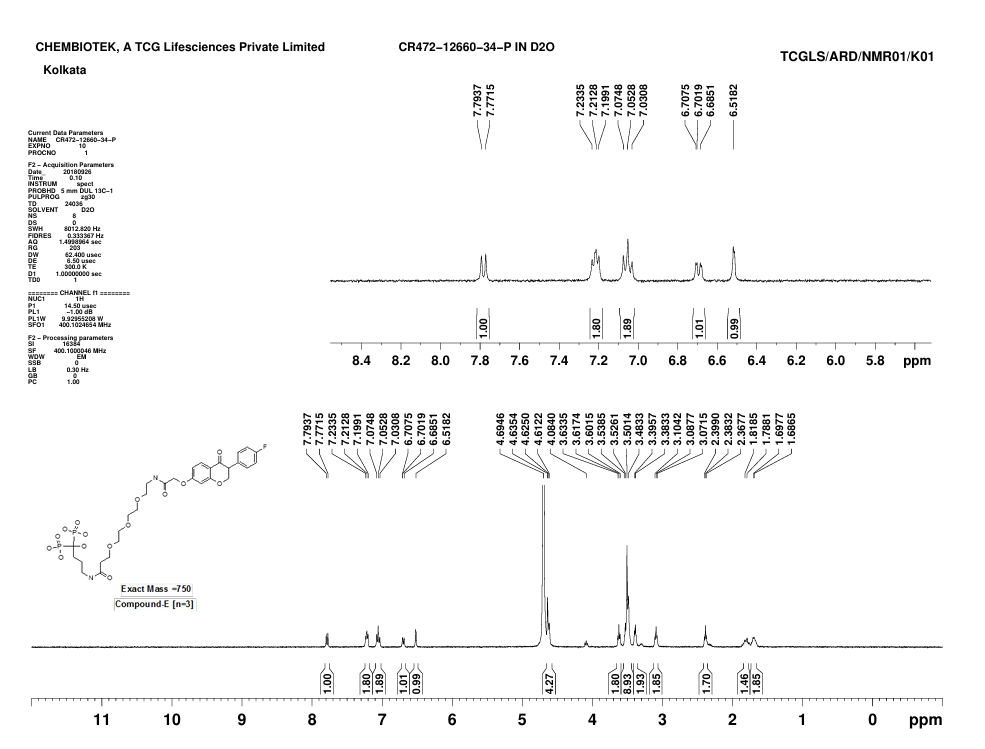

**^1^H NMR spectra of compound BTE-EN3**

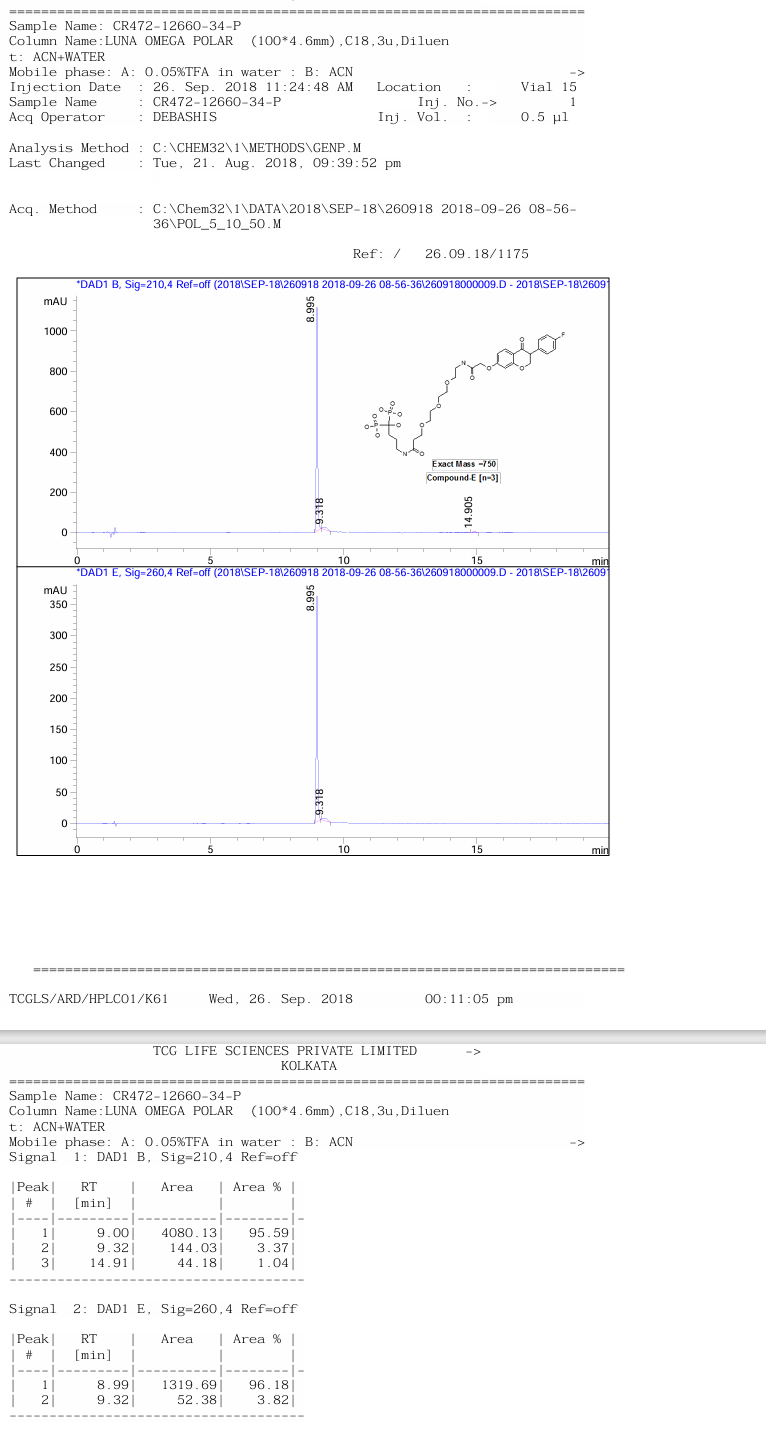

**HPLC trace of compound BTE-EN3**

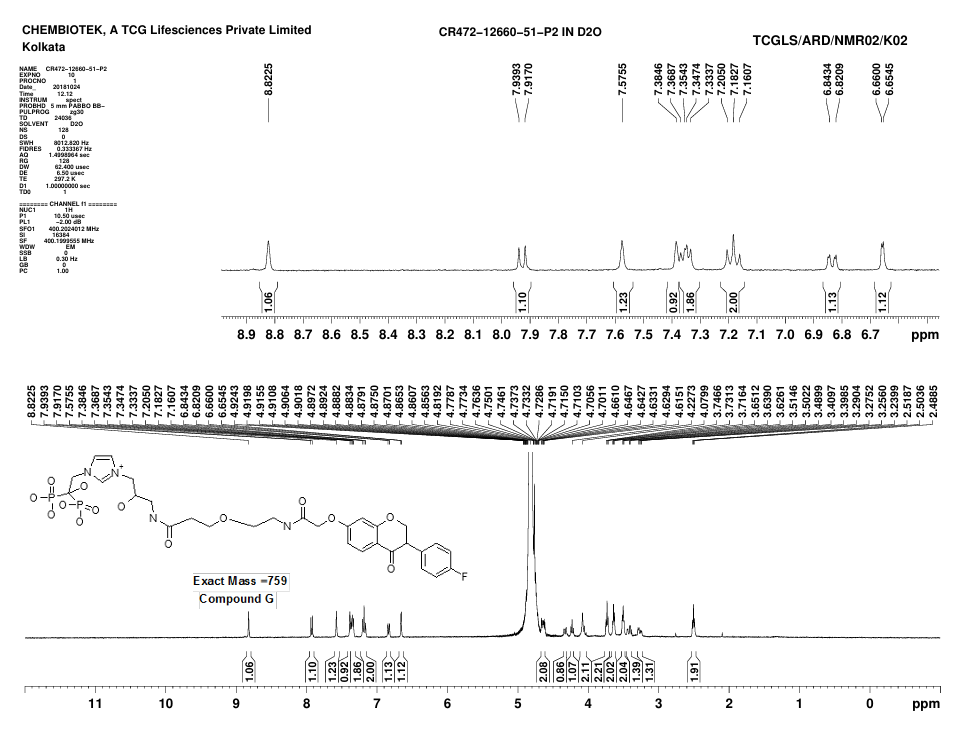

**^1^H NMR spectra of compound BTC-GN1**

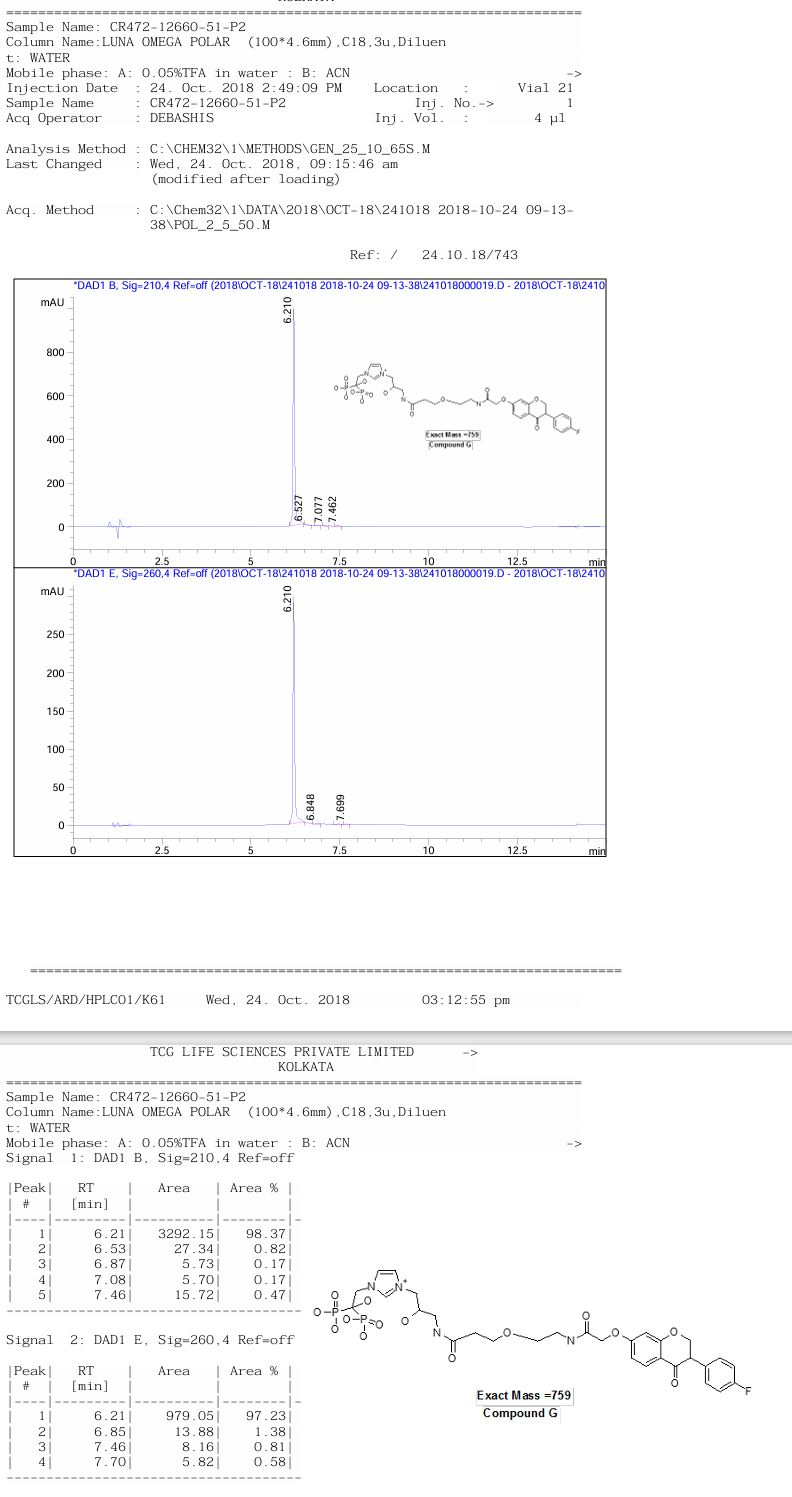

**HPLC trace of compound BTC-GN1**

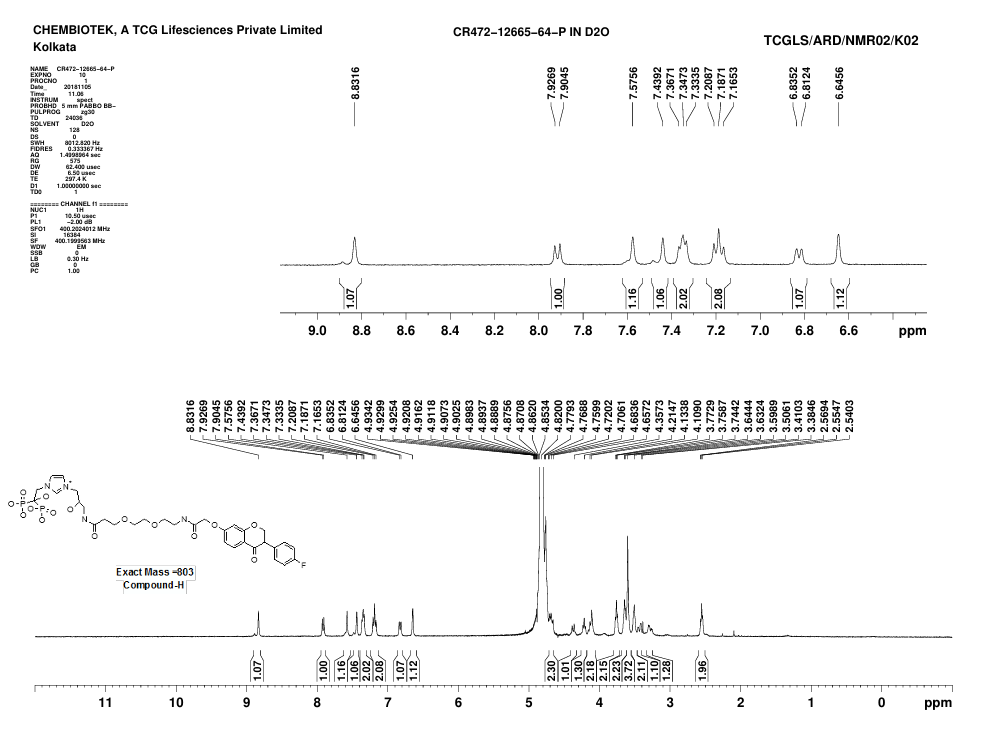

**^1^H NMR spectra of compound BTC-HN1**

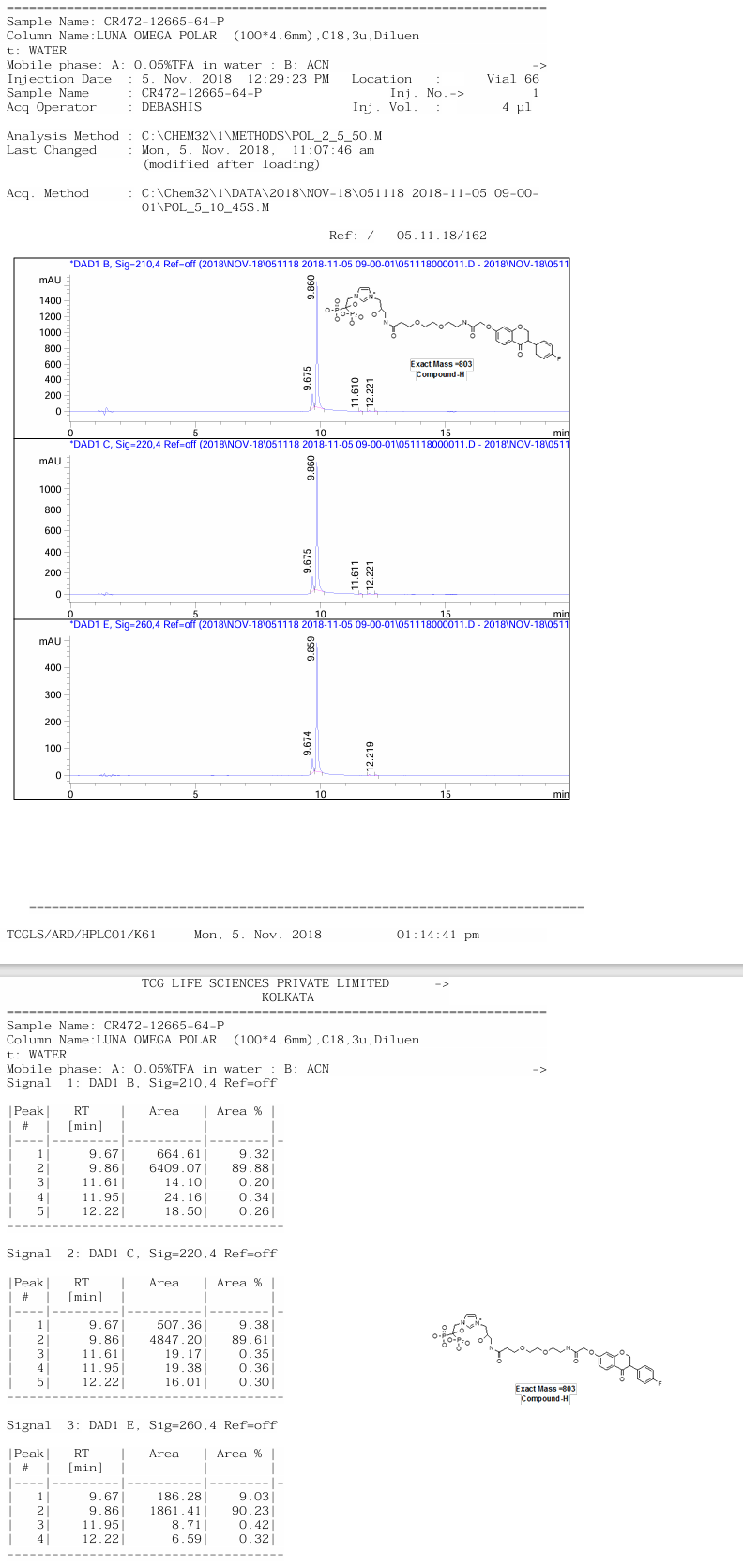

**HPLC trace of compound BTC-HN1**

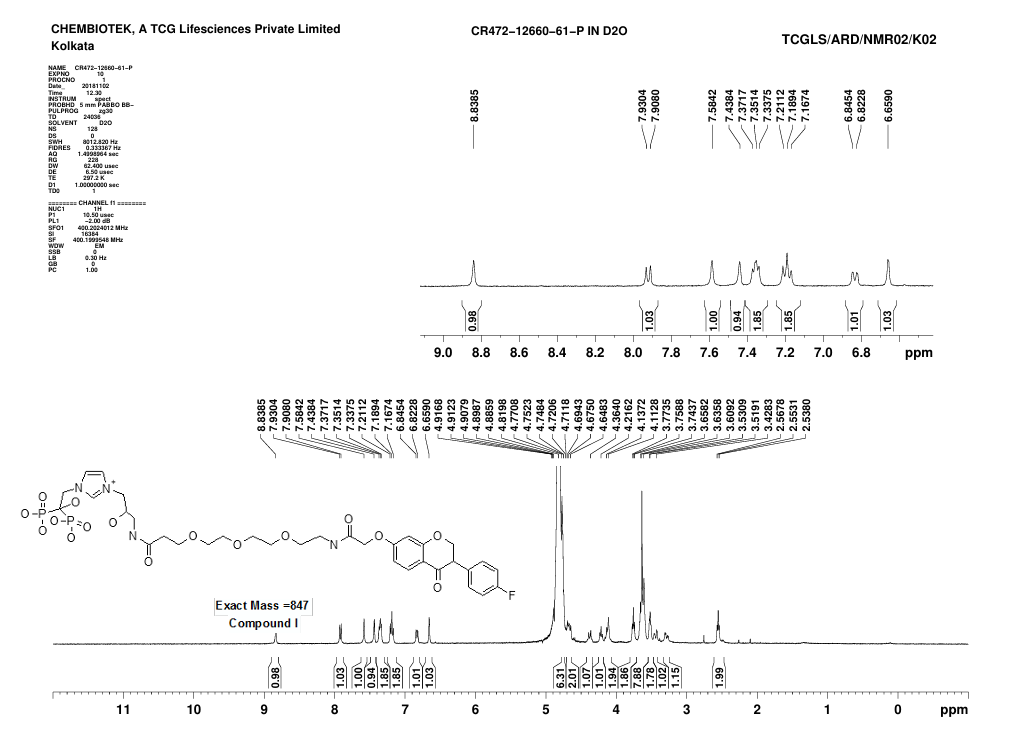

**^1^H NMR spectra of compound BTC-IN1**

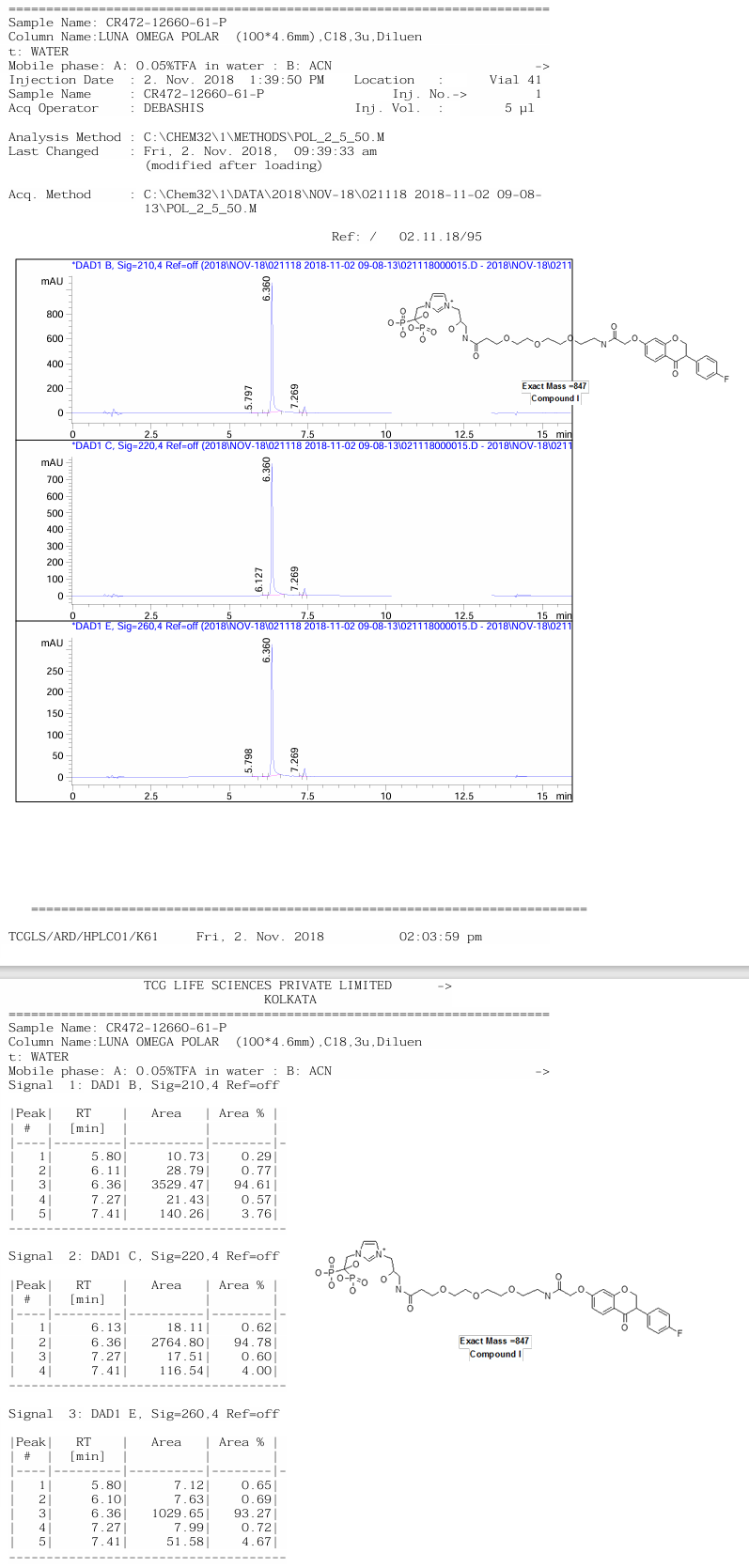

**HPLC trace of compound BTC-IN1**
